# The geometric properties of the molecular surfaces of protein crystal structures and AlphaFold predicted models

**DOI:** 10.1101/2024.11.17.624000

**Authors:** Lincong Wang

## Abstract

Previous analyses of protein structures have focused primarily on three-dimensional folds, their secondary structures, and binding or active sites while their molecular surfaces have received much less attention, due possibly to the lack of accurate and robust programs for their computation.

Using SESA we have analyzed the molecular surfaces of three mutually exclusive sets, **G, S** and **M**, of protein crystal structures. **G** and **S** include only non-membrane proteins with the latter having only monomers while **M** has only membrane proteins. The analyses show that SAS area per atom *µ*_*s*_ decreases while probe area per atom *µ*_*p*_ increases with the number of atoms in a molecule *n*. Most interestingly, the fitted power laws for *µ*_*s*_ intersect with those for *µ*_*p*_ at *n* = 957 for **G**, *n* = 875 for **S** and *n* = 1, 061 for **M**. They correspond respectively to 60, 57 and 64 amino acid residues. The power laws and their intersections provide an explanation for protein structural integrity and stability in general and the transition in particular from peptides typically with random conformations in solution to proteins usually with a dominant conformation.

We have also analyzed the molecular surfaces of the AlphaFold models for twenty seven proteomes. The analyses show that the molecular surfaces for thirteen prokaryotic proteomes resemble those for the crystal structures while those for fourteen eukaryotic ones differ largely from both of them. The variation may have significant implication in theory in that there exist genuine differences between prokaryotic and eukaryotic proteomes, and in application in that the current AlphaFold models for eukaryotic proteomes are likely not adequate for structure-based drug design in particular.

**Significance statement:** *A newly-developed analytic and robust program*, SESA, *has been applied to three mutually exclusive sets*, **G, S** *and* **M**, *of protein crystal structures and the AlphaFold models for twenty seven proteomes to compute their exterior solvent-excluded surface (SES) areas. The results show that for the crystal structures the areas per atom for SAS µ*_*s*_, *probe µ*_*p*_ *and toroidal µ*_*t*_ *patches each follows a power law with n, the number of atoms in a structure or model. Specifically, µ*_*s*_ *decreases while µ*_*p*_ *increases with n. Most interestingly, the power laws for µ*_*s*_ *intersect with those for µ*_*p*_ *at n* = 957 *for* **G**, *n* = 875 *for* **S** *and n* = 1, 061 *for* **M**. *They correspond respectively to* 60, 57 *and* 64 *residues. A SAS patch is convex while a probe one concave, thus a power law for µ*_*s*_ *intersects with that for µ*_*p*_ *when the total area of the patches with a negative curvature equals that with a positive curvature if one ignores toroidal patches. The points of intersection for* **G** *and* **S** *are close to the number of residues required for a polypeptide to adopt a dominant conformation in solution, and thus provide an explanation for why a chain with <* 50 *residues, that is, a peptide, has in general only random conformations in solution. In addition, the SESs of the AlphaFold models for thirteen prokaryotic proteomes resemble those for the crystal structures. However, in stark contrast with the crystal structures and the models for prokaryotic proteomes, the SESs for fourteen eukaryotic proteomes differ largely from both of them. The differences likely have significant implications for structural biology and the applications of AlphaFold models*.

## 1 Introduction

The experimentally-derived structures of proteins have been analyzed in depth to rationalize their physical properties such as solvation and folding, and their biochemical properties such as binding affinity [1] and catalytic efficiency [2]. Previous analyses though extensive focus largely on three-dimensional (3D)^1^ folds, secondary structure content, and the physical-chemical and geometric properties of ligand-protein interface. By contrast, protein surface has not been as well investigated. For example, the contributions of surface to the solvation, folding and structural stability of proteins have not been accurately quantified [3, 4, 5, 6, 7, 8, 9].

There exist three main models for the surface of a molecule called, respectively, van der Waals (VDW) surface, solvent accessible surface (SAS) [10, 11] and molecular surface (also called solvent-excluded surface (SES)^2^) [11, 12]. A SES is a two-dimensional (2D) manifold impenetrable to solvent molecules, and is composed of three types of 2D patches: convex spherical polygon on a solvent-accessible atom, saddle-shaped toroidal patch determined by two accessible atoms, and concave spherical polygon on a probe by three accessible atoms. Their surface areas are called, respectively, SAS area, toroidal area and probe area.

To evaluate SES’s contributions to protein solvation, folding and structural stability, and to obtain clue to the adaptation of native proteins to their environments, we have applied our robust and analytic SES area computation program SESA [13] to three mutually-exclusive sets of crystal structures, **G, S** and **M. G** and **S** include only non-membrane proteins with the former having both monomers and multimers and the latter only monomers, while **M** includes only membrane proteins. Importantly, the structures in **G** have no missing internal residues^3^ and a very low percentage of missing atoms per structure. Out of several SES geometric properties examined we found that SAS, probe and toroidal areas per atom, *µ*_*s*_, *µ*_*p*_ and *µ*_*t*_, each changes with the number of atoms in a structure *n* by following a power law. Specifically, *µ*_*s*_ decreases while *µ*_*p*_ increases with *n*. Most interestingly, the power laws for *µ*_*s*_ intersect with those for *µ*_*p*_ at *n* = 957 for **G**, *n* = 875 for **S** and *n* = 1, 061 for **M**. They correspond, respectively, to 60, 57 and 64 residues. An intersection occurs when *µ*_*s*_(*n*) = *µ*_*p*_(*n*), i.e. when the total SAS area of a molecule equals its total probe area. A SAS patch is convex while a probe one concave, thus a power law for *µ*_*s*_ intersects with that for *µ*_*p*_ when the total area of the patches with a negative curvature equals that with a positive curvature (if one ignores the toroidal patches, each of which includes regions of positive and negative curvatures). The points of intersection for **G** and **S** are close to the number of residues required for a polypeptide to have a dominant conformation in solution. The variations of *µ*_*s*_, *µ*_*p*_ and *µ*_*t*_ with molecular size (*n*) together with the power laws and their intersections provide an explanation for why a polypeptide of *<* 50 residues, that is, a peptide, has in general only random conformations in solution. In other words, the power laws and their intersections may explain why the transition from the broad conformational distribution of a typical peptide to the narrow one of a regular protein occurs between 50−60 residues. Furthermore, we hypothesize that the ratio 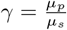 is closely related to protein stability and folding in the sense that *γ >* 1 is a necessary condition for the folding of a typical water-soluble protein.

The neural network-based program, AlphaFold [14, 15, 16, 17], represents a leap-forward in protein structure prediction in terms of structural coverage, and accuracy as judged by the RMSDs for both backbone and all atoms, global distance test (GDT), TM-score, electron density map [18], and other criteria [19, 20, 21, 22, 23, 24]. However, to our best knowledge neither the SASs nor the SESs of the predicted models have been analyzed. Our analyses of the SESs of the 350, 974 AlphaFold models for thirteen prokaryotic and fourteen eukaryotic proteomes show that the SESs for the former resemble those for crystal structures while the SESs for the latter differ largely from them. Specifically, the *µ*s^4^ of the models for the eukaryotic proteomes are broadly distributed, hardly change with *n* and only minimally follow a power law. The differences may have significant implication in theory in that there exist genuine differences between prokaryotic and eukaryotic proteomes, and in application in that the current AlphaFold models for eukaryotic proteomes are likely not adequate for structure-based drug design in particular.

## 2 Materials and Methods

In this section we first describe the three sets of crystal structures and the twenty seven AlphaFold proteomes, and then briefly present the computation of SES area and the definitions of SES patch areas per surface atom (*µ*s). Finally, we describe the fittings to power law of the changes with *n* of *µ*s, and to linear equation of the change with *n* of the number of surface atom *n*_*s*_ of a molecule.

### 2.1 Data sets and the preprocessing of the PDB files for SES area computation

From the PDB we downloaded three mutually exclusive sets of protein crystal structures, **S, G** and **M. S** has 4, 147 structures of monomeric proteins with a resolution range of 0.64 − 3.50Å and mean of 1.68Å, a *R*_free_ range of 0.05− 0.350 and mean of 0.207. The percentage of missing atoms^5^ ranges from 0.0 − 2.0% with a mean of 0.299% and a median of 0.051%. **G** has 8, 616 structures with a resolution range of 0.68 − 3.50Å and mean of 1.91Å, a *R*_free_ range of 0.078− 0.30 and mean of 0.224. **G** includes 695 monomeric proteins. The percentage of missing atoms ranges from 0.0− 2.0% with a mean of 0.218% and a median of 0.046%. Neither **S** nor **G** includes any membrane proteins. **M** has 2, 487 structures of membrane proteins with a resolution range of 1.0 − 5.0Å and mean of 2.70Å, a *R*_free_ range of 0.096 − 0.458 and mean of 0.258. The percentage of missing atoms ranges from 0.0−2.0% with a mean of 0.253% and a median of 0.052%. Among the three sets, only the structures in **G** have no missing internal residues.

From the AlphaFold Protein Structure Database [25] we downloaded 350, 974 models for twenty seven proteomes including thirteen prokaryotic ones, *Escherichia coli, Haemophilus influenzae, Helicobacter pylori, Klebsiella pneumoniae, Methanocaldococcus jannaschii, Mycobacterium leprae, Mycobacterium tuberculosis, Neisseria gonorrhoeae, Pseudomonas aeruginosa, Salmonella typhimurium, Staphylococcus aureus, Streptococcus pneumoniae, Shigella dysenteriae*; and fourteen eukaryotic ones. The latter consists of three yeasts, *Saccharomyces cerevisiae, Schizosaccharomyces pombe* and *Candida albicans*; four plants, *Arabidopsis thaliana, Glycine max, Oryza sativa* and *Zea mays*; and an amoeba, *Dictyostelium discoideum*; an insect, *Drosophila melanogaster*; a worm, *Caenorhabditis elegans*; a fish, *Danio rerio*; and three mammals, *Mus musculus, Rattus norvegicus* and *Homo sapiens*.

The PDB files are preprocessed for area computation as described previously [26]. Specifically, if a PDB file includes multiple models, only the first one is selected, and if an atom has multiple conformations, only the first one is chosen. Protons are added using REDUCE [27] to any PDB structure or model that lacks them. Only protein atoms are used for area computation.

### 2.2 The computation of SES area and the definitions of patch areas per surface atom

The program for analytic SES area computation, SESA^6^, has been described previously [13]. The salient features of SESA relevant to the present study are the algorithmic separation of the exterior surface of a molecule from its interior surfaces, and the analytic and individual computations of the three types of SES patches. An interior surface is composed of surface atoms inaccessible to the bulk solvent without the rearrangement of their neighboring atoms. The analyses presented in this paper use only the SES of the exterior surface of a structure or a model. Specifically, we analyze the following geometric properties of SES: the number of surface atoms, and the SAS, probe and toroidal areas per surface atom defined as follows.

#### 2.2.1 The definitions of SAS, toroidal and probe areas per surface atom

Let **A** be the set of all the atoms in a molecule. To surface atom *i* ∈ **A** we assign a nonzero SES area *a*(*i*):

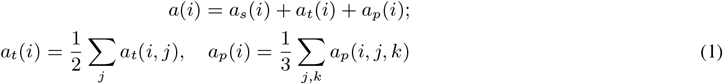

where *a*_*s*_(*i*), *a*_*t*_(*i*) and *a*_*p*_(*i*) are respectively the SAS, toroidal and probe areas for atom *i*; *a*_*t*_(*i, j*) is the total area of the toroidal patches determined by atom *i* and *j*, and *a*_*p*_(*i, j, k*) the total area of spherical polygons on a probe determined by atom *i, j* and *k*.

Given the individual patch areas for **A**, the total SES area *A*, the SAS, toroidal and probe areas per atom, *µ*_*s*_, *µ*_*t*_ and *µ*_*p*_, are computed as follows.

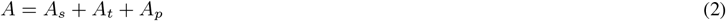

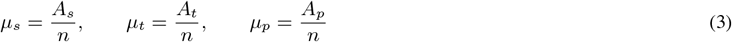

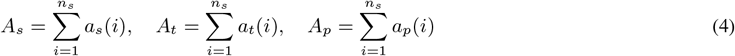

where *n* and *n*_*s*_ are, respectively, the numbers of total atoms and (exterior) surface atoms in a molecule, and *A*_*s*_, *A*_*t*_ and *A*_*p*_ are, respectively, SAS, toroidal and probe areas. The three summations are all over *n*_*s*_. SAS area *A*_*s*_, as a part of *A* (Eq. 2), differs from solvent-accessible surface area (SASA) with the latter being computed using a sphere radius of *r*_*a*_ + *r*_*w*_ where *r*_*a*_ and *r*_*w*_ are respectively the radii of a molecular atom and the probe [13]. The analyses presented in this paper focus on *µ*_*s*_, *µ*_*t*_, *µ*_*p*_ and *n*_*s*_ for the exterior surface of a molecule.

### 2.3 Fitting to power law and linear equation

The changes with *n* of the *µ*s for **G, S** and **M** and for each of the twenty seven AlphaFold proteomes are best-fitted using MATLAB to a power law with three coefficients, *a, b* and *c*.

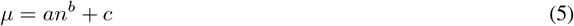

The point of intersection between the power laws for *µ*_*s*_ and *µ*_*p*_ is denoted as (*n*_*sp*_, *µ*_*sp*_). When the power laws are expressed in *n*_*r*_, the number of residues in a structure or a model, i.e., 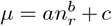, the point of intersection is denoted as (*N*_*sp*_, *µ*_*sp*_). The increase of *n*_*s*_ with *n* is fitted to a linear equation, *n*_*s*_ = *τn* + *b*, with a slope 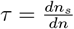. The changes of *µ*s with *n*, the power laws and their intersections, and the increase of *n*_*s*_ with *n* and its ratio *τ* are the main topics of this paper.

## 3 Results

In this section we describe the geometric properties of the molecular surfaces of both crystal structures and AlphaFold proteomes.

### 3.1 The geometric properties of the molecular surfaces of crystal structures

We first describe the changes of *µ*s with *n*, the power laws and their intersections, and then the increase of *n*_*s*_ with *n* and the linear equation.

#### 3.1.1 The changes of *µ*s with *n*, the power laws and their intersections for G, S and M

**G** has 8, 616 structures. The smallest and the largest ones are, respectively, 3FTR with 6 residues (76 atoms) and 2Y26 with 10, 080 residues (157, 300 atoms). As shown in Fig. 1a, both *µ*_*s*_ and *µ*_*t*_ decrease with *n* with the former faster than the latter while *µ*_*p*_ increases with *n*. The power laws for *µ*_*s*_ and *µ*_*p*_ are, respectively,

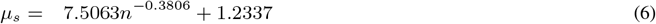

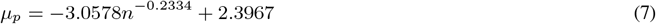

**Figure 1:**
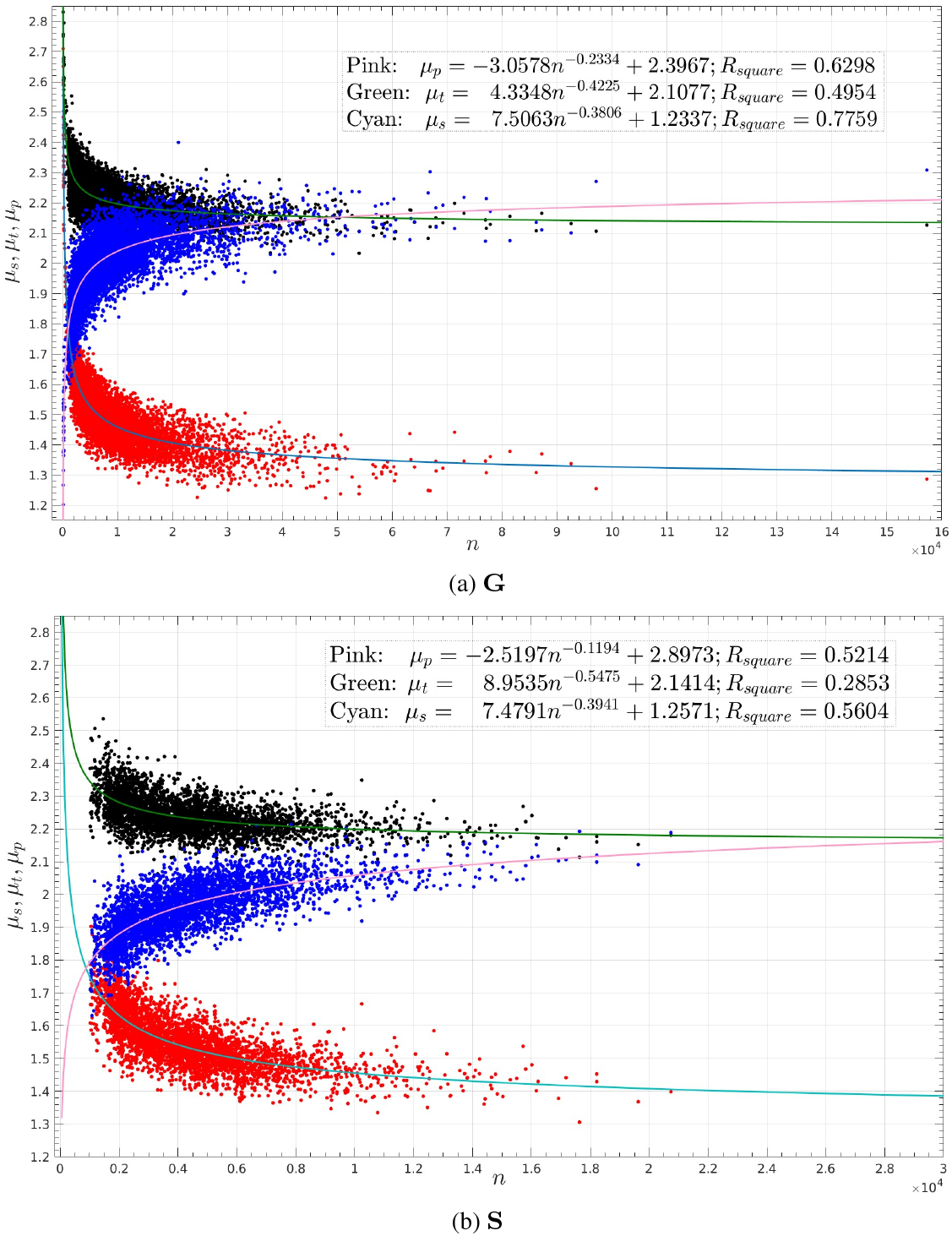
The *µ*_*s*_s, *µ*_*t*_s and *µ*_*p*_s of G and S and the power laws. Figs. (**a**) and (**b**) depict, respectively, the *µ*s and power law curves for **G** and **S**. The *µ*_*s*_s, *µ*_*t*_s and *µ*_*p*_s are colored, respectively, in red, black and blue with the power law curves colored respectively in cyan, green and pink. The inserts list the power laws and MATLAB fitting scores *R*_*square*_s. The x-axes are *n* and the y-axes are *µ*_*s*_, *µ*_*t*_ and *µ*_*p*_ in *Å*^2^.

They intersect at *n*_*sp*_ = 967 with *µ*_*sp*_ = 1.782. It equals 60 residues, i.e., *N*_*sp*_ = 60, using the fitted linear equation, *n*_*r*_ = 0.0647*n* −2.9278, where *n*_*r*_ is the number of residues in a protein (Fig. S1 of SI). In addition, the power law for *µ*_*p*_ intersects with that for *µ*_*t*_ at *n* = 50, 369 (3, 256 residues). However, the *µ*_*t*_s could not be fitted to a power law as well as either the *µ*_*s*_s or *µ*_*p*_s (Fig. 1a insert).

**S** has 4, 147 monomers. The smallest and largest ones are, respectively, 3ZR8 with 65 residues (1, 245 atoms) and 7MHM with 13, 158 residues (201, 246 atoms). As shown in Fig. 1b and as with **G**, both *µ*_*s*_ and *µ*_*t*_ decrease with *n* with the former faster than the latter while *µ*_*p*_ increases with *n*. The power laws for *µ*_*s*_ and *µ*_*p*_ are, respectively, *µ*_*s*_ = 7.4791*n*^−0.3941^ + 1.2571 and *µ*_*p*_ =−2.5197*n*^−0.1194^ + 2.8973. They intersect at *n*_*sp*_ = 875 with *µ*_*sp*_ = 1.775. Using the linear equation between *n*_*r*_ and *n, n*_*r*_ = 0.0643*n* + 0.9134 with a *R*_*square*_ = 0.9951, it equals *N*_*sp*_ = 57 residues, a purely theoretical number since all the monomers have *n*_*r*_ ≥ 65 residues. Furthermore, the differences in *µ*s between **S** and **G** are very small and their power laws are very close to each other (Table 1 and Fig. S2). In addition, the power law for *µ*_*p*_ intersects with that for *µ*_*t*_ at *n* = 33, 540 (2, 158 residues). However, the *µ*_*t*_s could not be fitted to a power law as well as either the *µ*_*s*_s or *µ*_*p*_s (Fig. 1b inserts).

**Table 1:** The differences in *µ*s between G and S, and between G and M. Column 1 lists set names. The differences of the *µ*_*s*_(*i*) and *µ*_*p*_(*i*) of structure *i* of either **S** or **M** from the *µ*_*s*_s and *µ*_*p*_s of **G** are quantified as 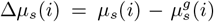 and 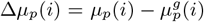,where 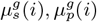 are computed using, respectively, Eqn. (6) and Eqn. (7) with *n* being the number of atoms in structure *i*. The smaller a Δ is, the closer the two power laws are. Column 2 lists the means and standard deviations of the lists of the Δ*µ*_*s*_s for **S** and **M**. Column 3 lists the means and standard deviations of the lists of the Δ*µ*_*p*_s for **S** and **M**. Column 4 lists *P*_*s*_s, the percentages of the structures in **S** and **M** whose Δ*µ*_*s*_ *>* 2*×* RMSE_*G,s*_, where RMSE_*G,s*_ is the RMSE (0.0532) of fitting to a power law the change with *n* of the *µ*_*s*_s of **G**. Column 5 lists *P*_*p*_s, the percentages of the structures in **S** and **M** whose Δ*µ*_*p*_ *>* 2 *×* RMSE_*G,p*_, where RMSE_*G,p*_ is the RMSE (0.0631) of fitting to a power law the change with *n* of the *µ*_*p*_s of **G**. Fig. S2 visualizes the *P*_*s*_ and *P*_*p*_ for **S**, and Figs. S3b and S3c visualize the *P*_*s*_ and *P*_*p*_ for **M**. Assuming a normal distribution, only 4.5% of the *µ*_*s*_s for **G** have a Δ*µ*_*s*_ *>* 2 *×* RMSE_*G,s*_, and only 4.5% of the *µ*_*p*_s for **G** have a Δ*µ*_*p*_ *>* 2*×* RMSE_*G,p*_. The smaller a *P*_*s*_ or a *P*_*p*_ is, the closer the two distributions are.

| Set | $\Delta\mu_s$ (mean / std) ($\text{\AA}^2$) | $\Delta\mu_p$ (mean / std) ($\text{\AA}^2$) | $P_s(\%)$ | $P_p(\%)$ |
| --- | --- | --- | --- | --- |
| **S** | -0.0121 / 0.0535 | 0.0070 / 0.0557 | 5.5 | 2.5 |
| **M** | -0.0335 / 0.0606 | -0.0155 / 0.0680 | 12.2 | 6.4 |

**M** has 2, 487 structures. The smallest and the largest ones are, respectively, 5N9H with 16 residues (244 atoms), and 3OAA with 13, 028 residues (198, 458 atoms). As shown in Fig. S3a and as with **S** and **G**, both *µ*_*s*_ and *µ*_*t*_ decrease with *n* with the former faster than the latter while *µ*_*p*_ increases with *n*. The power laws for *µ*_*s*_ and *µ*_*p*_ are, respectively, *µ*_*s*_ = 10.9135*n*^−0.4565^+1.2606 and *µ*_*p*_ = −4.1148*n*^−0.2491^+2.4396. They intersect at *n*_*sp*_ = 1, 061 with *µ*_*sp*_ = 1.714. It equals *N*_*sp*_ = 64 residues using the linear equation *n*_*r*_ = 0.0644*n* −4.7048 with a *R*_*square*_ = 0.9992. The *R*_*square*_ for *µ*_*s*_ is worse than those for both **G** and **S** (Fig. S3a). However, the differences in *µ*s between **M** and **G** remain small (Table 1 and Figs. S3b and S3c).

#### 3.1.2 The increases of *n*_*s*_ with *n*, and the linear equations for G, S and M

The linear equations between *n* and *n*_*s*_ for **G, S** and **M** are, respectively, *n*_*s*_ = 0.3770*n* + 499.3710 with a *R*_*square*_ = 0.9771 (Fig. S4a); *n*_*s*_ = 0.3952*n* + 297.2236 with a *R*_*square*_ = 0.9586; and *n*_*s*_ = 0.4234*n* + 559.2468 with a *R*_*square*_ = 0.9667. Their *τ* s (slopes), 0.3770, 0.3952 and 0.4234, are very close to each other.

### 3.2 The geometric properties of the molecular surfaces of AlphaFold models

We first describe the changes of *µ*s with *n*, the power laws and their intersections. We then describe the increases of *n*_*s*_ with *n* and the linear equations.

#### 3.2.1 The changes of *µ*s with *n* and the power laws and their intersections

The changes with *n* of the *µ*s of the AlphaFold models for each proteome are fitted to a power law as described above. We illustrate the variations, the power laws and their intersections using the *E*.*coli* proteome as a representative of the thirteen prokaryotic proteomes and the human proteome as a representative of the fourteen eukaryotic ones. Please refer to Table S1 for the *R*_*square*_s and points of intersection (*N*_*sp*_s) for the twenty seven proteomes. In addition, Figs. 4 and 5 depict the power laws and their *n*_*sp*_s, and Figs. S9–S35 depict the changes with *n* of their *µ*s and *n*_*s*_s.

AlphaFold predicted 4, 363 models for *E*.*coli* proteome. The smallest and largest ones are, respectively, 257 atoms (16 residues), and 34, 552 atoms (2, 358 residues). As with **S, G** and **M**, both *µ*_*s*_ and *µ*_*t*_ decrease with *n* with the former faster than the latter (Figs. 2c, 2d and S9) while *µ*_*p*_ increases with *n*. The power laws for *µ*_*s*_ and *µ*_*p*_ are, respectively, *µ*_*s*_ = 6.4488*n*^−0.3261^ + 1.1725 and *µ*_*p*_ = −3.8938*n*^−0.2619^ + 2.3657. They intersect at *n*_*sp*_ = 1, 393 (*N*_*sp*_ = 87 residues (the legend of Fig. S1)). In addition, the power law for *µ*_*p*_ intersects with that for *µ*_*t*_ at *n* = 42, 897, corresponding to 2, 681 residues, larger than the *n*_*r*_ for any *E*.*coli* protein.

**Figure 2:**
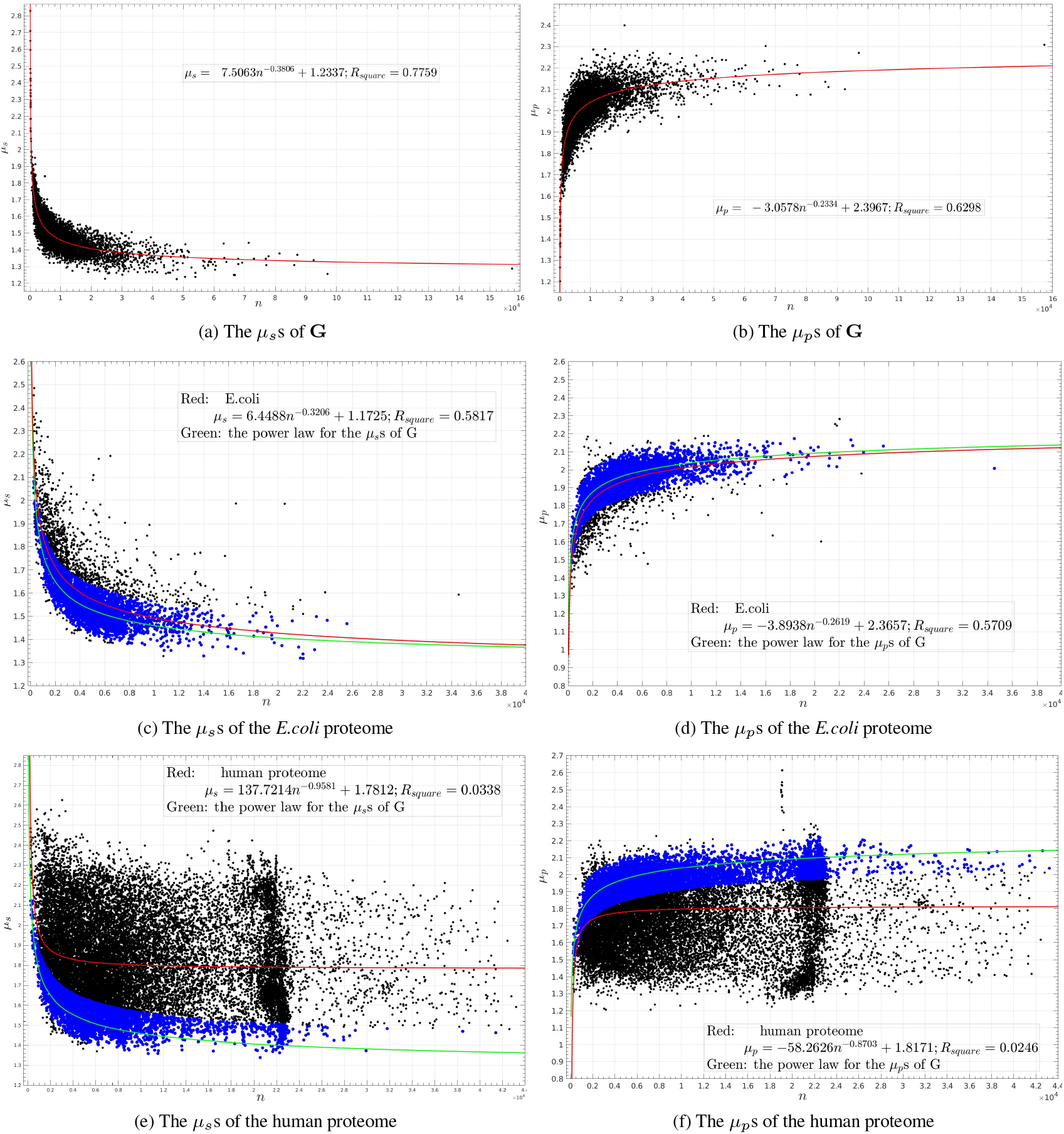
The changes with *n* of the *µ*_*s*_s and *µ*_*p*_s for G, and the *E*.*coli* and human proteomes. Each point represents either a *µ*_*s*_ (**a, c, e**) or a *µ*_*p*_ (**b, d, f**) value for a structure or a model. The power law curves are shown in red. In Figs. (**c, d, e, f**), the corresponding power law curves for **G** are shown in green. The insert in each figure lists the power law and *R*_*square*_. About 24% of the models for the *E*.*coli* proteome have a Δ*µ*_*s*_ *>* 2*×* RMSE_*G,s*_, where RMSE_*G,s*_ = 0.0532, and about 13% have a Δ*µ*_*p*_ *>* 2 *×* RMSE_*G,p*_, where RMSE_*G,p*_ = 0.0631 (Table 2). In stark contrast, about 80% of the models for the human proteome have a Δ*µ*_*s*_ *>* 2 *×* RMSE_*G,s*_, and about 61% have a Δ*µ*_*p*_ *>* 2 × RMSE_*G,p*_. In Figs. (**c, e**) the *µ*_*s*_s with Δ*µ*_*s*_ *<* 2 *×* RMSE_*G,s*_, and in Figs. (**d, f**) the *µ*_*p*_s with Δ*µ*_*p*_ *<* 2 *×* RMSE_*G,p*_ are colored in blue, and the rest in black. In Table 2, the percentages of the black data points over the total number of data points are listed, respectively, as *P*_*s*_ and *P*_*p*_. Assuming a normal distribution, only 4.5% of the *µ*_*s*_s for **G** have a Δ*µ*_*s*_ *>* 2 *×* RMSE_*G,s*_, and only 4.5% of the *µ*_*p*_s for **G** have a Δ*µ*_*p*_ *>* 2 *×* RMSE_*G,p*_. The x-axes are *n*. The y-axes are, respectively, *µ*_*s*_ (**a, c, e**) and *µ*_*p*_ (**b, d, f**) in *Å*^2^.

AlphaFold predicted 23, 391 models for the human proteome. The smallest and largest ones are, respectively, 238 atoms (16 residues) and 43, 190 atoms (2, 699 residues). As shown in Figs. 2e and 2f, in stark contrast to the *µ*s for both the crystal structures (Figs. 1, 2a, 2b and S3) and the models for the thirteen prokaryotic proteomes (Figs. 2c, 2d and S9–S21), the *µ*s for the fourteen eukaryotic proteomes (Figs. S22–S35) differ largely from both of them in the sense that when plotted with *n* as x-axis and *µ* as y-axis, the *µ*s are broadly distributed and hardly change with *n*. The changes of the *µ*s with *n* follow only minimally a power law with a *R*_*square*_ = 0.0338 for *µ*_*s*_ and *R*_*square*_ = 0.0246 for *µ*_*p*_. The power laws for *µ*_*s*_ and *µ*_*p*_ intersect at *n*_*sp*_ = 11, 100 (*N*_*sp*_ = 694 residues (the legend of Fig. S1)). However, the power law for *µ*_*p*_ *does not* intersect with that for *µ*_*t*_.

**Table 2:** The differences in SES between the 350, 974 AlphaFold models for twenty seven proteomes and crystal structures. Column 1 lists organism names in abbreviation with their full names listed in Table S3. The differences in *µ*_*s*_ and *µ*_*p*_ of model *i* of proteome from the *µ*_*s*_s and *µ*_*p*_s of **G** are quantified as 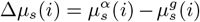 and 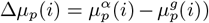 where 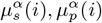 are the *µ*_*s*_, *µ*_*p*_ values for model *i*, and 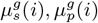 are computed using, respectively, Eqn. (6) and Eqn. (7) with *n* being the number of atoms of model *i*. Column 2 lists the mean and standard deviation of the list of the Δ*µ*_*s*_s for a proteome, and column 3 lists the mean and standard deviation of the list of the Δ*µ*_*p*_s for a proteome. Columns 4 and 5 list, respectively, the *P*_*s*_s and *P*_*p*_s for **S, M** and the twenty seven proteomes. They have the same meanings as those in Table 1. Fig. 2 visualizes the *P*_*s*_s and *P*_*p*_s for the *E*.*coli* and human proteomes. The rows for **S** and **M** are repeated here for easy comparison. As shown in the last row, the *µ*s, *P*_*s*_ and *P*_*p*_ for the human proteome are the furthest from those for **G**.

| Organism | $\Delta\mu_s$ (mean / std) ($\text{\AA}^2$) | $\Delta\mu_p$ (mean / std) ($\text{\AA}^2$) | $P_s$(%) | $P_p$(%) |
| --- | --- | --- | --- | --- |
| **Crystal structures** |  |  |  |  |
| **S** | **-0.0121 / 0.0535** | **0.0070 / 0.0557** | **5.5** | **2.5** |
| **M** | **-0.0335 / 0.0606** | **-0.0155 / 0.0680** | **12.2** | **6.4** |
| **Prokaryotic proteomes** |  |  |  |  |
| **E.coli** | *0.0512 / 0.1011* | *-0.0357 / 0.0825* | 23.7 | 12.8 |
| haeIn | 0.0505 / 0.0994 | -0.0318 / 0.0811 | 21.2 | 11.7 |
| helpy | 0.0609 / 0.1119 | -0.0409 / 0.0894 | 26.7 | 16.6 |
| klePh | 0.0557 / 0.1047 | -0.0453 / 0.0873 | 25.1 | 16.5 |
| **metJa** | **0.0310 / 0.0850** | **-0.0156 / 0.0724** | **16.4** | **8.6** |
| mycLe | 0.0960 / 0.1362 | -0.0645 / 0.1090 | 37.0 | 25.0 |
| mycTu | 0.0859 / 0.1391 | -0.0752 / 0.1098 | 35.7 | 26.9 |
| neiG1 | 0.0761 / 0.1276 | -0.0511 / 0.1027 | 32.3 | 21.8 |
| pseAr | 0.0529 / 0.1117 | -0.0521 / 0.0858 | 25.2 | 16.2 |
| salTy | 0.0569 / 0.1046 | -0.0384 / 0.0846 | 25.7 | 14.0 |
| shiDs | 0.0604 / 0.1067 | -0.0429 / 0.0883 | 27.0 | 16.7 |
| staA8 | 0.0553 / 0.1120 | -0.0236 / 0.0914 | 23.1 | 12.5 |
| strR6 | 0.0549 / 0.0997 | -0.0296 / 0.0837 | 24.4 | 13.7 |
| **Yeast proteomes** |  |  |  |  |
| schpo | 0.2043 / 0.1976 | -0.1220 / 0.1601 | 62.5 | 41.4 |
| yeast | 0.2295 / 0.2063 | -0.1366 / 0.1647 | 66.2 | 46.1 |
| canAl | 0.2360 / 0.2129 | -0.1348 / 0.1655 | 66.6 | 44.0 |
| **Plant proteomes** |  |  |  |  |
| araTh | 0.2539 / 0.2087 | -0.1511 / 0.1699 | 71.8 | 50.3 |
| soyBn | 0.2555 / 0.2112 | -0.1508 / 0.1755 | 71.8 | 50.3 |
| maize | 0.2900 / 0.2140 | -0.1844 / 0.1764 | 77.6 | 58.5 |
| orySj | 0.3075 / 0.2251 | -0.1896 / 0.1854 | 77.9 | 60.9 |
| **Eukaryotic proteomes** |  |  |  |  |
| celEg | 0.2203 / 0.2067 | -0.1240 / 0.1637 | 64.0 | 41.6 |
| dicDi | 0.2556 / 0.2125 | -0.1251 / 0.1808 | 70.7 | 46.0 |
| danre | 0.2809 / 0.2153 | -0.1793 / 0.1858 | 76.3 | 53.2 |
| drome | 0.2883 / 0.2335 | -0.1882 / 0.1946 | 73.0 | 54.0 |
| mouse | 0.2798 / 0.2290 | -0.1917 / 0.1910 | 73.0 | 54.7 |
| rat | 0.2763 / 0.2272 | -0.1896 / 0.1894 | 72.8 | 55.0 |
| **human** | **0.3181 / 0.2287** | **-0.2219 / 0.2025** | **80.2** | **60.8** |

#### 3.2.2 The increases of *n*_*s*_ with *n* and the linear equations

The linear equations for the twenty seven proteomes are shown in Figs. 3, S5a and S5b with their *τ* s listed in their inserts and Table S1. As in the last section, we illustrate the variations using the *E*.*coli* proteome as a representative of the thirteen prokaryotic proteomes and the human proteome as a representative of the fourteen eukaryotic proteomes.

**Figure 3:**
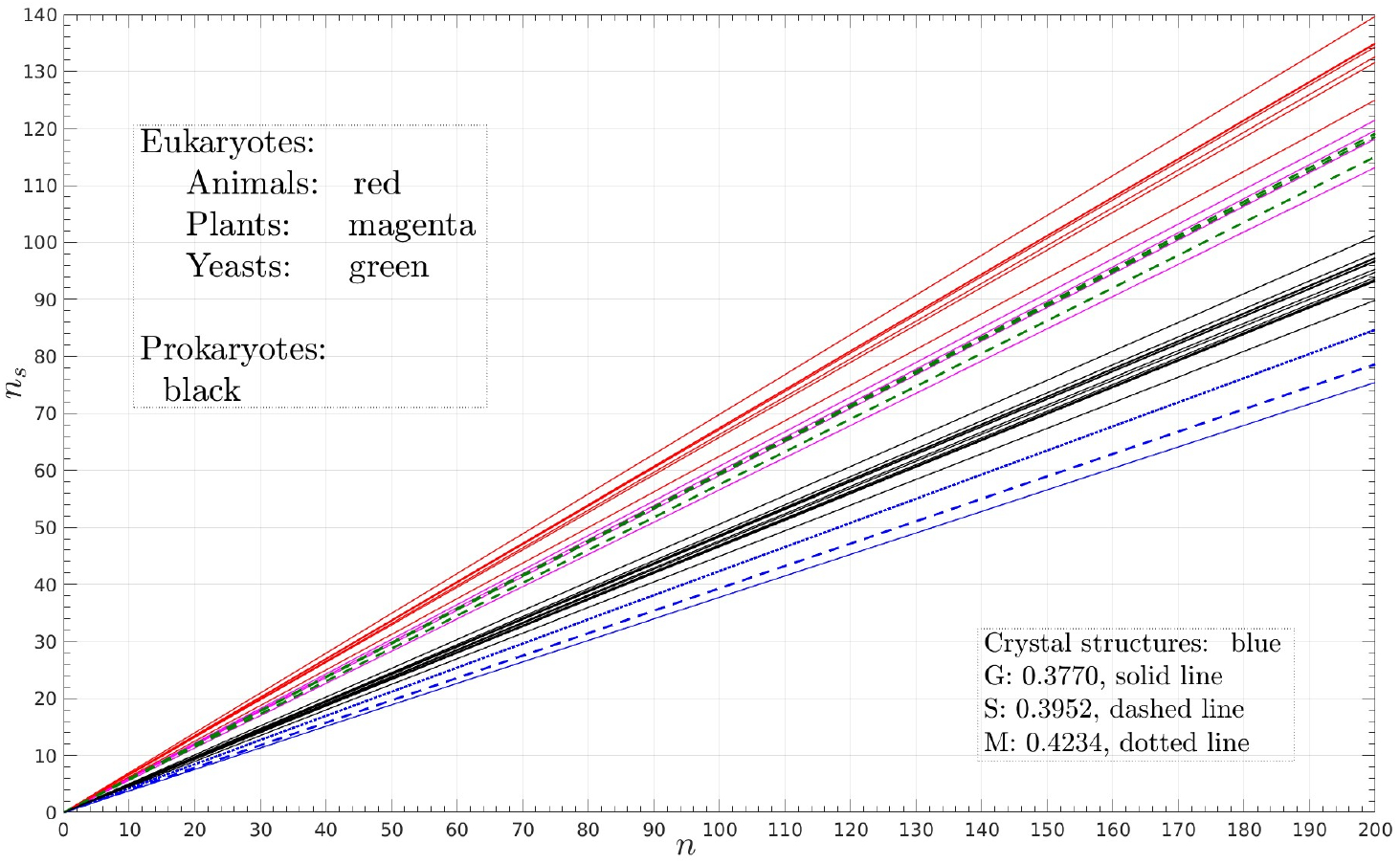
The linear equations for the increase of *n*_*s*_ with *n*. The lines for the fourteen eukaryotic proteomes are colored respectively in red (animals), magenta (plant) and yeast (green). The lines for the thirteen prokaryotic proteomes are colored in black, and those for the crystal structures in blue. The lines all pass through the origin to highlight their differences in *τ*. Table S1 and the inserts in Fig. S5 list their *τ* values. The *τ* s for the crystal structures are all smaller than those for the prokaryotic proteomes and the latter are all smaller than those for eukaryotic proteomes. The x-axis is *n*, and the y-axis is *n*_*s*_.

**Figure 4:**
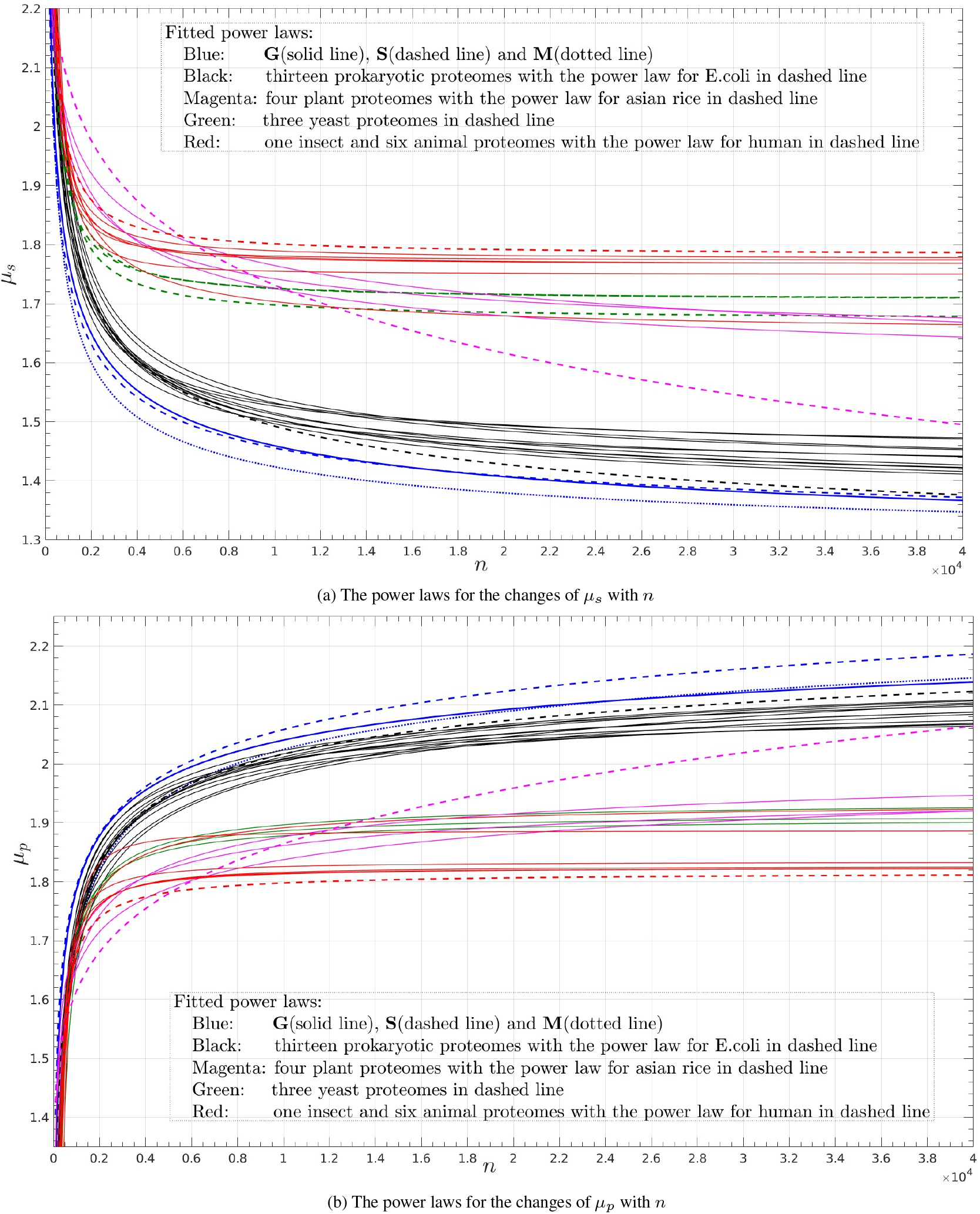
The power laws for G, S, M and the twenty seven proteomes. Figs. (**a**) and (**b**) show, respectively, the power law curves for *µ*_*s*_ and *µ*_*p*_. From bottom up, the three blue curves in (**a**) are, respectively, the power laws for **M, S** and **G**. Right above them are the power law curves for the thirteen prokaryotic proteomes colored in black. Above them are the power law curves for the fourteen eukaryotic proteomes, each of them is well separated from the curves for both the crystal structures and prokaryotic proteomes. The thirty power law curves for *µ*_*p*_ (Fig. (**b**) are similarly distributed but in a top down manner. Please see the inserts for coloring scheme. The x-axes are *n*, and the y-axes are *µ*_*s*_ (**a**) and *µ*_*p*_ (**b**) in *Å*^2^.

**Figure 5:**
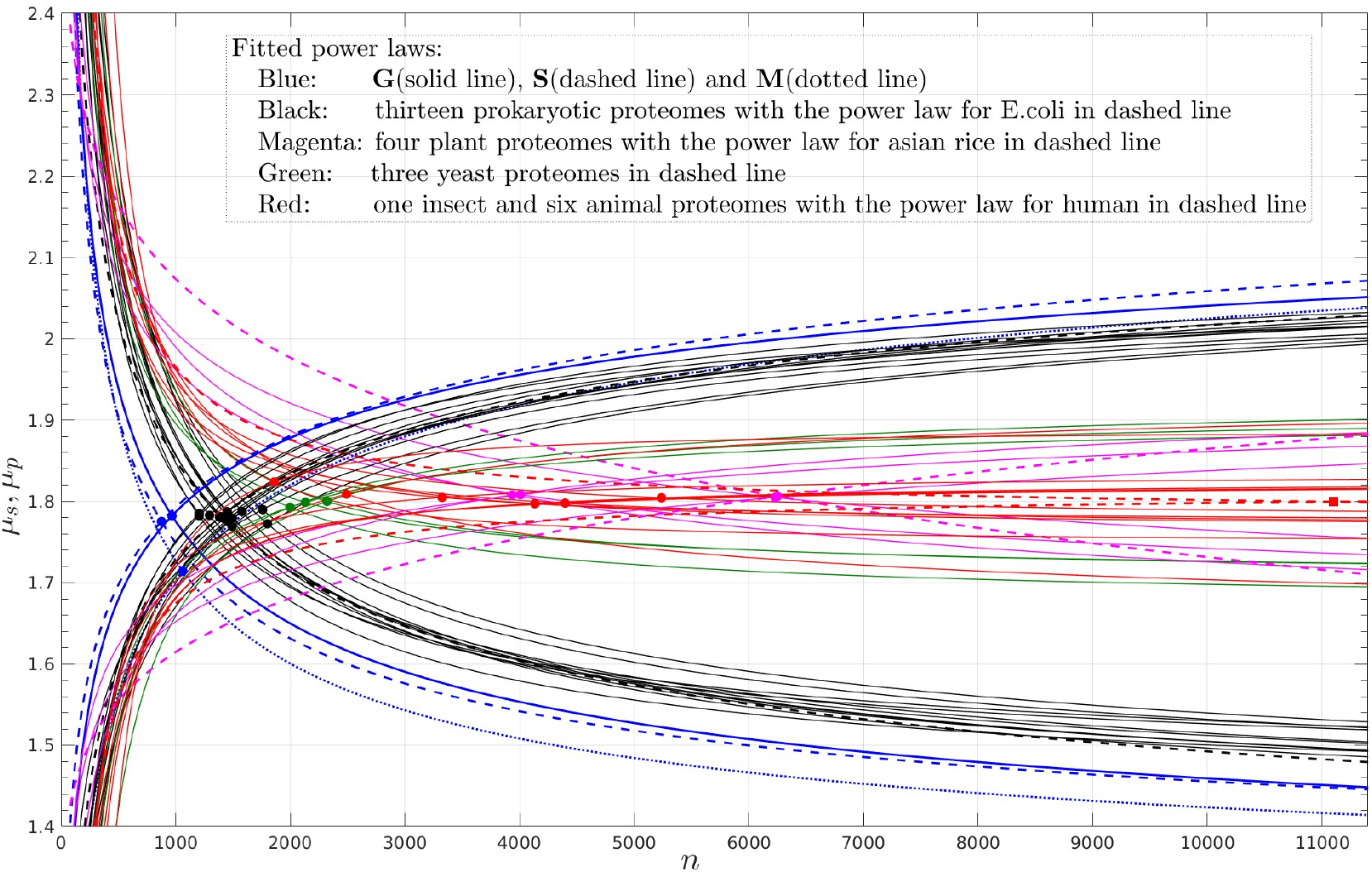
The *n*_*sp*_s for G, S, M and the twenty seven proteomes. Except for the human proteome whose *n*_*sp*_ is depicted as a square, the rest are depicted as a filled circle. The *n*_*sp*_s for the crystal structures are all smaller than any of those for the twenty seven proteomes, and the *n*_*sp*_s for the prokaryotic proteomes are all smaller than any for the eukaryotic proteomes (Table S1). **S** has the smallest *n*_*sp*_ = 875 while the human proteome the largest *n*_*sp*_ = 11, 000. The x-axis is *n* and the y-axis is *µ*_*s*_, *µ*_*p*_ in *Å*^2^.

The line for the *E*.*coli* proteome, *n*_*s*_ = 0.4659*n* + 263.2263, has a *τ* = 0.4659, which is only 1.2-fold larger than that for **G** (0.3770, Figs. 3 and S4b). In contrast, the line for the human proteome (Figs. 3, S4c and S5b), *n*_*s*_ = 0.6742*n* −38.9199, has a *τ* = 0.6742, 1.8-fold larger than that for **G**. As shown in Fig. S4c, the increase with *n* of the *n*_*s*_s for the human proteome looks artificial.

## 4 Discussion

We start with the previous analyses of experimentally-derived protein structures and their surfaces. Next we describe the meanings of the geometric properties *µ*_*s*_, *µ*_*p*_ and *τ* and follow up with the discussion of the geometric properties of the SESs for both crystal structures and AlphaFold models. Specifically, we discuss first the geometric properties of the crystal structures and their implications for protein structure and folding, and then those of the AlphaFold models and their implications for the quality and application of AlphaFold prediction. Finally we discuss the possible limitations of this study.

### 4.1 Previous analyses of protein structures and their surfaces

Previous analyses of experimental protein structures have focused primarily on their folds, secondary structures, binding or active sites and structural quality [28], while their surfaces have received much less attention. Out of the three surface types, previous analyses have largely been performed on SASA and at residue-level rather than on SAS (as a part of SES) or SES and at atom-level, even though it has been shown that SES could be better suited for quantifying the contribution of surface to protein structure and function [29], possibly due to the lack of robust programs for accurate SES computation. Specifically, except for SESA no robust and accurate programs are available for individual computation of the three SES patch types. Furthermore, to our best knowledge none of the three SES types have been studied *separately and at atom-level* for their possible relevance to protein structure, folding and function.

### 4.2 The geometric properties *µ*_*s*_, *µ*_*p*_ and *τ* describe the packing of surface atoms

A SAS patch of surface atom *i* is a spherical polygon with up to a few tens of sides [13]. Each side corresponds to an intersection between *i* and a neighboring atom with each of them modeled as a sphere with a radius of *r*_*a*_ + *r*_*p*_ where *r*_*a*_ and *r*_*p*_ are respectively atomic and probe radii. A toroidal patch exists when two neighboring atoms are close enough to form an impenetrable barrier to the probe, and a probe patch exists when a triple of them are close enough to form such a barrier. Thus the *µ*s are averages over the spatial distribution of surface atoms in a molecule. Specifically, a large *µ*_*s*_ means that either *n*_*s*_ is large, i.e., many atoms are on surface, or *a*_*s*_ per surface atom is large (Eqs. (3, 4)). A large *µ*_*p*_ means that most of the surface atoms are close enough to form an impenetrable barrier to the probe. Since the vast majority of the surface atoms of a polypeptide come from its side chains, a large *µ*_*s*_ but a small *µ*_*p*_ indicates that the surface side chains are loosely packed with large space among them, i.e., most side chain atoms are largely exposed, a hallmark of the structure of a typical peptide. By contrast, a large *µ*_*p*_ but a small *µ*_*s*_ indicates that the surface side chains are tightly packed with the space among them largely impenetrable to the probe, a hallmark of a folded and spherical-shaped protein. Thus we could use the geometric property of SES, 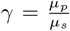,to characterize the spatial distribution of surface atoms.

The three power laws and their points of intersection could be looked upon as an average over either a set of crystal structures or an AlphaFold proteome. The power law for *µ*_*s*_ intersects with that for *µ*_*p*_ when *γ* = 1 or *n* = *n*_*sp*_, i.e., when the total SAS area of a molecule equals its total probe area. A SAS patch is convex while a probe one concave, thus the two power laws intersect when the total area of the patches with a negative curvature equals that with a positive curvature (if one ignores the toroidal patches, each of which includes regions of positive and negative curvatures). A typical peptide has a *γ <* 1 while a folded protein usually has *γ >* 1.

The *τ* for a set of crystal structures or a proteome describes, on average, how fast *n*_*s*_ increases with *n*. As shown in Fig. 3, the *τ* for **G** (0.3770), is very close to that for **S** (0.3952). Both of them are smaller than that for **M** (0.4243). A small *τ* together with a large *γ* is a feature of a set of well-packed crystal structures of soluble proteins. The simultaneous increase of *n*_*s*_ and decrease of *a*_*s*_ occur in the transmembrane region of a membrane protein where hydrogen atoms are more abundant than in the rest of the protein (Fig. S6).

### 4.3 The geometric properties of the molecular surfaces of crystal structures

We have analyzed the changes of *A*_*s*_, *A*_*t*_, *A*_*p*_ and *A* (Eq. 2) with *n, n*_*r*_ and *n*_*s*_, and with secondary structure content and surface atom type. Among them we find that the changes with *n* of *µ*s, particularly *µ*_*s*_ and *µ*_*p*_, are most characteristic (Figs. 1 and 2) and thus likely most relevant to protein structure, folding and function. The *µ*s for the crystal structures are fitted to *n* slightly better than to either *n*_*s*_ or *n*_*r*_ (Fig. S7), though there exist excellent linear relationships among them (Figs. S1 and S4a). For example, the number of atoms per residue, i.e., 16 atoms per residue, could be determined accurately from the linear equation between *n*_*r*_ and *n* for **G** (Fig. S1). The fitting to *n* is better than to *n*_*r*_ likely because of the existence of missing atoms in many crystal structures, and to *n*_*s*_ likely because of the diversity in surface shape among the structures. Furthermore, the *µ*s change only slightly with secondary structure content and the composition of surface atom types (data not shown). In the following we focus our discussion on the changes with *n* of *µ*_*s*_ and *µ*_*p*_, and *N*_*sp*_.

#### 4.3.1 The characteristic changes of the *µ*s with *n* and their geometric and biophysical meanings

The characteristic changes of the *µ*s with *n* could be (1) a global geometrical feature, (2) the optimization of solvent-protein interaction, and (3) the minimization of misinteraction.

##### 4.3.1.1 The change of *µ*s with *n* is unlikely a global geometric feature

SES is computed with the molecule modeled as a set of possibly intersecting spheres having different radii. With an increasing number of spheres, as a whole such a set is expected to become more and more similar to a large sphere with a small positive curvature. This is in contrast with the simultaneous increase of *µ*_*p*_ and decrease of *µ*_*s*_ with *n*, and the fact that *N*_*sp*_ ≤ 65 for **S, G** and **M**. The power laws and the *N*_*sp*_ values show that with an increasing number of atoms (spheres) and particularly after the point of intersection, the total area of the regions of negative curvature becomes larger than that of positive curvature. In other words, the results show that both *µ*_*s*_ and *µ*_*p*_ are only local geometric properties of SES, and as such they should not change with molecular size.

##### 4.2.1.2 The optimization of solvent-protein interaction

If the representation of solvent molecules as spheres is a decent approximation, the increase of *µ*_*p*_ with *n* means that the surface area where solvent molecules could interact more optimally with the protein increases with *n*. The solvent could be either the aqueous solvent for a water-soluble protein or both the lipids and aqueous solvent for a membrane protein. In other words, the increase in *µ*_*p*_ suggests that the disruption to solvent’s intrinsic structure is reduced. For a water-soluble protein this means that the disruption to water hydrogen-bond network is reduced, the entropic increase of the whole system reduced, and the inter-molecular hydrogen bonding and VDW interaction optimized. For a membrane protein, the disruption to the structural integrity of the membrane is reduced.

#### 4.3.1.3 The minimization of protein-protein misinteraction

A naturally-evolved protein must guarantee that the disruptions to and by all its neighbors are minimum in terms of free energy [30]. In other words, in a crowded intracellular environment, except for its cognate binding partners, a naturally-evolved protein should avoid binding to other molecules. A possible way to prevent noncognate binding is to have a molecular surface with a negative area-weighted average curvature. With a larger probe area, solvent-protein interaction becomes stronger, and that in turn makes it less likely the occurrence of intermolecular hydrophobic interactions between hydrophobic surface side chains, and that in turn reduces the chance for two proteins to stick together and for them to further form inclusion bodies inside a cell.

#### 4.3.2 A hypothesis on protein structure and folding from a molecular surface perspective

Up to now, no satisfactory explanations have been given to the occurrence of the transition at about 50 residues from a peptide with a broad conformational distribution to a protein with a narrow one. A polypeptide chain with *<* 50 residues is typically called a peptide. It is not known whether the transition is determined by geometric properties, biophysical properties, solvent-protein interaction, or something else. It is suggested to be of geometric origin [31] based purely on theoretical calculation: the surface to volume ratio of a globular protein does not resemble that of a prolate ellipsoid until its chain length is ≥ 50 residues, and of biophysical origin since a polypeptide chain has to be long enough to enclose an inside in which hydrophobic residues could be sequestered. However, even for membrane proteins (set **M**), *µ*_*s*_ decreases and *µ*_*p*_ increases with *n*, and their power laws intersect at 64 residues. Furthermore, if the sequester of hydrophobic residues is the sole driving force, then after the point of intersection, *µ*_*s*_ and *µ*_*p*_ should remain constant. Thus the sequester of hydrophobic residues could provide only a partial explanation.

The *N*_*sp*_s for **G** and **S** are respectively 60 and 57, close to the number of residues required for a polypeptide to adopt a dominant conformation in solution. It suggests that a surface with a zero, or better, a negative average curvature is required for the folding of a polypeptide into a 3D structure with a dominant conformation in solution. In other words, the geometric properties of the SESs of the crystal structures provide a plausible explanation for the transition from peptide to protein. In turn, it suggests that 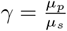 is an important parameter for characterizing the transition, a type of phase transition, from a form resembling liquid to a form resembling solid. We hypothesize that *γ* is closely related to the structural stability of proteins [32] and protein folding in the sense that *γ >* 1 is a necessary condition for protein folding.

### 4.4 The geometric properties of the molecular surfaces of AlphaFold models

The neural-network based AlphaFold [14, 15, 16, 17] has been hailed as a breakthrough in protein structure prediction based on several criteria for accuracy and structural quality [20, 21, 22, 23, 24, 28]. To objectively assess AlphaFold’s accuracy, some models have been compared directly with crystallographic maps [18]. In addition, the topology of the models has been evaluated [33]. However, to our best knowledge, neither their SASs nor SESs have been analyzed.

The AlphaFold models studied here are for monomeric proteins containing entire protein sequence. For a fair comparison with them the crystal structures of non-membrane proteins have been separated into two sets **S** and **G** with the former including only monomeric proteins. In addition, the structures in **G** have no missing internal residues and an average of 0.218% missing atoms per structure. As shown in Table 1 and Figs. 4, 5 and S2, the power laws and *τ* s for **S** and **G** are very close to each other. In addition, the power laws for the SESs of the set of the individual chains of the multimers in **G** are almost identical to those for **G** itself (section S8 and Fig. S8), and the *τ* for the set of chains is 0.4087, only 8.4% larger than that for **G** itself. In other words, though **G** consists mainly of multimers, it introduces only negligent errors when it is used to represent the crystal structures of monomeric proteins. Consequently, the comparisons with the AlphaFold models are presented with references to the geometric properties of the SESs of **G**.

#### 4.4.1 The similarity in SES between the crystal structures and AlphaFold models for prokaryotic proteomes

The changes with *n* of the *µ*s of the AlphaFold models for the thirteen prokaryotic proteomes resemble those for the crystal structures (Tables 2 and S1, and Figs. 4, 5 and S9–S21). Among them the models of the *E*.*coli* proteome have geometric properties quite similar to those of the crystal structures (Figs. 2c, 2d and S9). In addition, the *τ* s for the thirteen proteomes range from 0.4497 −0.5057 (Figs. 3 and S5a, and Table S1), all larger than any of the *τ* s for the crystal structures, but smaller than any for the fourteen eukaryotic proteomes. This shows that though the SESs of the models for prokaryotic proteomes differ only slightly from those for crystal structures, their surface side chains are still packed less tightly than those of crystal structures.

The sensitivity of neural-network to training data [34] suggests that the resemblances result likely from (1) the abundance of sequenced prokaryotic genomes, (2) the similarity among prokaryotic genomes [35], (3) the high experimental coverage of at least one of the thirteen prokaryotic proteomes [35], (4) prokaryotic proteins are generally smaller in size and simpler in structure than their counterparts in eukaryotic proteomes, (5) the low content of disordered regions in prokaryotic proteomes [36], and (6) the low level of glycosylation during post-translation modification. In other words, the abundance of high quality training data somewhat compensates for the lack of surface features for neural-network model training.

#### 4.4.2 The large differences in SES between the crystal structures and AlphaFold models for eukaryotic proteomes

As illustrated in Figs. 2e and 2f, and shown in Tables 2 and S1 and Figs. S22–S35, the *µ*s of the AlphaFold models for each of the fourteen eukaryotic proteomes are broadly distributed, hardly change with *n* and only minimally follow a power law. Among them the *µ*s for the human proteome (Figs. 2e and 2f) are the furthest from those for crystal structures. In addition, their *τ* s range from 0.5752 −0.6984 (Figs. 3 and Table S1); all are larger than any of the *τ* s for the thirteen prokaryotic proteomes, and much larger than any for the crystal structures. This means that for the same *n*, the models for the eukaryotic proteomes have more surface atoms per model than either the crystal structures or the models for prokaryotic proteomes. In other words, the surface side chains of these models are peptide-like. Among them the human proteome has the largest *N*_*sp*_ = 694, *P*_*s*_ = 80.2% and *P*_*p*_ = 60.8% (Table 2 and Figs. 2e and 2f), and the third largest *τ* = 0.6742 (Table S1). The differences suggest that in terms of SES, either the models are accurate and consequently there exist genuine variations between them, or the models have errors. If the latter is true, it means that the current AlphaFold models for eukaryotic proteomes are not adequate for structure-based drug design in particular.

##### 4.4.2.1 Possible reasons for the genuine differences

It is estimated that there exist 30% intrinsically-disordered regions in a eukaryotic proteome. The SES of a disordered region resembles the SES of a typical peptide with a *γ <* 1.0. However, as shown in Figs. 2e and 2f and Table 2, 80.2% of the models for the human proteome have a Δ*µ*_*s*_ *>* 2 *×*RMSE_*G,s*_. In **G** itself only 4.5% structures have a Δ*µ*_*s*_ *>* 2 *×*RMSE_*G,s*_, thus about 75.7% (75.7 = 80.2 −4.5) of the AlphaFold models have a *µ*_*s*_ outside of the *µ*_*s*_ range^7^ for **G**. In other words, the estimated 30% disordered regions is not large enough to account for the 75.7% difference in *µ*_*s*_ between the crystal structures and the models for the human proteome. Among them *Schizosaccharomyces pombe* proteome has the smallest *P*_*s*_ = 62.5%, meaning that 58.0% of its models have a *µ*_*s*_ outside of the *µ*_*s*_ range for **G**. It also has the smallest |Δ*µ*_*s*_| and |Δ*µ*_*p*_| (Table 2).

A eukaryotic proteome includes about 30% membrane proteins. However, the power laws for the *µ*s, *N*_*sp*_ and *τ* of **M** are all much closer to those for **G** than to any of the eukaryotic proteomes (Figs. 3, 4, 5 and S3).

The low-confidence regions of an AlphaFold model, as marked by a low pLDDT score, and often corresponding to disordered regions, are included in our calculation. If the differences are attributed solely to such regions, then it means that about 75.7% of the surface residues of the models for the human proteome have a low pLDDT score, which seems to disagree largely with the current quality assessment of AlphaFold models [17, 18]. It has been claimed that the high-confidence regions of an AlphaFold model are usually very close to experimental structures [15, 17].

##### 4.4.2.2 The differences are due possibly to the lack of surface features in AlphaFold model training

The models are predicted using an AI model trained with the currently available crystal structures and other data. Thus we are puzzled by the up to 75.7% difference in SES between the models for eukaryotic proteomes and the crystal structures. At present we think that the large differences are due possibly to the lack of surface features in AI model training, which seems to affect the eukaryotic models much more than the prokaryotic models.

#### 4.4.3 The large differences in SES between prokaryotic and eukaryotic proteomes

It is somewhat surprising that there exist large differences in SES between the models for prokaryotic and eukaryotic proteomes (Figs. 2-6, and Tables 2, S1 and S2) since they are all predicted using the same AI model. In the following we discuss two possible explanations: (1) the differences result from genuine structural variations between them, and (2) there exist errors in the models for eukaryotic proteomes.

##### 4.4.3.1 The differences reflect the genuine variations between prokaryotic and eukaryotic proteomes

A eukaryotic proteome has a higher percentage of disordered regions than a prokaryotic one. It is estimated that on average a eukaryotic protein contains 32% disordered residues while a prokaryotic one only 12% [36]. However, the 20% difference is unlikely to be large enough to rationalize 53.2% difference^8^ in *µ*_*s*_ between the models for the human and the *E*.*coli* proteomes (Fig. 6 and Table S2), even though it has been shown that the disordered regions of several AlphaFold models agree well with experimental results [24].

**Figure 6:**
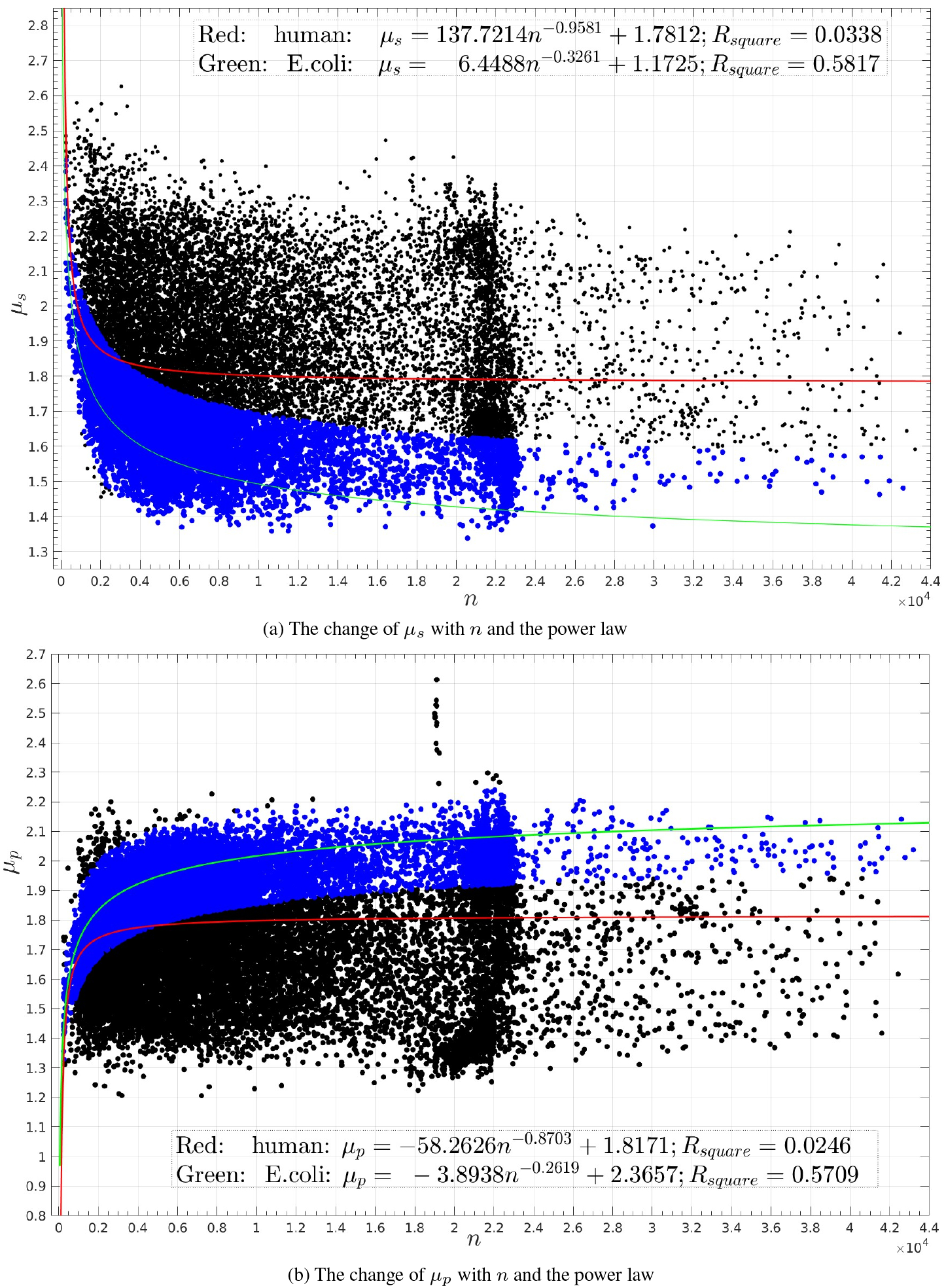
The changes with *n* of the *µ*s for the human proteome and the comparisons with those for the *E*.*coli* proteome. Figs. (**a**) and (**b**) depict, respectively, the *µ*_*s*_s and *µ*_*p*_s for the human proteome. The power law curves for the human and *E*.*coli* proteomes are colored respectively in red and green. In Figs. (**a**) the *µ*_*s*_s with Δ*µ*_*s*_ *<* 2 *×* RMSE_*E,s*_, where RMSE_*E,s*_ = 0.1002 is the RMSE of fitting the *µ*_*s*_s of the *E*.*coli* proteome to a power law, and in Figs. (**b**) the *µ*_*p*_s with Δ*µ*_*p*_ *<* 2*×* RMSE_*E,p*_, where RMSE_*E,p*_ = 0.0816 is the RMSE of fitting the *µ*_*p*_s of the *E*.*coli* proteome to a power law, are colored in blue while the rest in black. In Table S2, the percentages of the black data points over the total number of data points are listed, respectively, as *P*_*s*_ and *P*_*p*_. The x-axes are *n*, and the y-axes are *µ*_*s*_ (**a**) and *µ*_*p*_ (**b**) in *Å*^2^.

A typical eukaryotic protein has a high level glycosylation during past-translation modifications. However, glycosylation occurs in both prokaryotic and eukaryotic proteins. Though 50% −70% of eukaryotic proteins are possibly glycosylated, the percentage of glycosylated surface side chains of a eukaryotic protein is unknown but likely low. The *µ*s are an average over a whole surface while the power laws are an average over a set of crystal structures or models. In addition, glycosylation has not been considered in AlphaFold2 and the vast majority of the structures in **G** are not glycosylated. So it is unlikely that glycosylation could account for the large *P*_*s*_s (37.7%−57.7%) and *P*_*p*_s (29.9%−48.5%) computed with the power laws for the *E*.*coli* proteome as references (Fig. 6 and Table S2).

##### 4.4.3.2 The differences are due possibly to the lack of high quality training data for eukaryotic proteomes

As discussed in the last section, the percentages of disordered residues and glycosylation are not large enough to account for the differences in SES between the prokaryotic and eukaryotic proteomes. At present we think that the differences are attributed at least partially to the lack of high quality training data for eukaryotic proteomes.

### 4.5 Possible limitations

SES is computed by modeling both the probe and protein atoms as hard spheres. This means that SES itself is only an approximation to the physical surface of a protein. Consequently, both the power laws and their intersections are only an approximation to a likely more complicated physical relationship between protein surface and molecular size.

The surface of a soluble protein is inherently flexible in solution [2] while crystal structure is static in nature. However, if one assumes that a crystal structure freezes one conformation out of an ensemble of conformations in solution, then by ergodic hypothesis the SES geometric properties described in this paper should have real physical meanings since they all are averages over a set of structures or models with each of the individual properties being an average over a whole surface. In addition, though we have not quantified the contribution to the SES geometric properties of the crystal packing for the structures in **G** and **M**, such contributions, if exist, should be minor (section S8 and Fig. S36).

## Supporting information

PDBIDs of the crystal structures

## Supporting Information

Section S1 describes the increase of the number of residues *n*_*r*_ with the number of atoms *n* for **G**, and the best-fitted linear equation between *n*_*r*_ and *n*. Section S2 presents the differences in *µ*s between **S** and **G**. Section S3 describes the changes of the *µ*s with *n*, the power laws and point of intersection for **M**. Section S4 describes the increases with *n* of the *n*_*s*_s for **G**, the *E*.*coli* and human proteomes. Section S5 presents the *τ* s for the crystal structures and AlphaFold proteomes. Section S6 describes the SES of the transmembrane region of a membrane protein. Section S7 presents the changes of the *µ*s with both *n*_*r*_ and the number of surface atoms *n*_*s*_ for **G**. Section S8 describes the SESs of the individual chains of the multimers in **G**. Section S9 presents the *R*_*square*_s, *N*_*sp*_s and *τ* s for the crystal structures and AlphaFold models. Section S10 presents the differences in SES between the *E*.*coli* proteome and the other twenty six proteomes. Section S11 describes the changes with *n* of the *µ*s and *n*_*s*_s of the 350, 974 AlphaFold models for twenty seven proteomes. Section S12 lists the biological names of the twenty seven species and their abbreviations. Section S13 describes an example of a protein-protein interface formed by crystal packing.

### S1: The increase of *n*_*r*_ with *n* for G

As shown in Fig. S1, the increase of *n*_*r*_ with *n* for **G** could be fitted to a linear equation with a MATLAB *R*_*square*_ = 0.9982. Because some structures in **G** have missing atoms, it is more accurate to use this equation to compute *n*_*r*_ from *n* than to use directly the fitted power laws for the variations of the *µ*s with *n*_*r*_ (Fig. S6).

**Figure S1:**
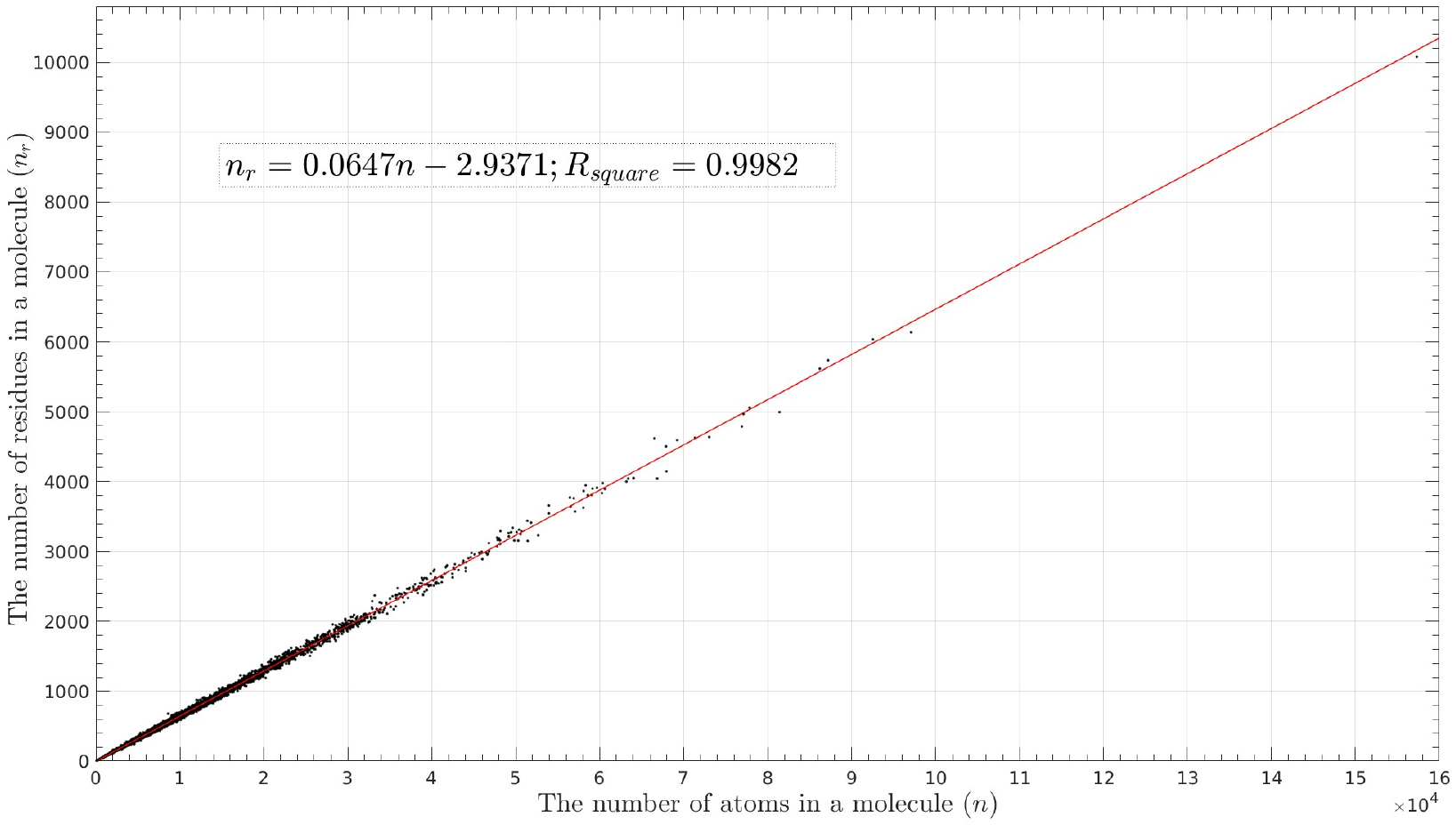
The increase of *n*_*r*_ with *n* for G. The insert is the MATLAB fitted linear equation, *n*_*r*_ = 0.0647*n*−2.9371, with a *R*_*square*_ = 0.9982. As described in the main text, the equation is used to compute *N*_*sp*_ from *n*_*sp*_. The number of atoms per residue, **16**, estimated from its slope 0.0647, is used to compute *N*_*sp*_ from *n*_*sp*_ for the twenty seven AlphaFold proteomes (Table S1). The x-axis is *n*, and y-axis *n*_*s*_.

### S2: The differences in *µ*s between S and G

As shown in Fig. S2, the variations with *n* of the *µ*_*s*_s and *µ*_*p*_s of **S** and **G**, and their power laws are very close in the sense that *<* 6% of the *µ*_*s*_s of **S** has a Δ*µ*_*s*_ *>* 2 *×* RMSE_*G,s*_, and *<* 3% of the *µ*_*p*_s of **S** has a Δ*µ*_*p*_ *>* 2 *×* RMSE_*G,p*_.

**Figure S2:**
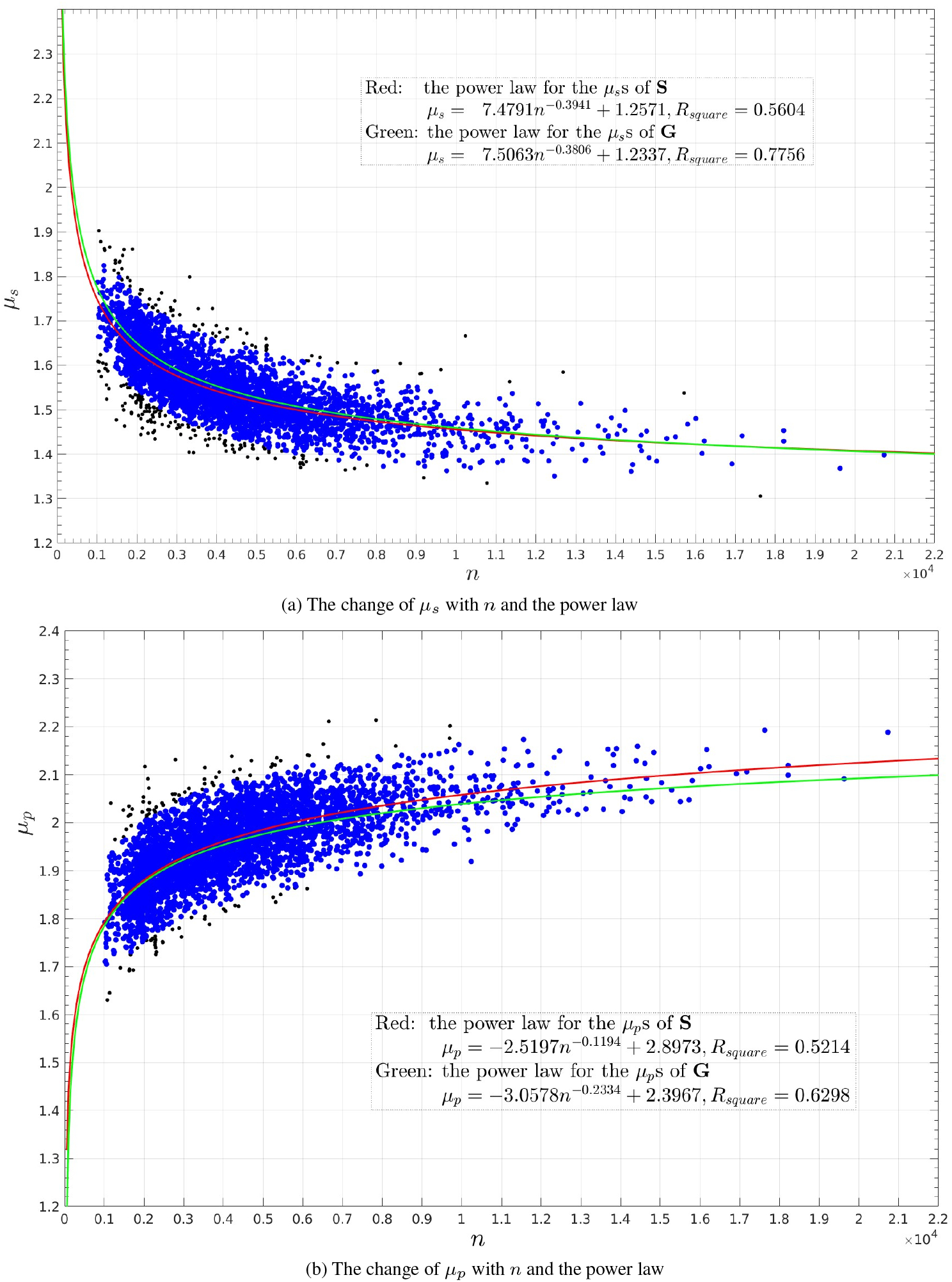
The changes with *n* of the *µ*s for S and the comparisons with those for G. Figs. (**a**) and (**b**) depict, respectively, the *µ*_*s*_s and *µ*_*p*_s of **S** and the power law curves. In Fig. (**a**) the *µ*_*s*_s with Δ*µ*_*s*_ *<* 2*×* RMSE_*G,s*_, where RMSE_*G,s*_ = 0.0532 is the RMSE of fitting the *µ*_*s*_s for **G** to a power law, and in Fig. (**b**) the *µ*_*p*_s with Δ*µ*_*p*_ *<* 2 *×* RMSE_*G,p*_, where RMSE_*G,p*_ = 0.0631 is the RMSE of fitting the *µ*_*p*_s for **G** to a power law, are colored in blue and the rest in black. There are only 229 structures in **S** with a Δ*µ*_*s*_ *>* 2*×* RMSE_*G,s*_, and 103 with a Δ*µ*_*p*_ *>* 2 *×* RMSE_*G,p*_ (Table 1 of the main text). The inserts list the power laws, *R*_*square*_s and coloring schemes. The x-axes are *n*, and the y-axes are *µ*_*s*_ (**a**) and *µ*_*p*_ (**b**) in *Å*^2^.

### S3: The changes of the *µ*s with *n*, the power laws and point of intersection for M

As shown in Fig. S3, the changes with *n* of the *µ*s for **M** and the power laws are close to those for **G**.

**Figure S3:**
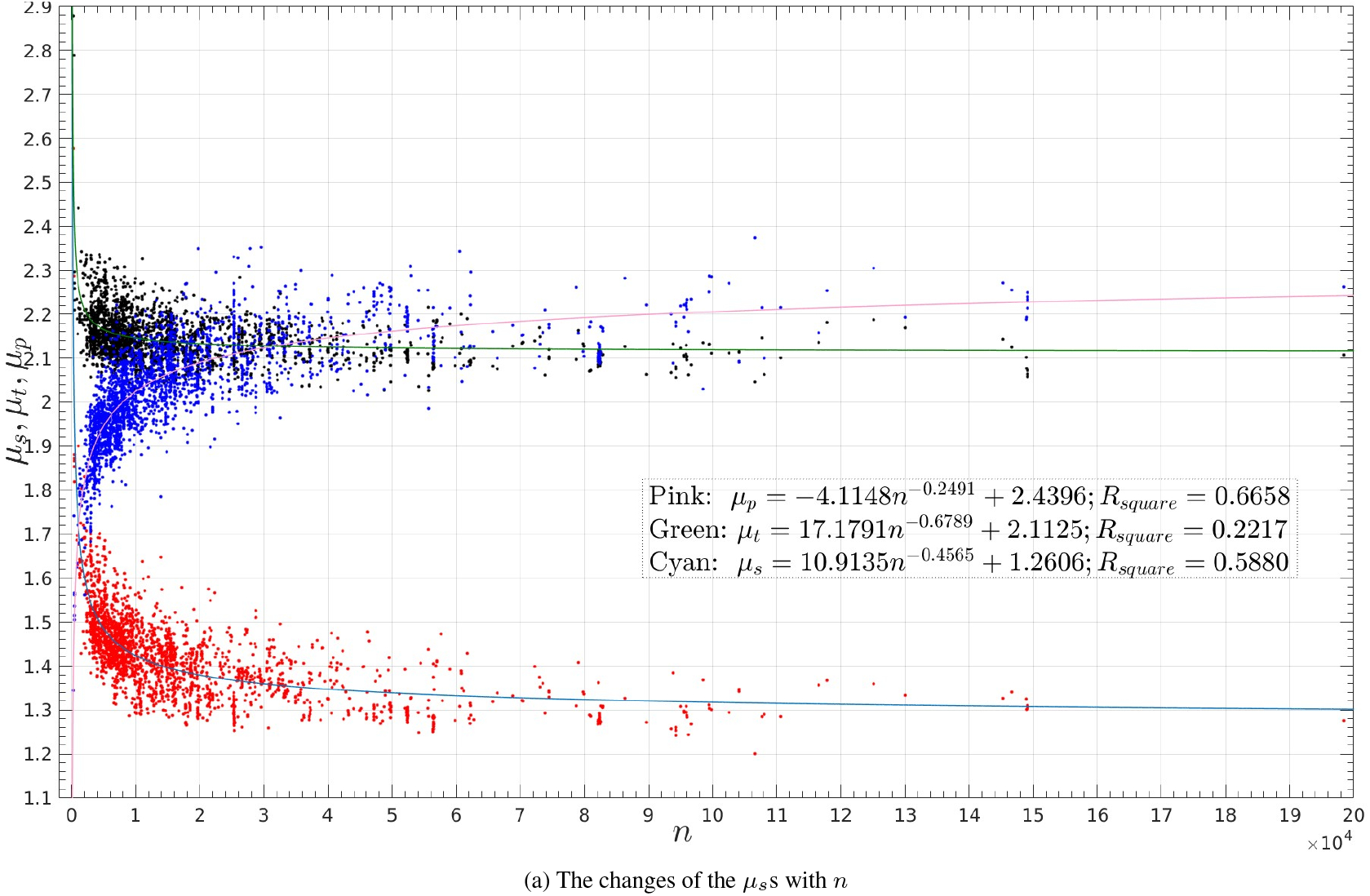
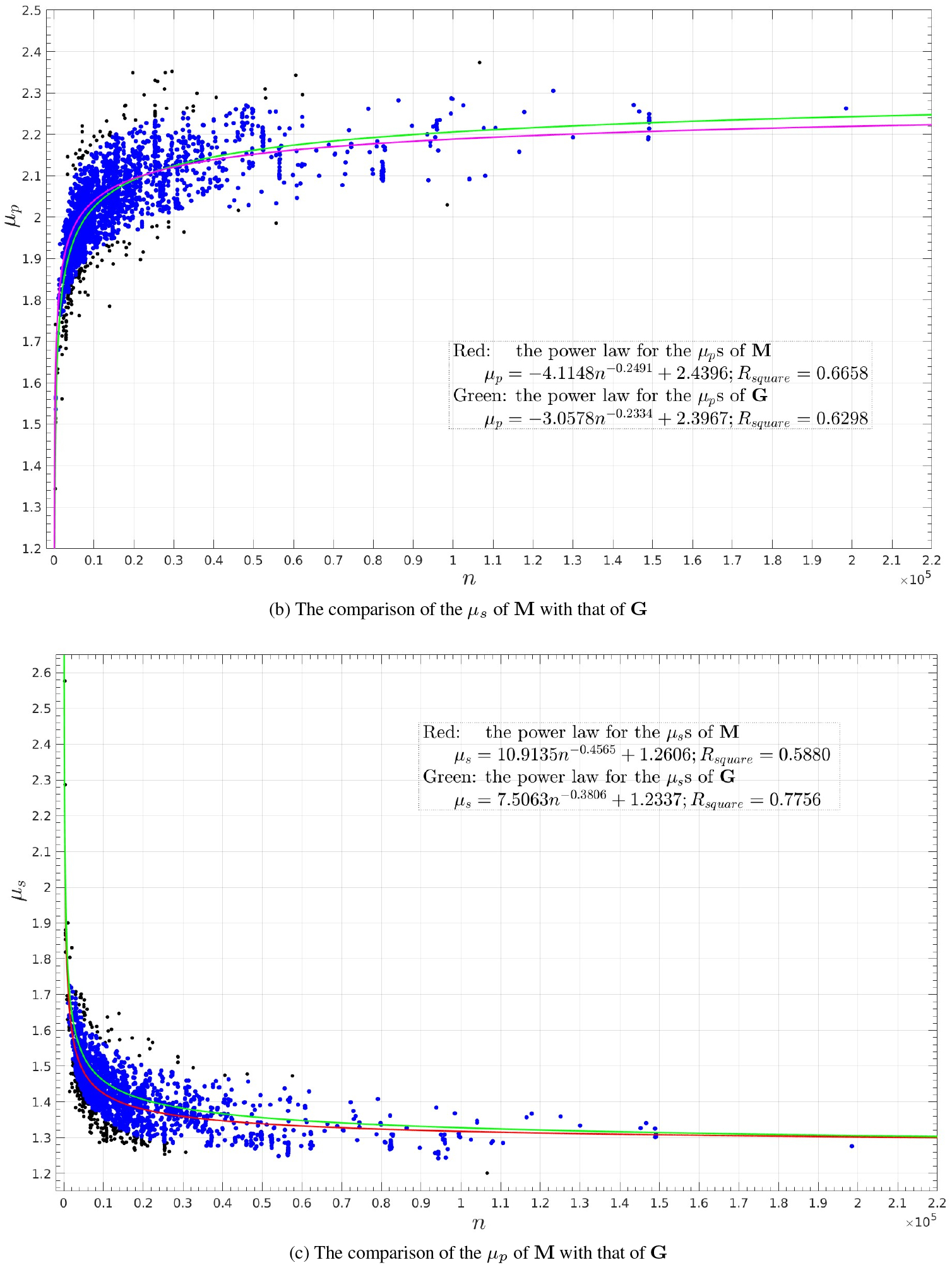
The changes with *n* of the *µ*s for M and the comparisons with those for G. Fig. (**a**) shows the *µ*_*s*_s, *µ*_*t*_s and *µ*_*p*_s of **M** colored, respectively, in red, black and blue with the power law curves colored, respectively, in cyan, green and pink. Figs. (**b**) and (**c**) depict, respectively, the *µ*_*s*_s and *µ*_*p*_s of **M** and the power law curves. In Fig. (**b**) the *µ*_*s*_s with Δ*µ*_*s*_ *<* 2*×* RMSE_*G,s*_, where RMSE_*G,s*_ = 0.0532 is the RMSE of fitting the *µ*_*s*_s for **G** to a power law, and in Fig. (**c**) the *µ*_*p*_s with Δ*µ*_*p*_ *<* 2 *×* RMSE_*G,p*_, where RMSE_*G,p*_ = 0.0631 is the RMSE of fitting the *µ*_*p*_s for **G** to a power law, are colored in blue and the rest in black. There are only 303 structures in **S** with a Δ*µ*_*s*_ *>* 2 *×* RMSE_*G,s*_, and 159 with a Δ*µ*_*p*_ *>* 2 *×* RMSE_*G,p*_ (Table 1 of the main text). The inserts list the power laws, *R*_*square*_s and coloring schemes. The x-axes are *n*, and y-axes are *µ*_*s*_, *µ*_*t*_ and *µ*_*p*_ (**a**), *µ*_*s*_ (**b**) and *µ*_*p*_ (**c**) in *Å*^2^.

### S4: The increases with *n* of the *n*_*s*_s for G, the *E*.*coli* and human proteomes

For **G**, the number of surface atoms *n*_*s*_ increases almost linearly with molecular size *n* with a *τ* = 0.3770 (Fig. S4a). The *τ* s for the *E*.*coli* proteome and human proteome are, respectively, 20.6% and 78.8% larger than that for **G** (Figs. S4b and S4c). Strangely, as shown in Fig. S4c and Figs. S22d–S35d, the changes with *n* of the *n*_*s*_s for the fourteen eukaryotic proteomes look artificial with the range of *n*_*s*_ for approximately the same *n* also increases linearly with *n*.

**Figure S4:**
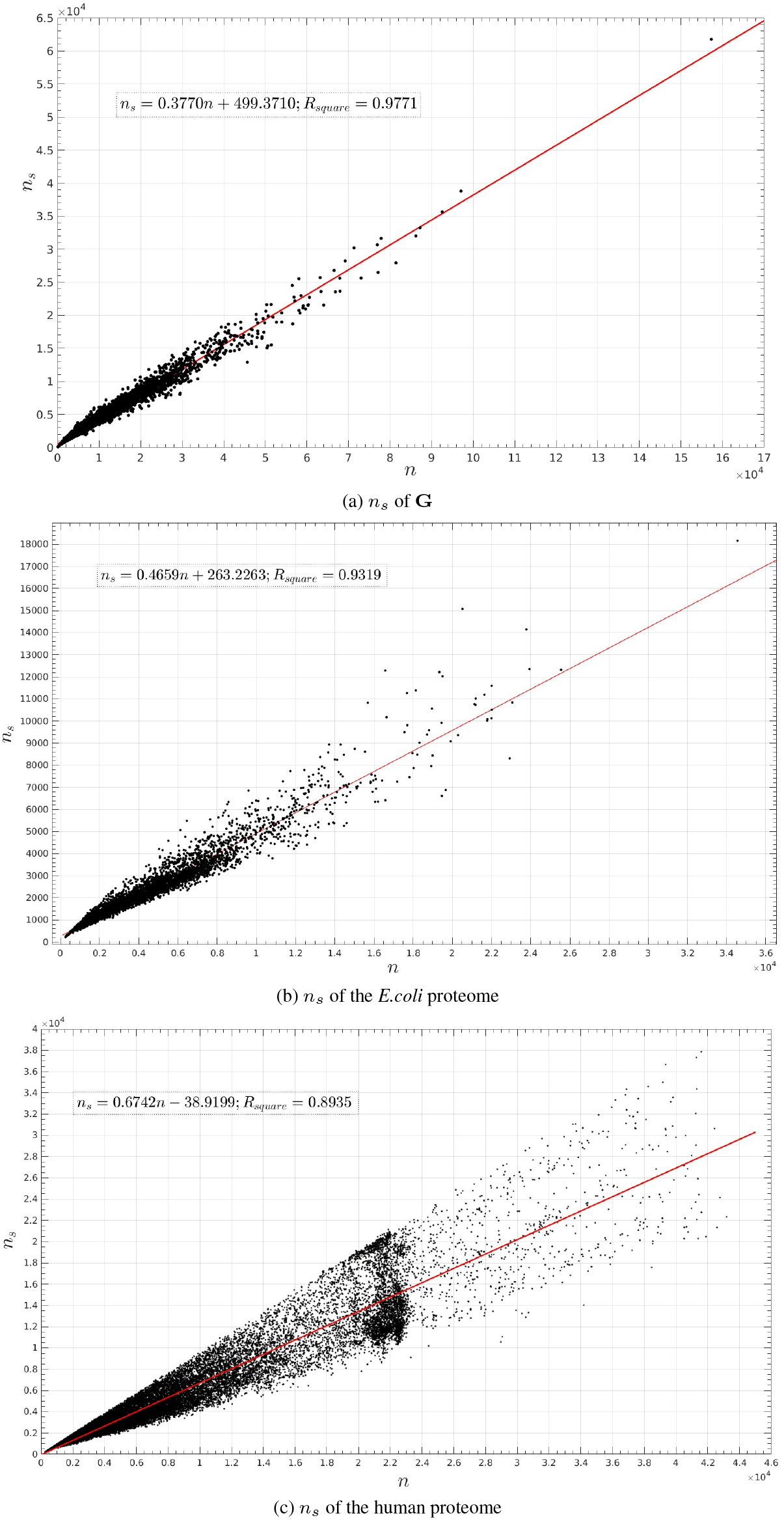
The increases of the *n*_*s*_s with *n* for G, and the *E*.*coli* and human proteomes. The fitted lines are depicted in red. The inserts list the linear equations and their *R*_*square*_s. As shown in (**c**) the increase of *n*_*s*_ with *n* looks artificial. The x-axes are *n*, and y-axes *n*_*s*_.

### S5: The *τ* s for the crystal structures and AlphaFold proteomes

As shown in Fig. S5, the *τ* s for the three sets of crystal structures are all smaller than any of the *τ* s for the twenty seven proteomes.

**Figure S5:**
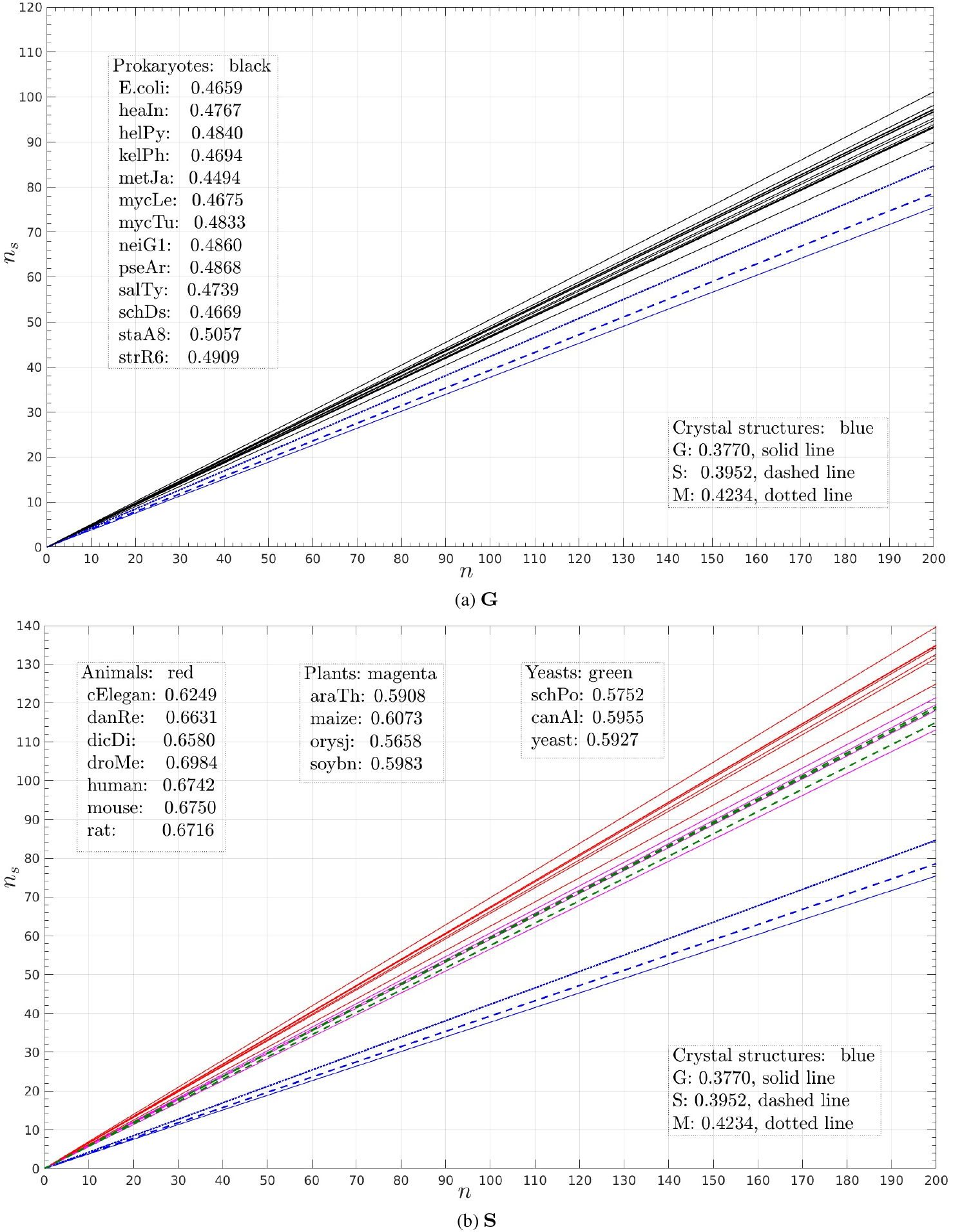
The *τ*s for the crystal structures and AlphaFold proteomes. Figs. (**a**) and (**b**) depict respectively the *τ* s for the thirteen prokaryotic proteomes and fourteen eukaryotic ones. The *τ*s for **G, S** and **M** are shown in both (**a**) and b(**b**) for comparison. The inserts list the *τ* values for **G, S** and **M** and the twenty seven proteomes. Please refers to Table S3 for their biological full names.

### S6: The SES of the transmembrane region of a membrane protein

As shown in Fig. S6, the transmembrane region of a typical membrane protein is dominated by carbon and proton atoms; and the atoms with a partial charge differing largely from zero, if there, generally having a small SAS area.

**Figure S6:**
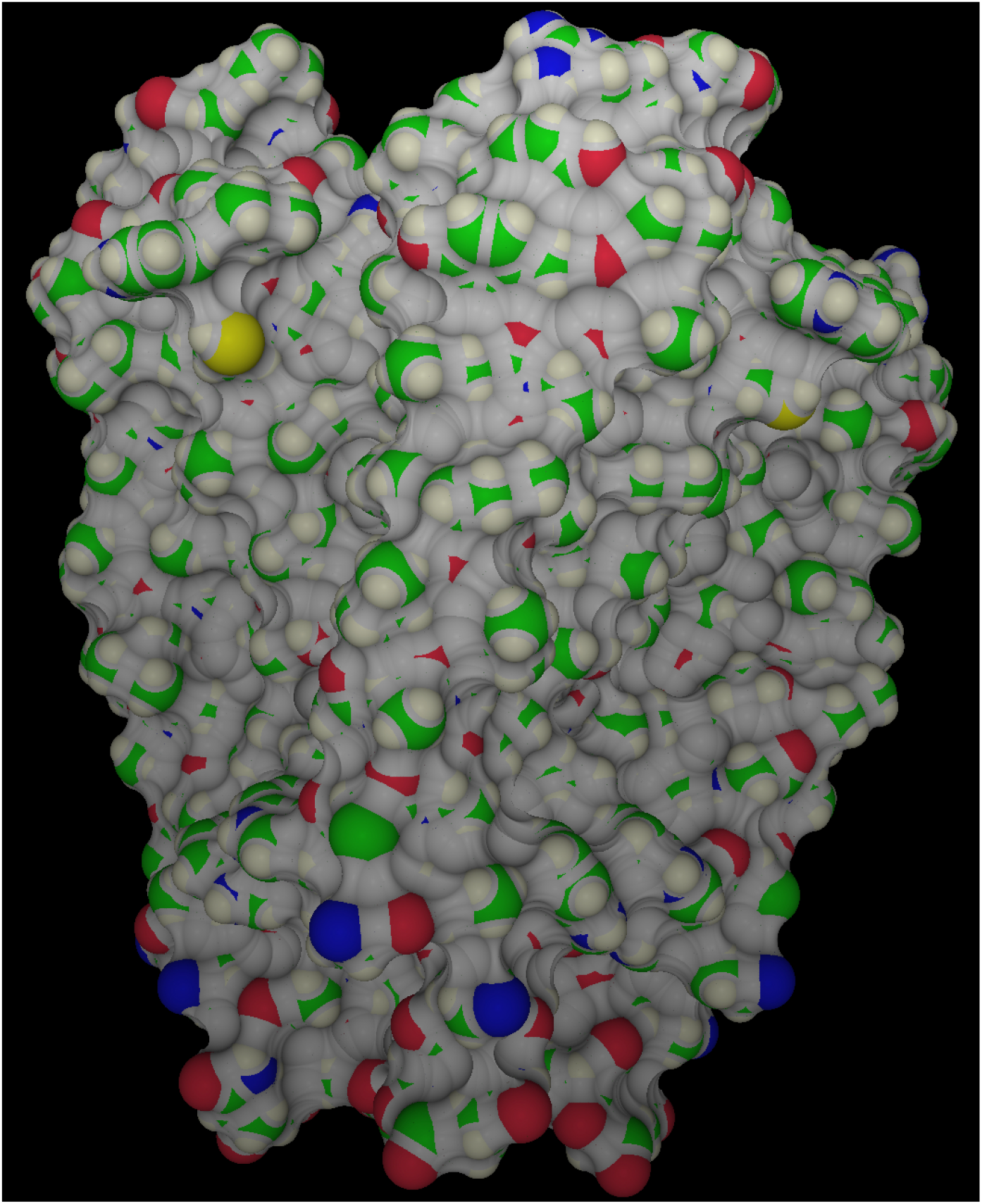
The SES of the transmembrane region of a membrane protein (1BL8). The SAA polygons for the C, N, O and H atoms of a potassium channel are colored respectively in green, blue, red and misty rose while the rest of the SES in gray. The figure is prepared by our program SESA which is freely available at https://github.com/wlincong/SESexact.

### S7: The changes with *n*_*r*_ and with *n*_*s*_ of the *µ*s of G

Fig. S7 depicts the changes of *µ*_*s*_, *µ*_*t*_ and *µ*_*p*_ with *n*_*r*_ and *n*_*s*_. As shown in the figure, compared with the *R*_*square*_s for fitting to a power law, the changes of *µ*s with *n*, the *R*_*square*_s for fitting the changes of *µ*s with *n*_*r*_ and *n*_*s*_ are all smaller, except for the change of *µ*_*t*_ with *n*_*s*_, which has a *R*_*square*_ slightly larger than that for fitting to a power law the change of *µ*_*t*_ with *n*.

**Figure S7:**
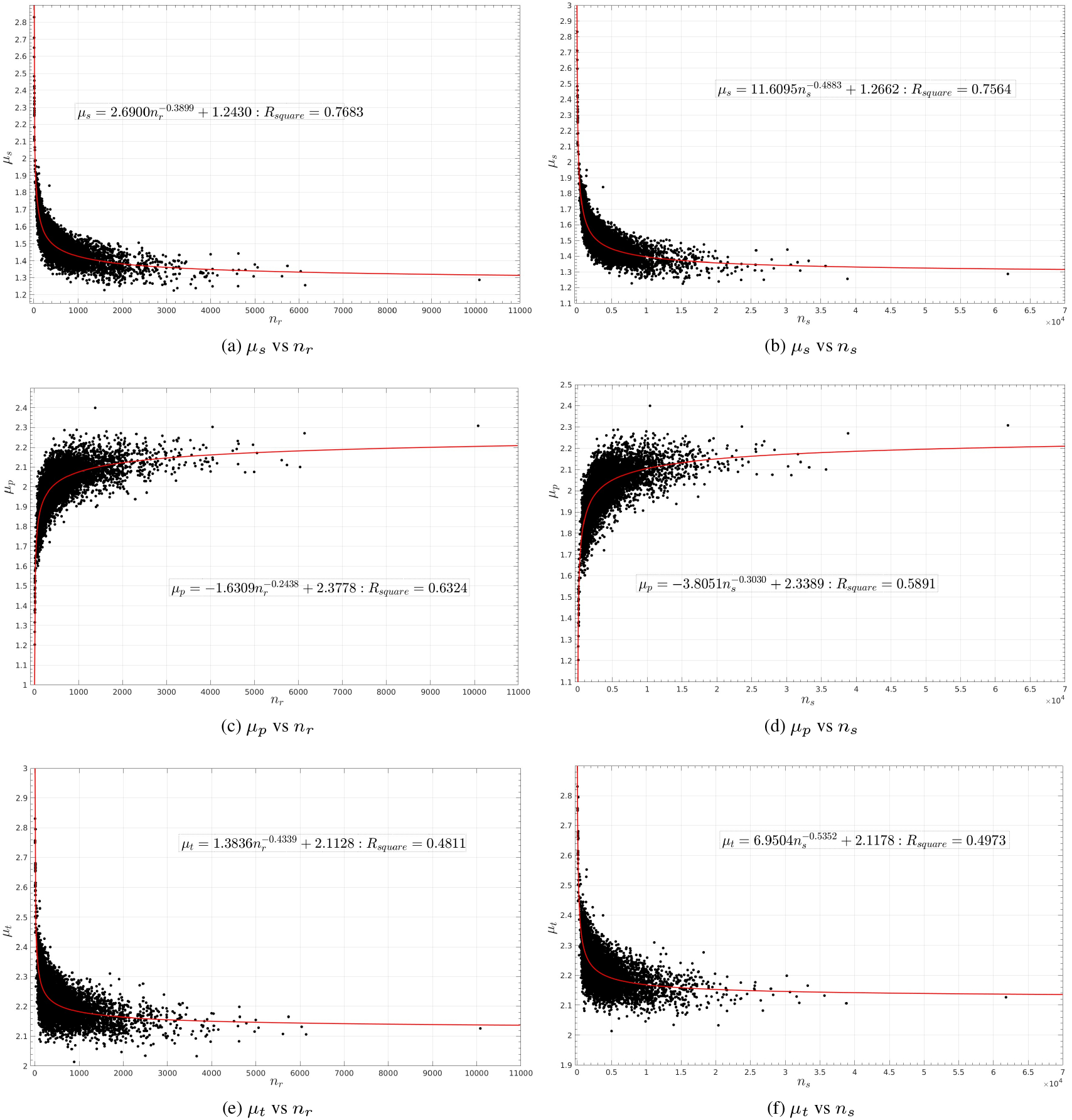
The changes of *µ*_*s*_, *µ*_*p*_ and *µ*_*t*_ with *n*_*r*_ and *n*_*s*_ for G. Figs. (**a, c, e**) show the changes of *µ*s with *n*_*r*_ while Figs. (**b, d, f**) the changes with *n*_*s*_. The insets list the power laws and *R*_*square*_s. The power law curves are depicted in red. The point of intersection of the power laws for *µ*_*s*_ and *µ*_*p*_ is *n*_*r*_ = 62, which is slightly larger than 60 residues computed using the linear equation in Fig. S1. The point of intersection of the power laws for *µ*_*t*_ and *µ*_*p*_ is *n*_*r*_ = 3, 241. The existence of missing atoms in many structures means that these two *n*_*r*_ values are upper limits.

### S8: The SESs of the individual chains of the multimers in G

We extract a set **G**_*c*_ of 20, 050 chains from all the multimers in **G** and compute their SESs individually. **G**_*c*_ includes no monomeric proteins. Among them the smallest chain is chain H of 4TY9 with only 3 residues and 60 atoms, and the largest one is chain A of 3EGW with 1, 244 residues and 19, 403 atoms. As shown in Fig. S8, the changes with *n* of the *µ*s and the power laws for **G**_*c*_ are almost identical to those for **G** itself. In addition, both the *µ*_*s*_s and *µ*_*p*_s of **G**_*c*_ could be fitted to a power law better than the corresponding ones for **G**. In addition, the closeness in SES of **G**_*c*_ to **G** and **S** to **G** shows that crystal packing, if exists, should have minimal impacts on the geometric properties studied in this paper.

**Figure S8:**
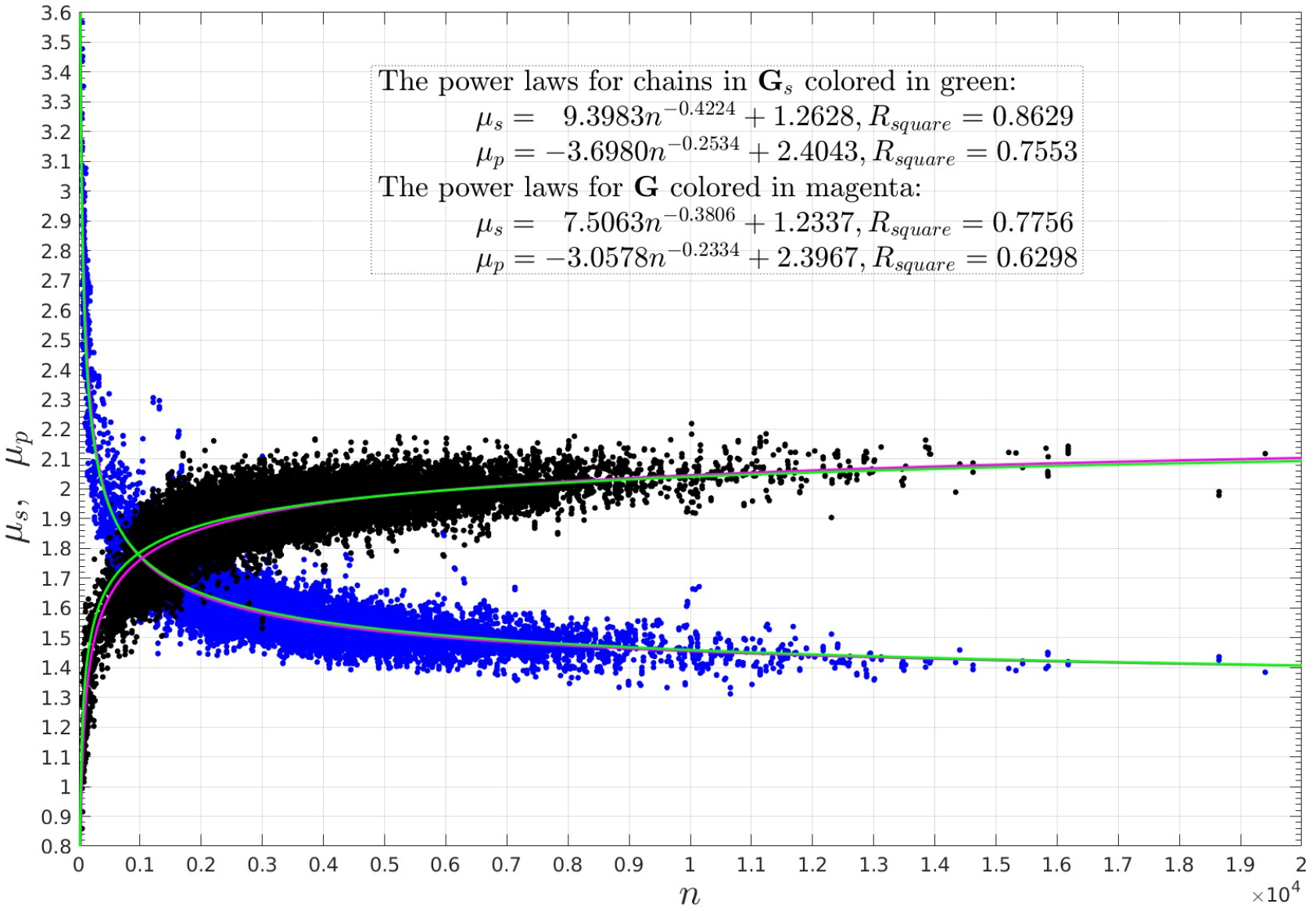
The power laws for the *µ*s of the individual chains in G. The *µ*_*s*_s and *µ*_*p*_s are depicted, respectively, in blue and black. The insert lists the power laws for the *µ*s of both **G** and **G**_*c*_, the set of **G**’s individual chains. The power laws for **G**_*c*_ and *n*_*sp*_ are almost identical to those for **G** itself. The mean and standard deviation of the list of the Δ*µ*_*s*_s (defined in the legend of Table 1 of the main text) are only −0.0054Å ^2^ and 0.0637Å ^2^, the absolute value of the mean is almost 10-fold smaller than RMSE_*G,s*_ = 0.0532. The mean and standard deviation of the list of the Δ*µ*_*p*_s are only −0.0097Å ^2^ and 0.0673Å ^2^, the absolute value of the mean is almost 7-fold smaller than RMSE_*G,p*_ = 0.0631.

### S9: The *R*_*square*_s, *N*_*sp*_s and *τ* s for the crystal structures and AlphaFold models

Table S1 lists the *R*_*square*_s, *N*_*sp*_s (points of intersection) and *τ* s for the three sets of crystal structures and the twenty seven AlphaFold proteomes.

**Table S1:** The *R*_*square*_**s**, *τ* **s and** *N*_*sp*_**s for the three sets of crystal structures and twenty seven AlphaFold proteomes**. Column 1 lists species names (Table S3). Column 2 lists the *R*_*square*_s for *µ*_*s*_ and *µ*_*p*_. Column 3 lists the slopes (*τ* s) of the fitted linear equations between *n* and *n*_*s*_. Column 4 lists the *N*_*sp*_s (computed using, 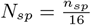,Fig. S1). Among the twenty seven proteomes, the human proteome has the largest *N*_*sp*_ = 694 and the third largest *τ* = 0.6742.

| Organism | $R_{square} (\mu_s/\mu_p)$ | $\tau$ | $N_{sp}$ | $\mu_{sp}$ |
| --- | --- | --- | --- | --- |
| **Crystal structures** |  |  |  |  |
| G | **0.776 / 0.630** | **0.3770** | **60** | **1.783** |
| S | **0.560 / 0.521** | **0.3952** | **57** | **1.775** |
| M | **0.666 / 0.588** | **0.4234** | **64** | **1.714** |
| **Prokaryotic proteomes** |  |  |  |  |
| E.coli | 0.582 / 0.571 | 0.4659 | 87 | 1.781 |
| haeIn | 0.487 / 0.501 | 0.4767 | 86 | 1.780 |
| helPy | 0.499 / 0.523 | 0.4840 | 89 | 1.779 |
| klePh | 0.629 / 0.582 | 0.4694 | 92 | 1.779 |
| metJa | 0.487 / 0.488 | 0.4494 | 75 | 1.782 |
| mycLe | 0.372 / 0.331 | 0.4675 | 110 | 1.790 |
| mycTu | 0.369 / 0.379 | 0.4833 | 113 | 1.772 |
| neiGl | 0.556 / 0.482 | 0.4860 | 99 | 1.787 |
| pseAr | 0.404 / 0.414 | 0.4868 | 93 | 1.769 |
| salTy | 0.504 / 0.457 | 0.4739 | 91 | 1.779 |
| shiDs | 0.509 / 0.503 | 0.4669 | 93 | 1.787 |
| staA8 | 0.516 / 0.546 | 0.5057 | 75 | 1.785 |
| strR6 | 0.505 / 0.503 | 0.4909 | 81 | 1.783 |
| **Yeast proteomes** |  |  |  |  |
| yeast | 0.056 / 0.083 | 0.5927 | 133 | 1.799 |
| schPo | 0.056 / 0.086 | 0.5752 | 125 | 1.793 |
| canAl | 0.036 / 0.072 | 0.5955 | 145 | 1.800 |
| **Plant proteomes** |  |  |  |  |
| araTh | 0.125 / 0.123 | 0.5908 | 246 | 1.808 |
| orySj | 0.206 / 0.172 | 0.5658 | 391 | 1.806 |
| soyBn | 0.093 / 0.081 | 0.5983 | 250 | 1.808 |
| maize | 0.104 / 0.094 | 0.6073 | 389 | 1.805 |
| **Eukaryotic proteomes** |  |  |  |  |
| celEg | 0.126 / 0.123 | 0.6249 | 156 | 1.809 |
| dicDi | 0.023 / 0.018 | 0.6631 | 208 | 1.824 |
| danRe | 0.092 / 0.085 | 0.6580 | 116 | 1.805 |
| droMe | 0.037 / 0.040 | 0.6984 | 327 | 1.804 |
| mouse | 0.028 / 0.023 | 0.6750 | 193 | 1.798 |
| rat | 0.030 / 0.025 | 0.6716 | 258 | 1.797 |
| human | **0.034 / 0.025** | **0.6742** | **694** | **1.800** |

### S10: The differences in SES between the *E*.*coli* proteome and the other twenty six proteomes

Table S2 lists the Δ*µ*_*s*_s, Δ*µ*_*p*_s and *P*_*s*_s and *P*_*p*_s for the twenty six proteomes computed with the power laws for the *E*.*coli* proteome as references.

**Table S2:** The differences in SES between the *E*.*coli* proteome and other twenty six proteomes. Column 1 lists organism names (Table S3). The differences in *µ*_*s*_ and *µ*_*p*_ of model *i* of a proteome from the *µ*_*s*_s and *µ*_*p*_s of the *E*.*coli* proteome are quantified as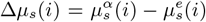 and 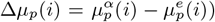, where 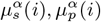 are the *µ*_*s*_, *µ*_*p*_ values of model *i*, and 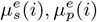 are computed using, respectively, the power laws for the *µ*_*s*_s and *µ*_*p*_s of the *E*.*coli* proteome, with *n* in Eq. (5) being the number of atoms of model *i*. Column 2 lists the mean and standard deviation of the list of the Δ*µ*_*s*_s for a proteome. Column 3 lists the mean and standard deviation of the list of the Δ*µ*_*p*_s for a proteome. Columns 4 and 5 list, respectively, the *P*_*s*_s and *P*_*p*_s for the twenty-six proteomes, and Fig. 6 of the main text visualizes the *P*_*s*_ and *P*_*p*_ for the human proteome. The *P*_*s*_ and *P*_*p*_ have the same meanings as those in Table 1 of the main text except that they are calculated with the RMSE_*E,s*_ = 0.1002 for fitting to a power law the *µ*_*s*_s of the *E*.*coli* proteome, and RMSE_*E,p*_ = 0.0816 for fitting to a power law the *µ*_*p*_s of the *E*.*coli* proteome. As shown in the last row, the *µ*s, *P*_*s*_ and *P*_*e*_ for the human proteome are the furthest from those for the *E*.*coli* proteome.

| Organism | $\Delta\mu_s$ (mean / std) ($\text{\AA}^2$) | $\Delta\mu_p$ (mean / std) ($\text{\AA}^2$) | $P_s$ (%) | $P_p$ (%) |
| --- | --- | --- | --- | --- |
| **Prokaryotic proteomes** |  |  |  |  |
| haeIn | -0.0001 / 0.0992 | 0.0030 / 0.0804 | 5.1 | 4.9 |
| helPy | 0.0105 / 0.1117 | -0.0057 / 0.0885 | 7.1 | 6.1 |
| klePh | 0.0029 / 0.1027 | -0.0082 / 0.0857 | 4.9 | 6.1 |
| **metja** | **-0.0207 / 0.0859** | **0.0199 / 0.0724** | **2.4** | **4.3** |
| mycLe | 0.0458 / 0.1352 | -0.0300 / 0.1089 | 12.9 | 12.7 |
| mycTu | 0.0355 / 0.1382 | -0.0404 / 0.1089 | 12.6 | 14.5 |
| neiG1 | 0.0219 / 0.1252 | -0.0130 / 0.1015 | 9.5 | 10.7 |
| pseAr | 0.0036 / 0.1112 | -0.0184 / 0.0854 | 6.0 | 5.8 |
| salTy | 0.0064 / 0.1036 | -0.0036 / 0.0843 | 5.7 | 5.8 |
| shiDs | 0.0076 / 0.1057 | -0.0064 / 0.0872 | 5.6 | 6.5 |
| staA8 | 0.0014 / 0.1135 | 0.0147 / 0.0911 | 6.0 | 5.8 |
| strR6 | 0.0025 / 0.1008 | 0.0069 / 0.0836 | 5.2 | 5.5 |
| **Yeast proteomes** |  |  |  |  |
| schPo | 0.1603 / 0.2009 | -0.0914 / 0.1617 | 34.8 | 28.7 |
| yeast | 0.1859 / 0.2104 | -0.1058 / 0.1669 | 39.7 | 32.9 |
| canAl | 0.1936 / 0.2159 | -0.1053 / 0.1668 | 40.9 | 31.5 |
| **Plant proteomes** |  |  |  |  |
| araTh | 0.2074 / 0.2100 | -0.1188 / 0.1709 | 43.1 | 36.3 |
| soyBn | 0.2076 / 0.2136 | -0.1174 / 0.1775 | 43.3 | 36.6 |
| maize | 0.2423 / 0.2155 | -0.1512 / 0.1778 | 51.6 | 43.7 |
| orysj | 0.2548 / 0.2245 | -0.1527 / 0.1857 | 53.1 | 47.9 |
| **Eukaryotic proteomes** |  |  |  |  |
| celEg | 0.1740 / 0.2089 | -0.0916 / 0.1652 | 37.7 | 29.9 |
| dicDi | 0.2119 / 0.2182 | -0.0938 / 0.1841 | 46.0 | 33.6 |
| danRe | 0.2386 / 0.2196 | -0.1497 / 0.1883 | 50.2 | 40.2 |
| droMe | 0.2445 / 0.2380 | -0.1573 / 0.1973 | 49.8 | 41.9 |
| mouse | 0.2370 / 0.2330 | -0.1617 / 0.1934 | 50.1 | 43.1 |
| rat | 0.2329 / 0.2312 | -0.1593 / 0.1918 | 49.5 | 42.7 |
| **human** | **0.2786 / 0.2330** | **-0.1935 / 0.2050** | **57.7** | **48.5** |

### S11: The changes with *n* of the *µ*s and the power laws, and the *n*_*s*_s and the linear equations for the twenty seven AlphaFold proteomes

The Figs. S9 to S35 depict the changes of *µ*_*s*_, *µ*_*p*_, *µ*_*t*_ and *n*_*s*_ with respect to *n* for the twenty seven proteomes in the order of thirteen prokaryotic (Figs. S9–S21), three yeast (Figs. S22–S24), four plant (Figs. (S25–S28)) and seven animal (Figs. S29–S35) proteomes.

#### S11.1: The thirteen prokaryotic proteomes

**Figure S9:**
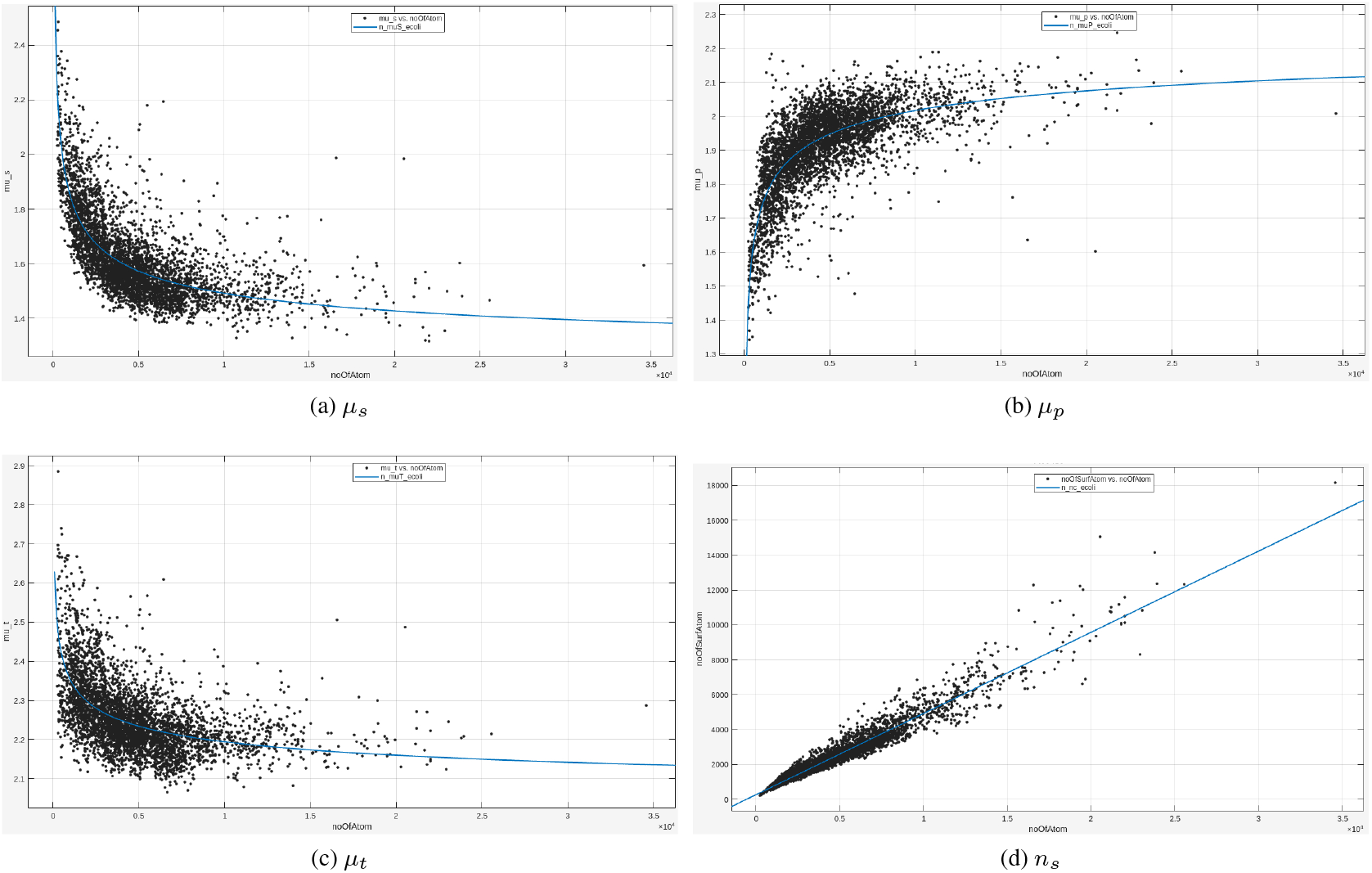
The changes with *n* of the *µ*_*s*_, *µ*_*p*_, *µ*_*t*_ **and** *n*_*s*_ **for the *Escherichia coli* proteome**.

**Figure S10:**
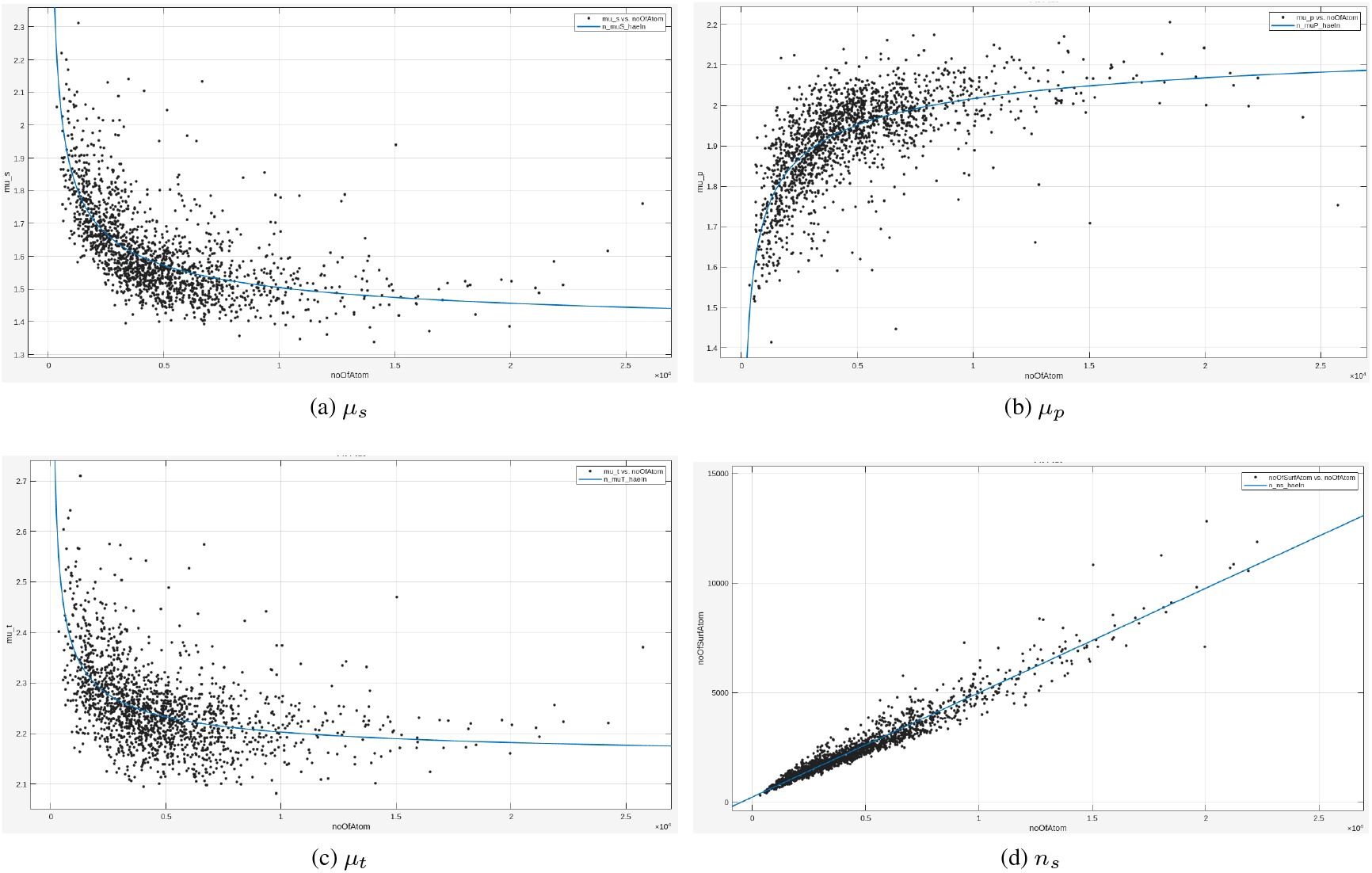
The changes with *n* of the *µ*_*s*_, *µ*_*p*_, *µ*_*t*_**s and** *n*_*s*_**s for the *Haemophilus influenzae* proteome**.

**Figure S11:**
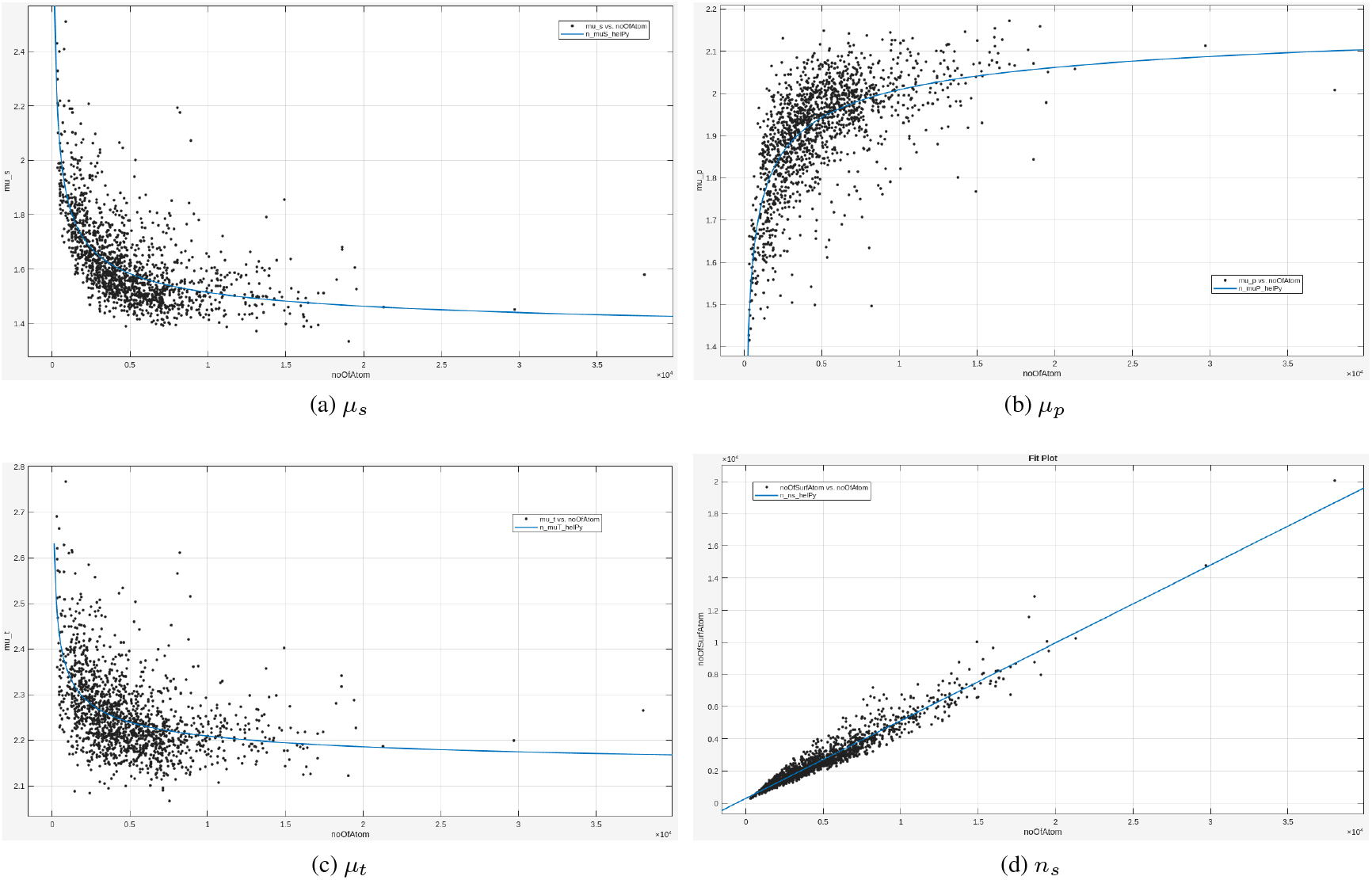
The changes with *n* of the *µ*_*s*_, *µ*_*p*_, *µ*_*t*_**s and** *n*_*s*_**s for the *Helicobacter pylori* proteome**.

**Figure S12:**
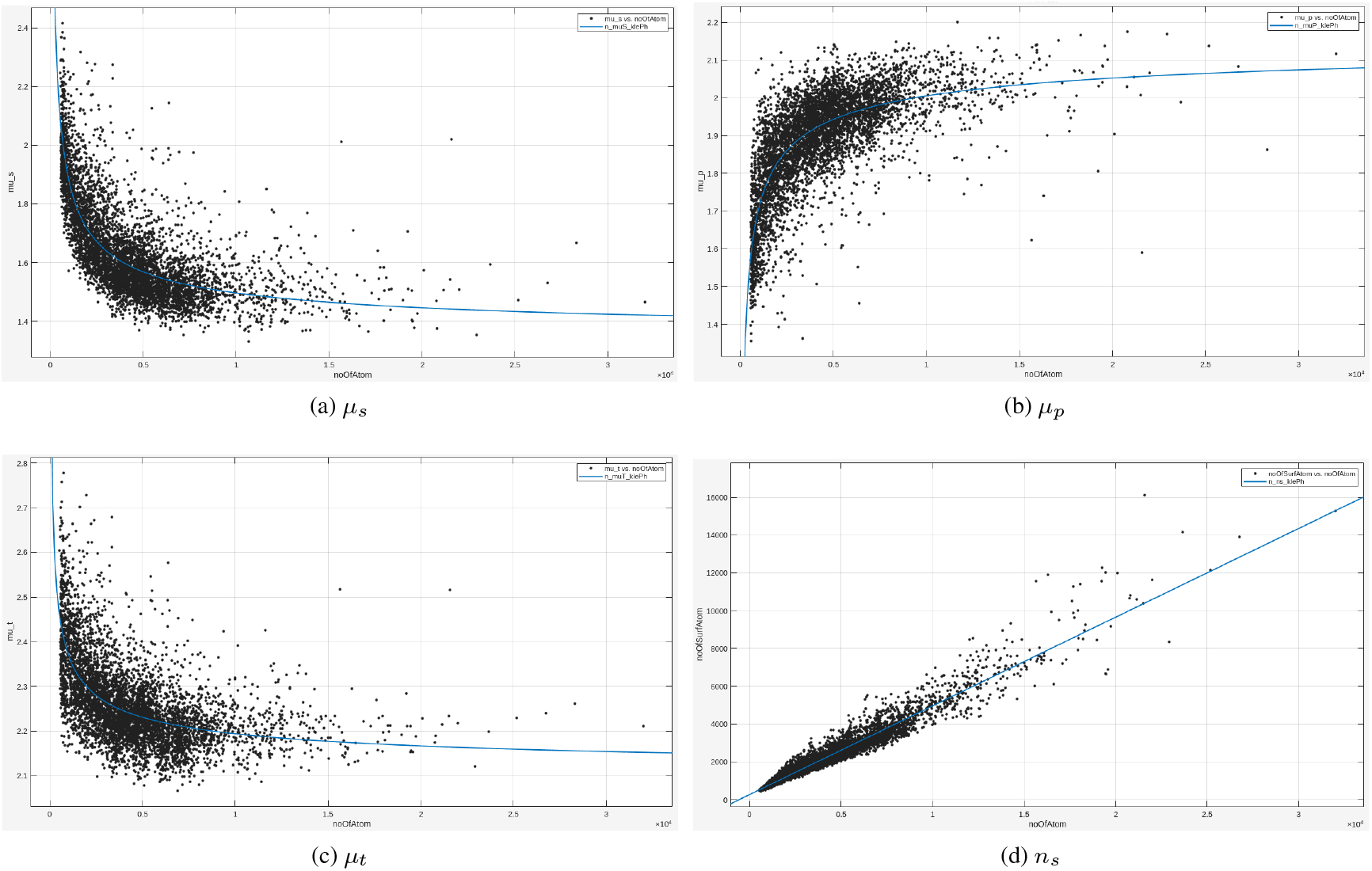
The changes with *n* of the *µ*_*s*_, *µ*_*p*_, *µ*_*t*_**s and** *n*_*s*_**s for the *Klebsiella pneumoniae* proteome**.

**Figure S13:**
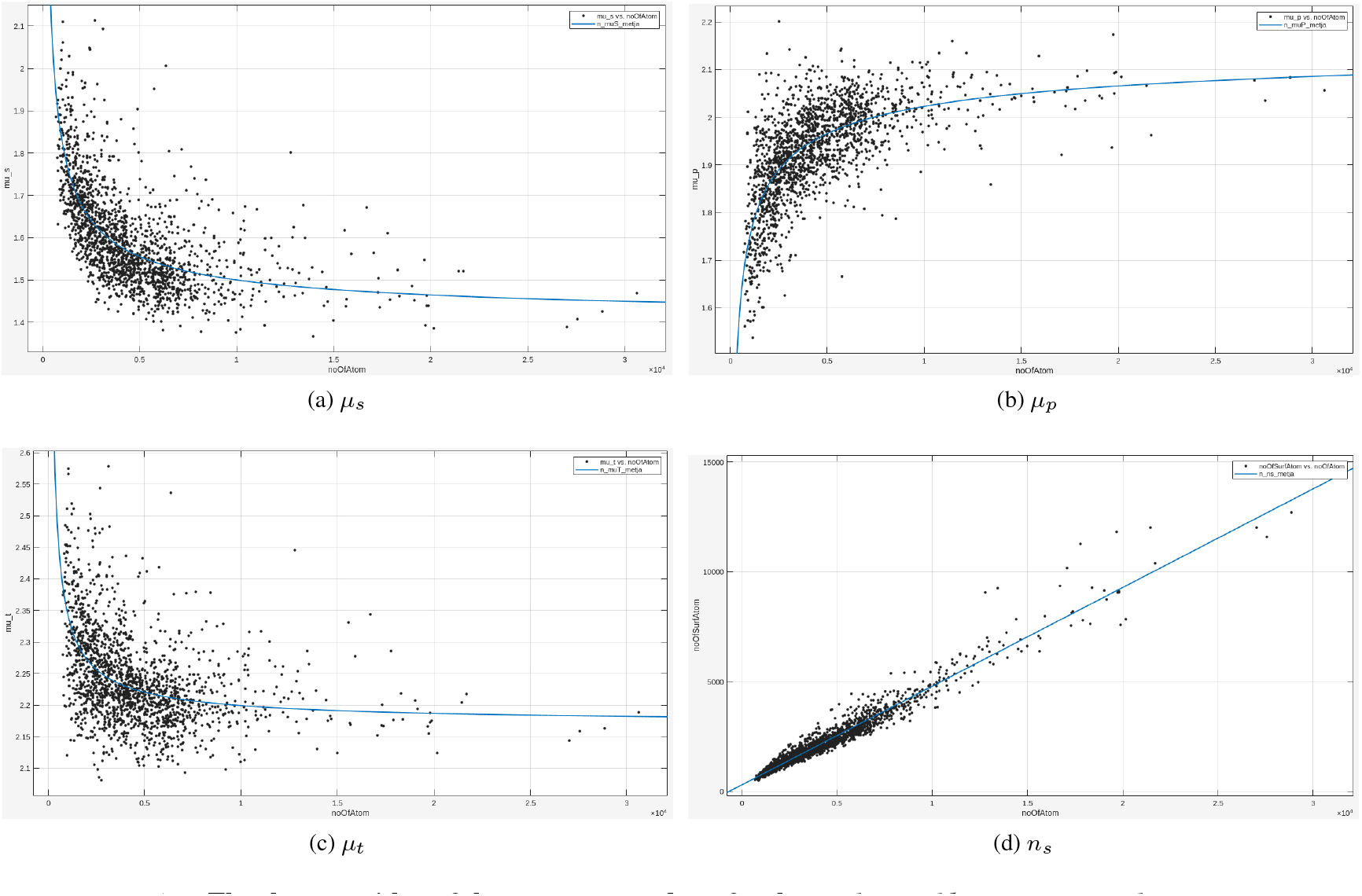
The changes with *n* of the *µ*_*s*_, *µ*_*p*_, *µ*_*t*_**s and** *n*_*s*_**s for the *Methanocaldococcus jannaschii* proteome**.

**Figure S14:**
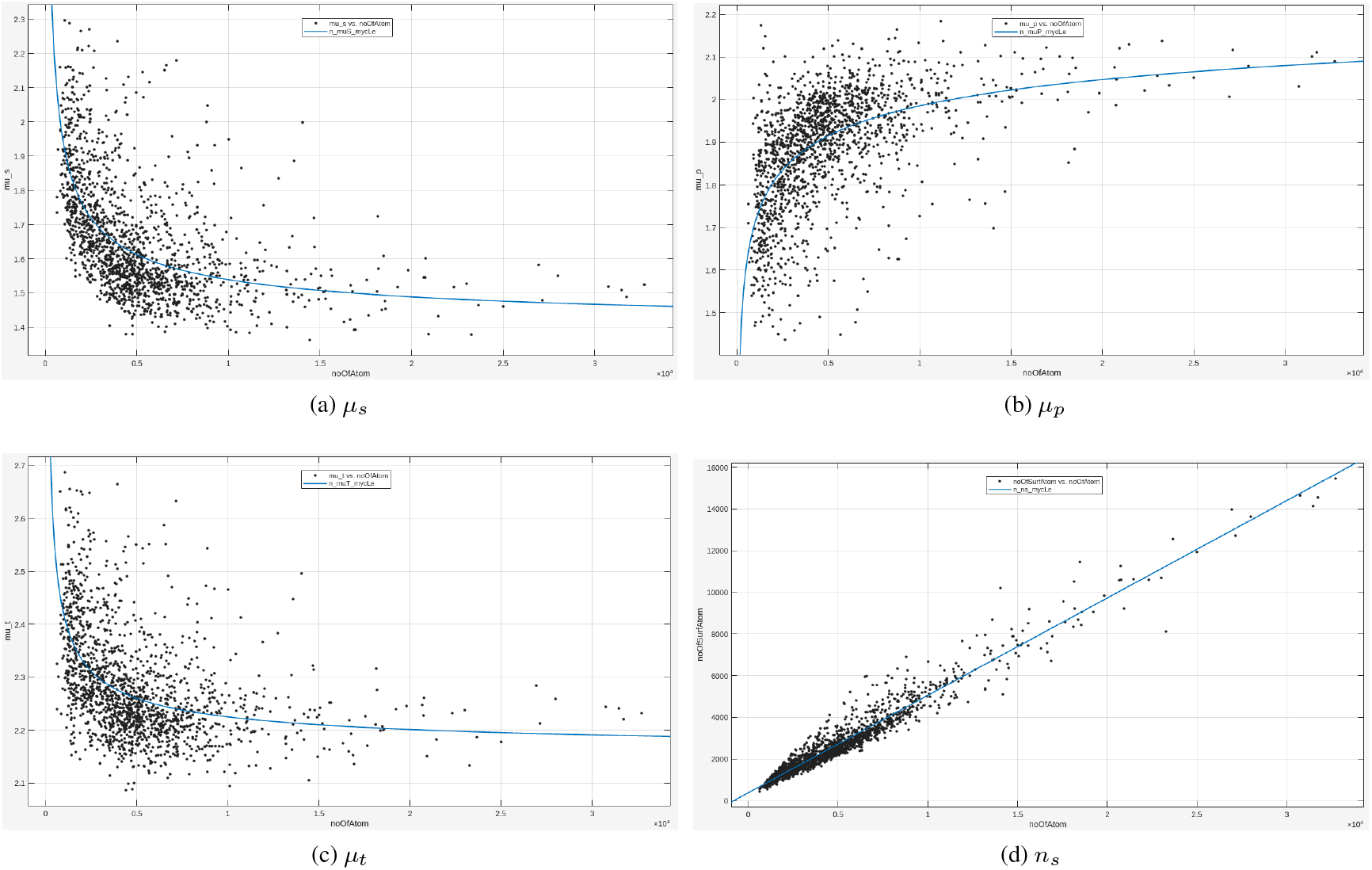
The changes with *n* of the *µ*_*s*_, *µ*_*p*_, *µ*_*t*_**s and** *n*_*s*_**s for the *Mycobacterium leprae* proteome**.

**Figure S15:**
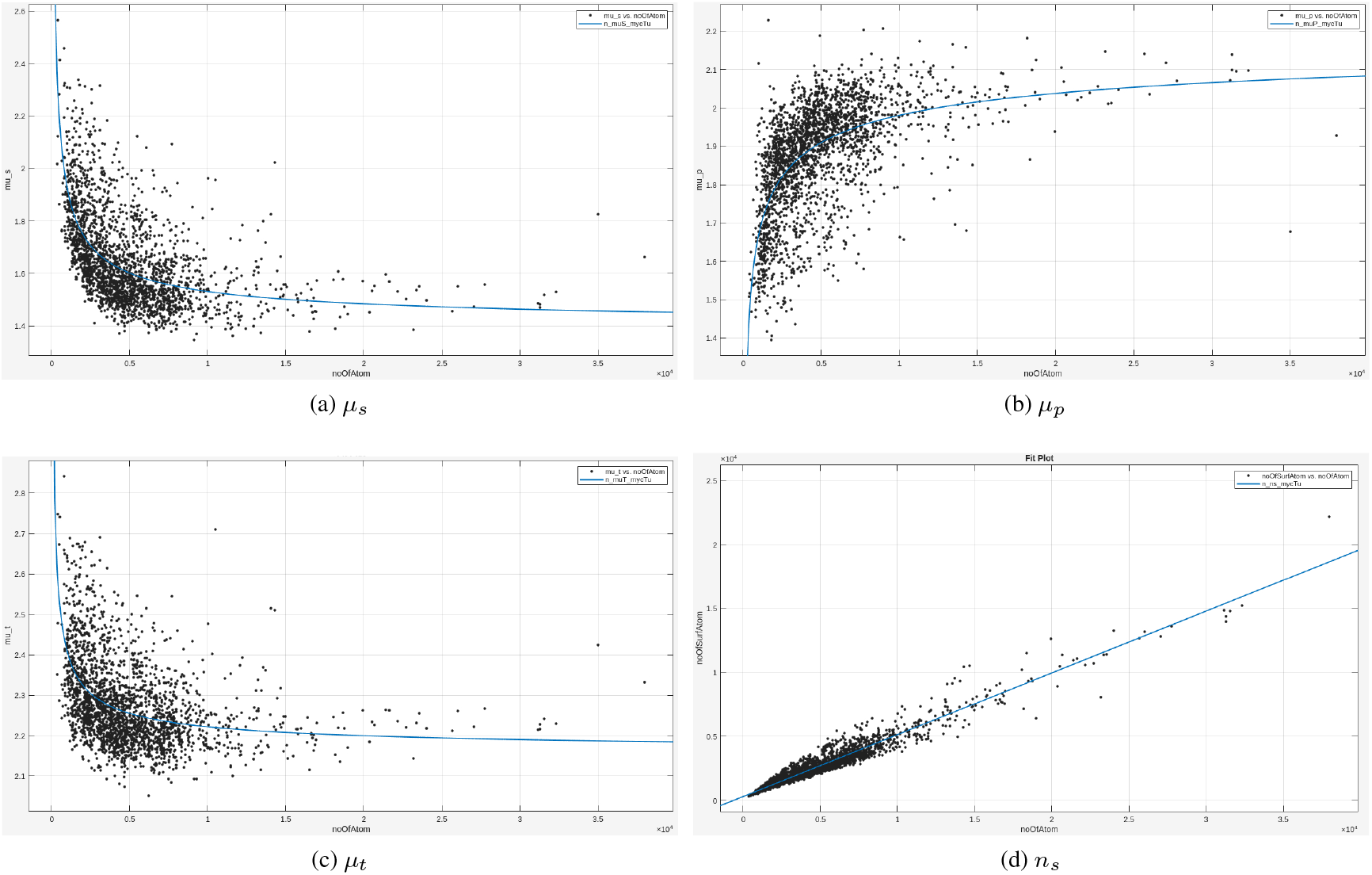
The changes with *n* of the *µ*_*s*_, *µ*_*p*_, *µ*_*t*_**s and** *n*_*s*_**s for the *Mycobacterium tuberculosis* proteome**.

**Figure S16:**
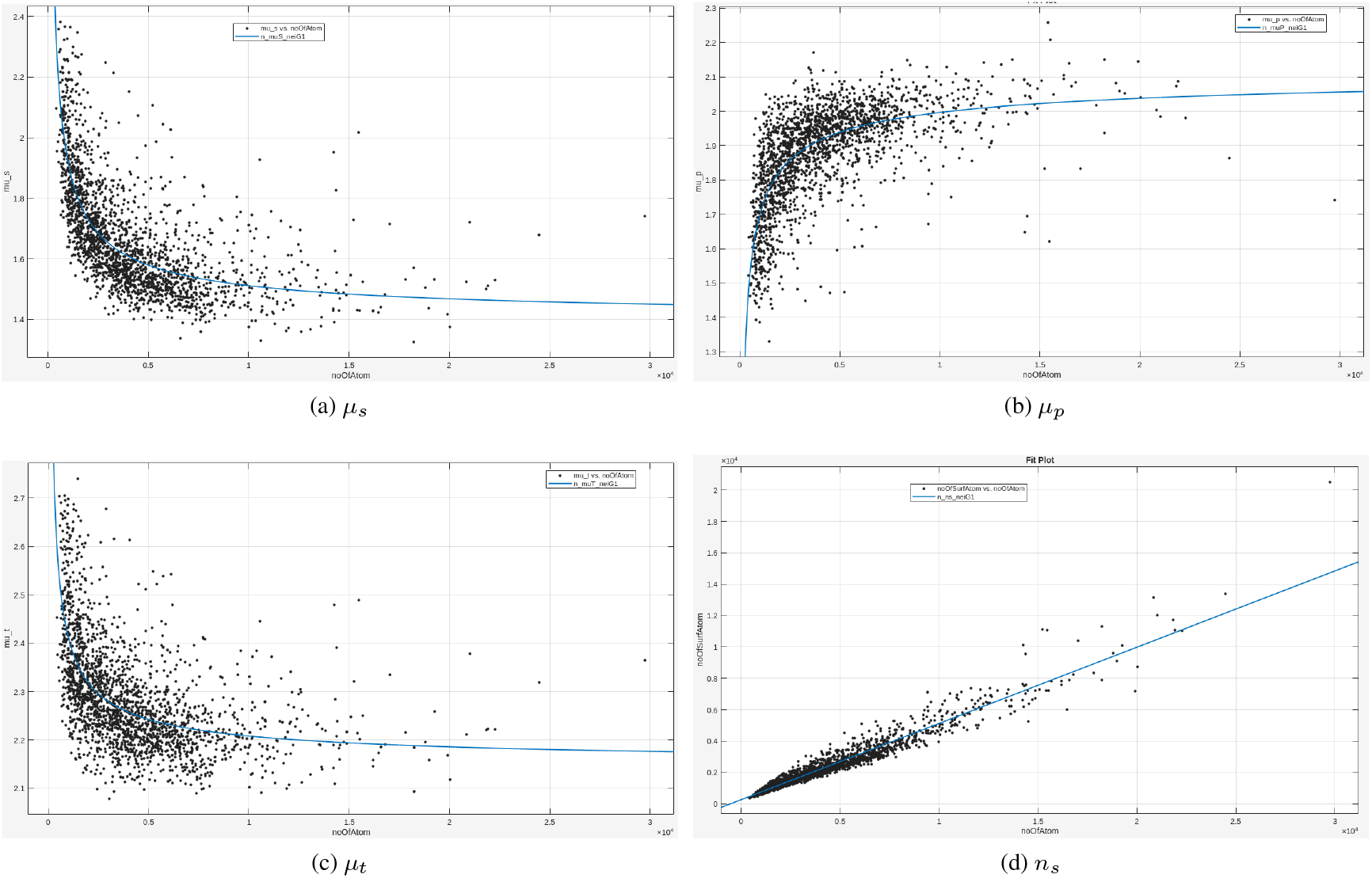
The changes with *n* of the *µ*_*s*_, *µ*_*p*_, *µ*_*t*_**s and** *n*_*s*_**s for the *Neisseria gonorrhoeae* proteome**.

**Figure S17:**
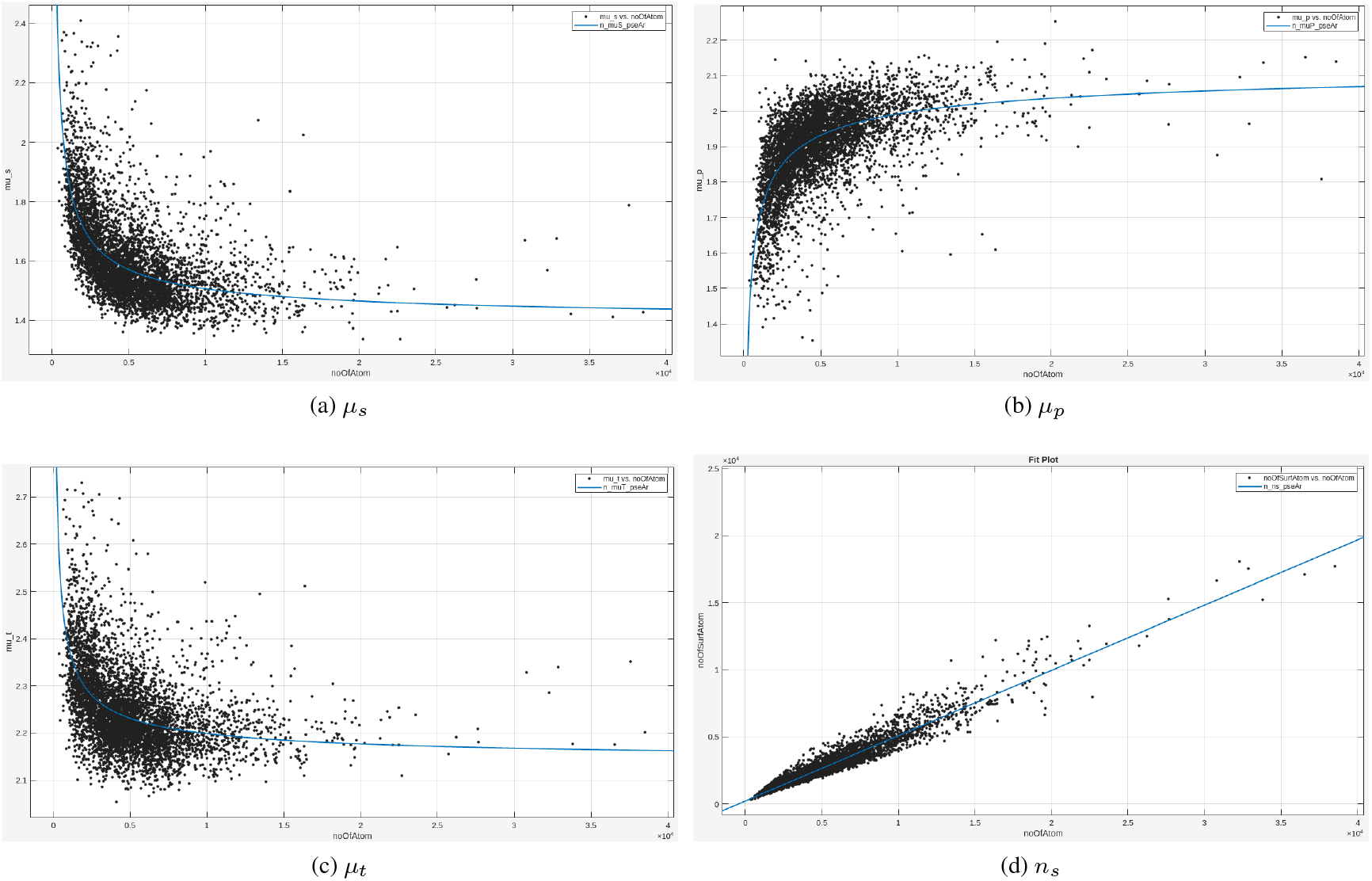
The changes with *n* of the *µ*_*s*_, *µ*_*p*_, *µ*_*t*_**s and** *n*_*s*_**s for the *Pseudomonas aeruginosa* proteome**.

**Figure S18:**
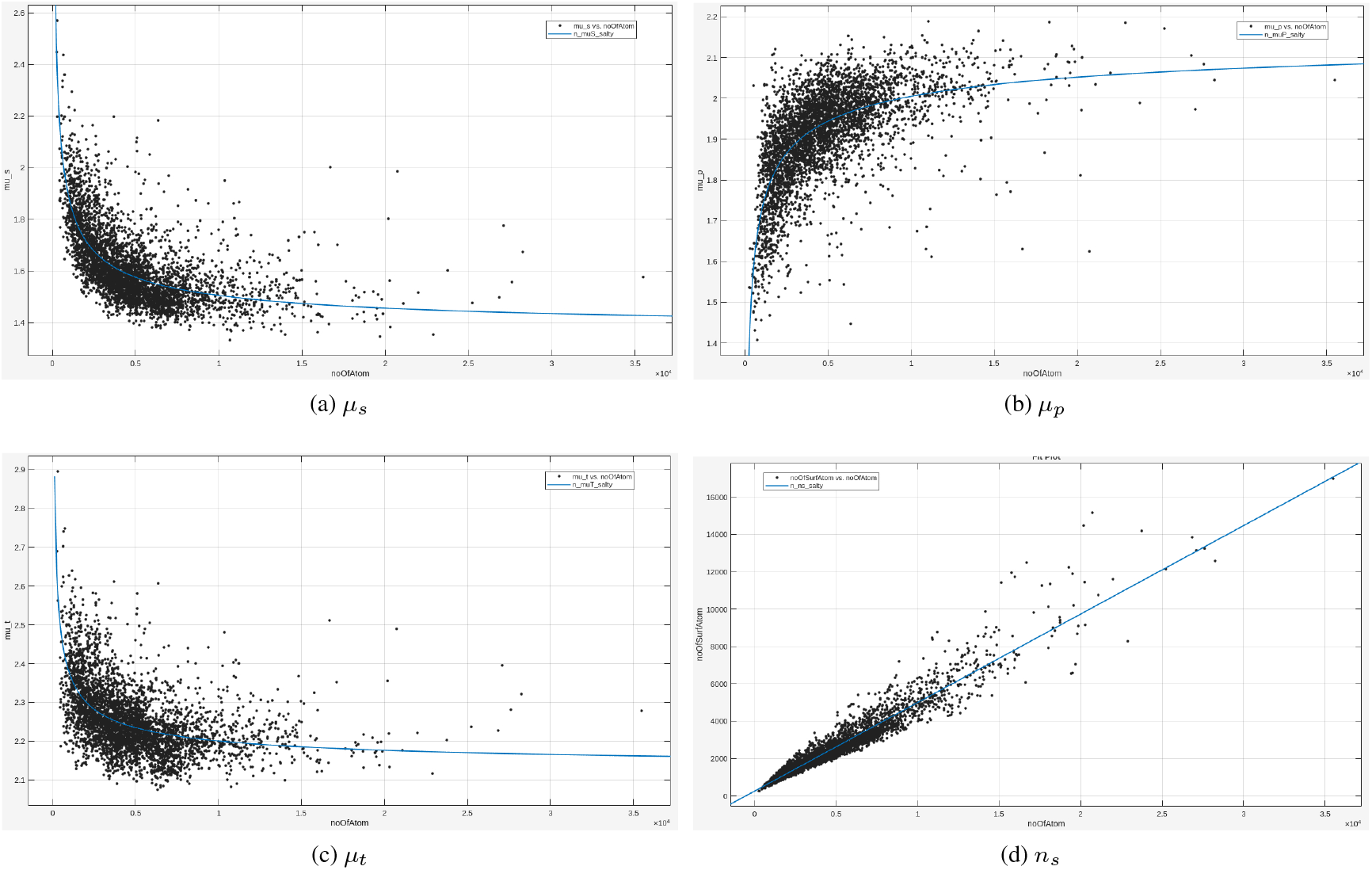
The changes with *n* of the *µ*_*s*_, *µ*_*p*_, *µ*_*t*_**s and** *n*_*s*_**s for the *Salmonella typhimurium* proteome**.

**Figure S19:**
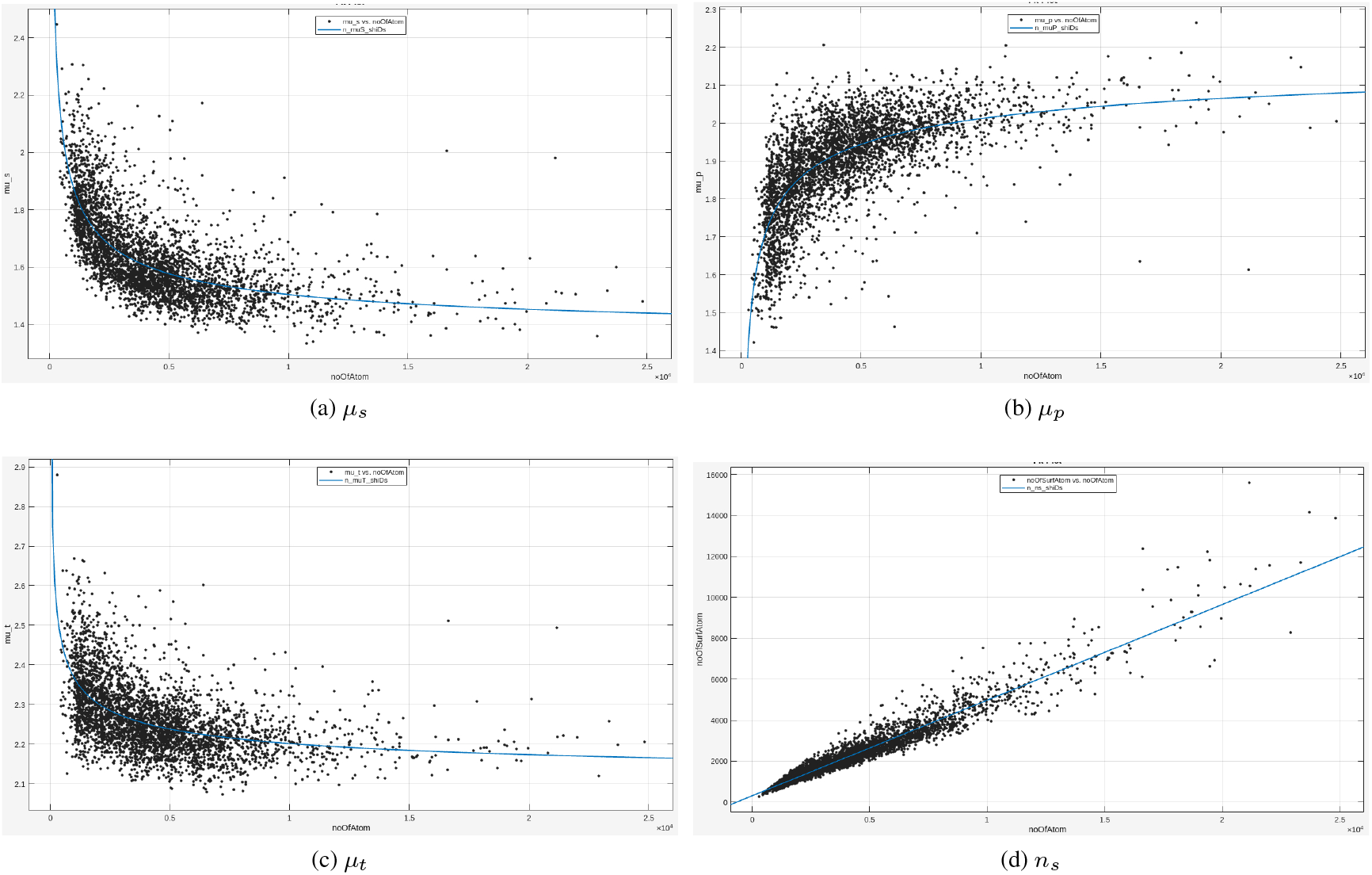
The changes with *n* of the *µ*_*s*_, *µ*_*p*_, *µ*_*t*_**s and** *n*_*s*_**s for the *Shigella dysenteriae* proteome**.

**Figure S20:**
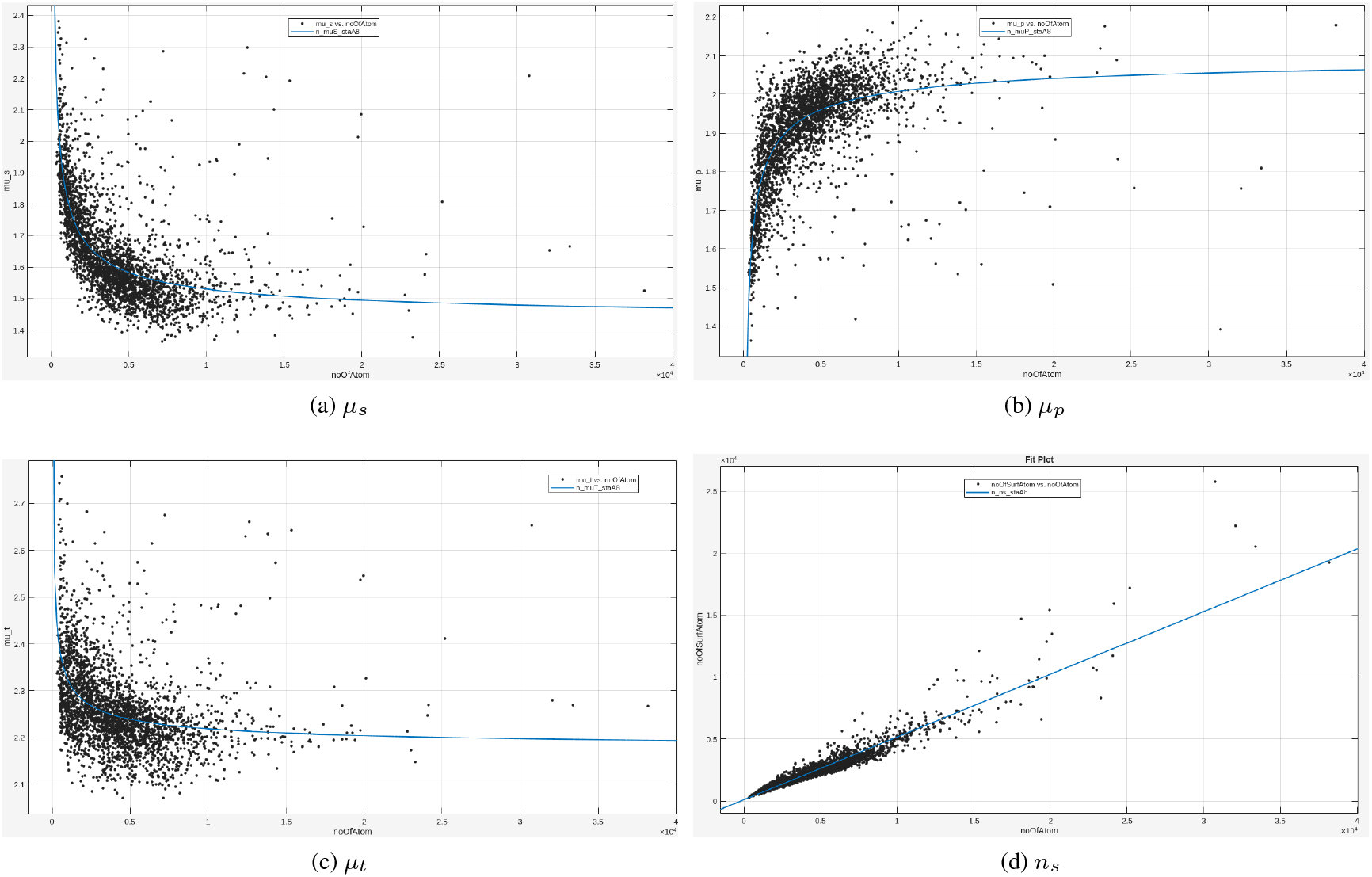
The changes with *n* of the *µ*_*s*_, *µ*_*p*_, *µ*_*t*_**s and** *n*_*s*_**s for the *Staphylococcus aureus* proteome**.

**Figure S21:**
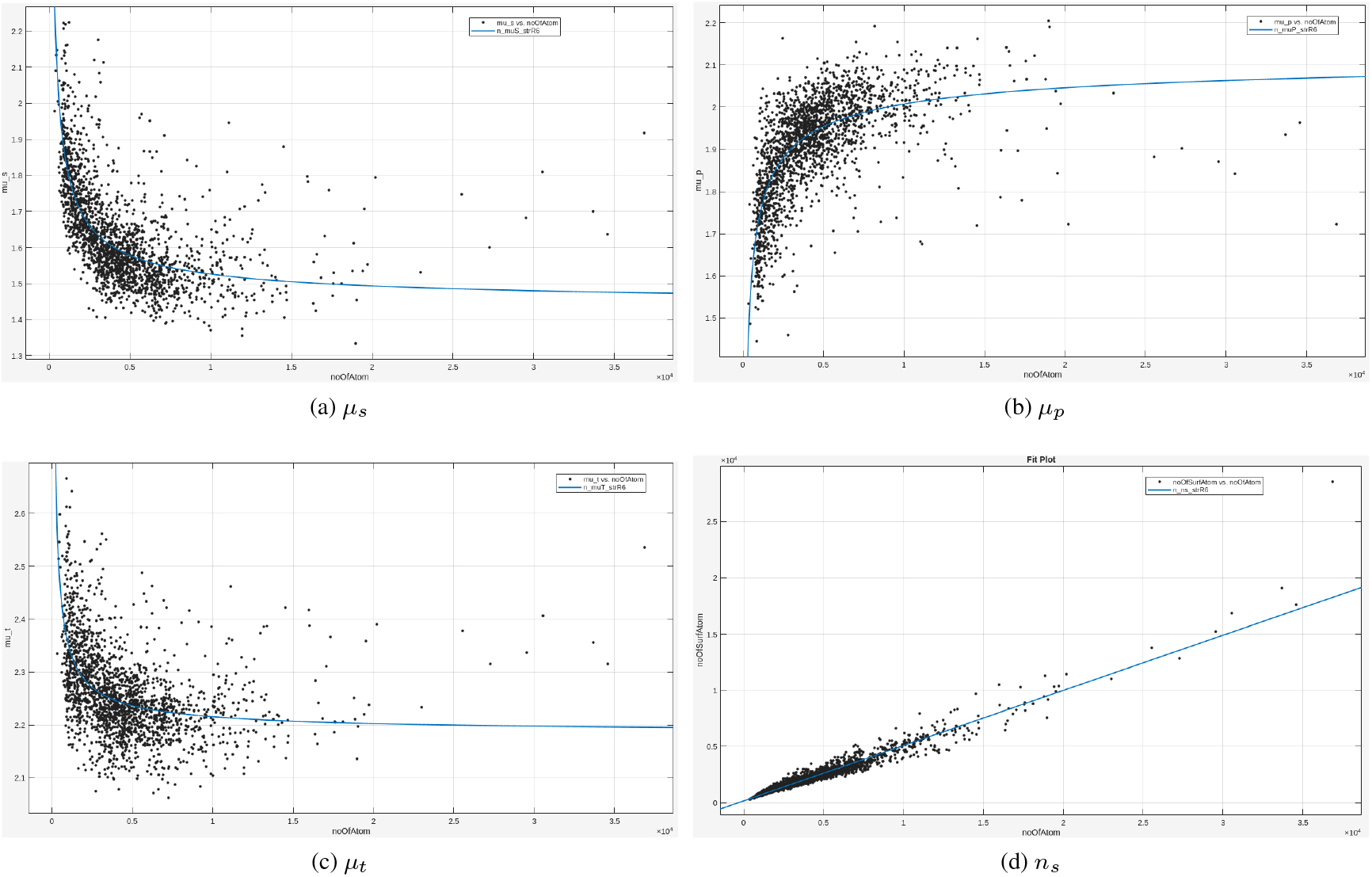
The changes with *n* of the *µ*_*s*_, *µ*_*p*_, *µ*_*t*_**s and** *n*_*s*_**s for the *Streptococcus pneumoniae* proteome**.

#### S11.2: The fourteen eukaryotic proteomes

##### S11.2.1: Three yeast proteomes

**Figure S22:**
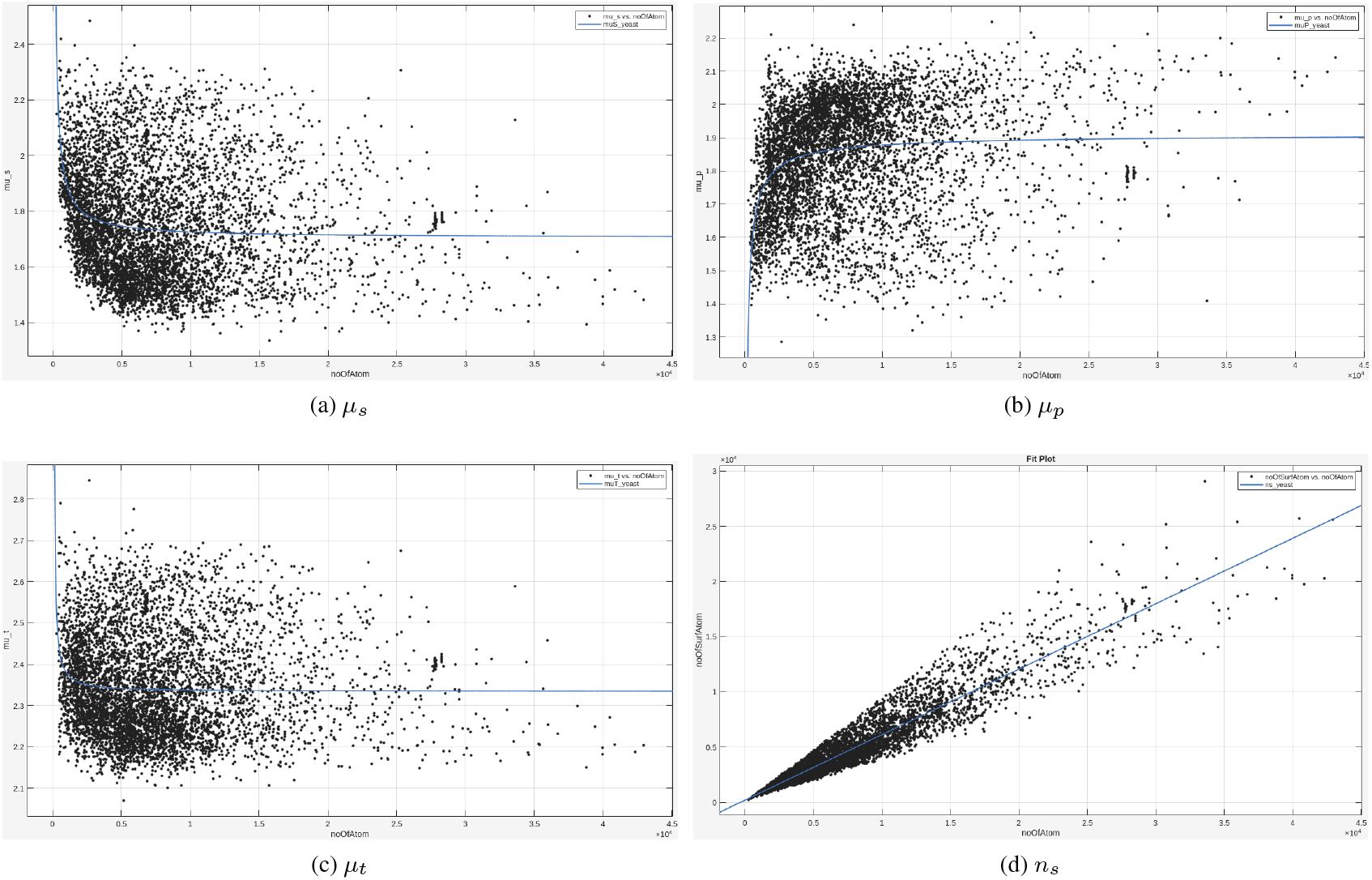
The changes with *n* of the *µ*_*s*_, *µ*_*p*_, *µ*_*t*_**s and** *n*_*s*_**s for the *Saccharomyces cerevisiae* proteome**.

**Figure S23:**
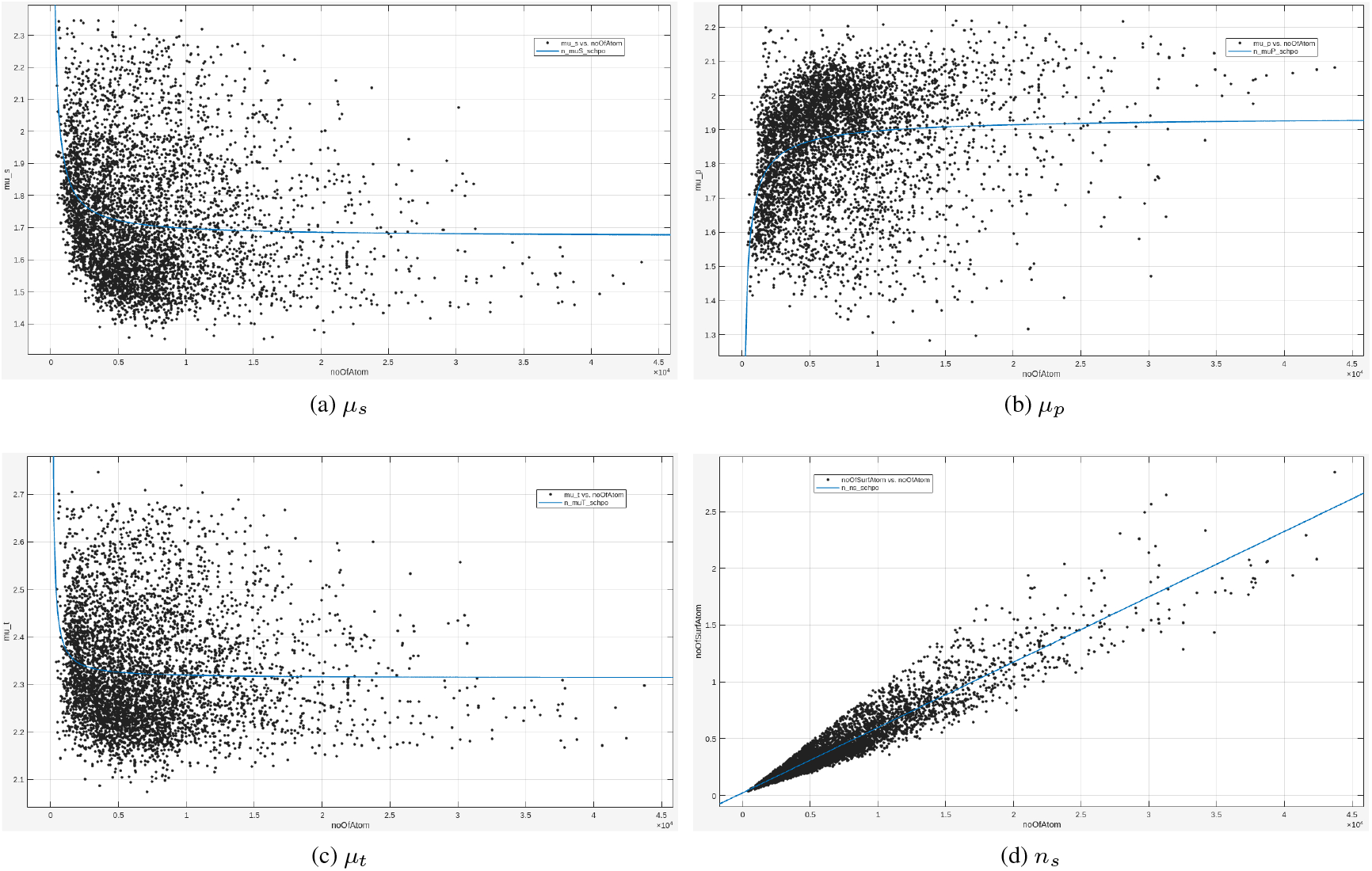
The changes with *n* of the *µ*_*s*_, *µ*_*p*_, *µ*_*t*_**s and** *n*_*s*_**s for the *Schizosaccharomyces pombe* proteome**.

**Figure S24:**
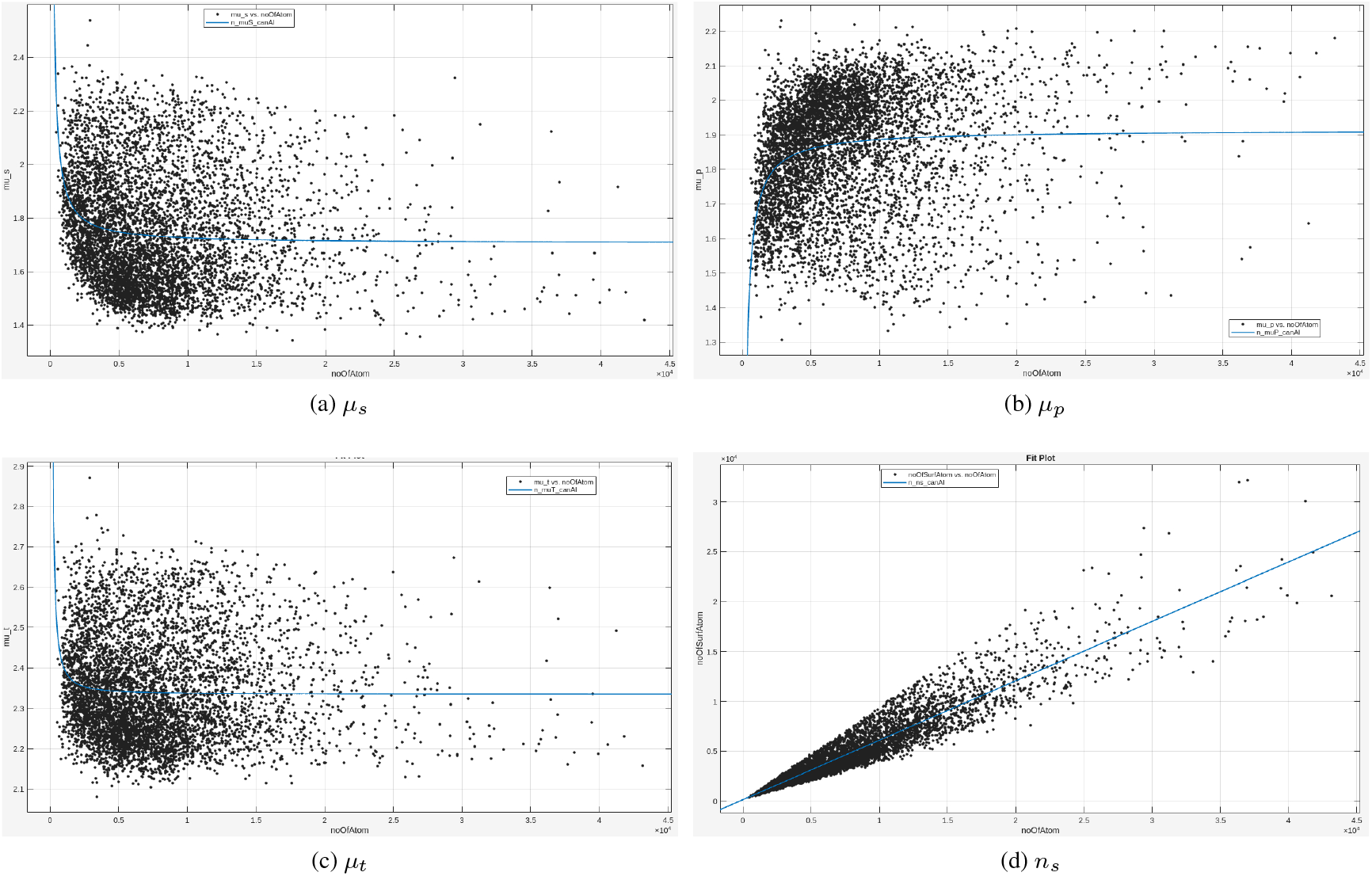
The changes with *n* of the *µ*_*s*_, *µ*_*p*_, *µ*_*t*_**s and** *n*_*s*_**s for the *Candida albicans* proteome**.

##### S11.2.2: Four plant proteomes

**Figure S25:**
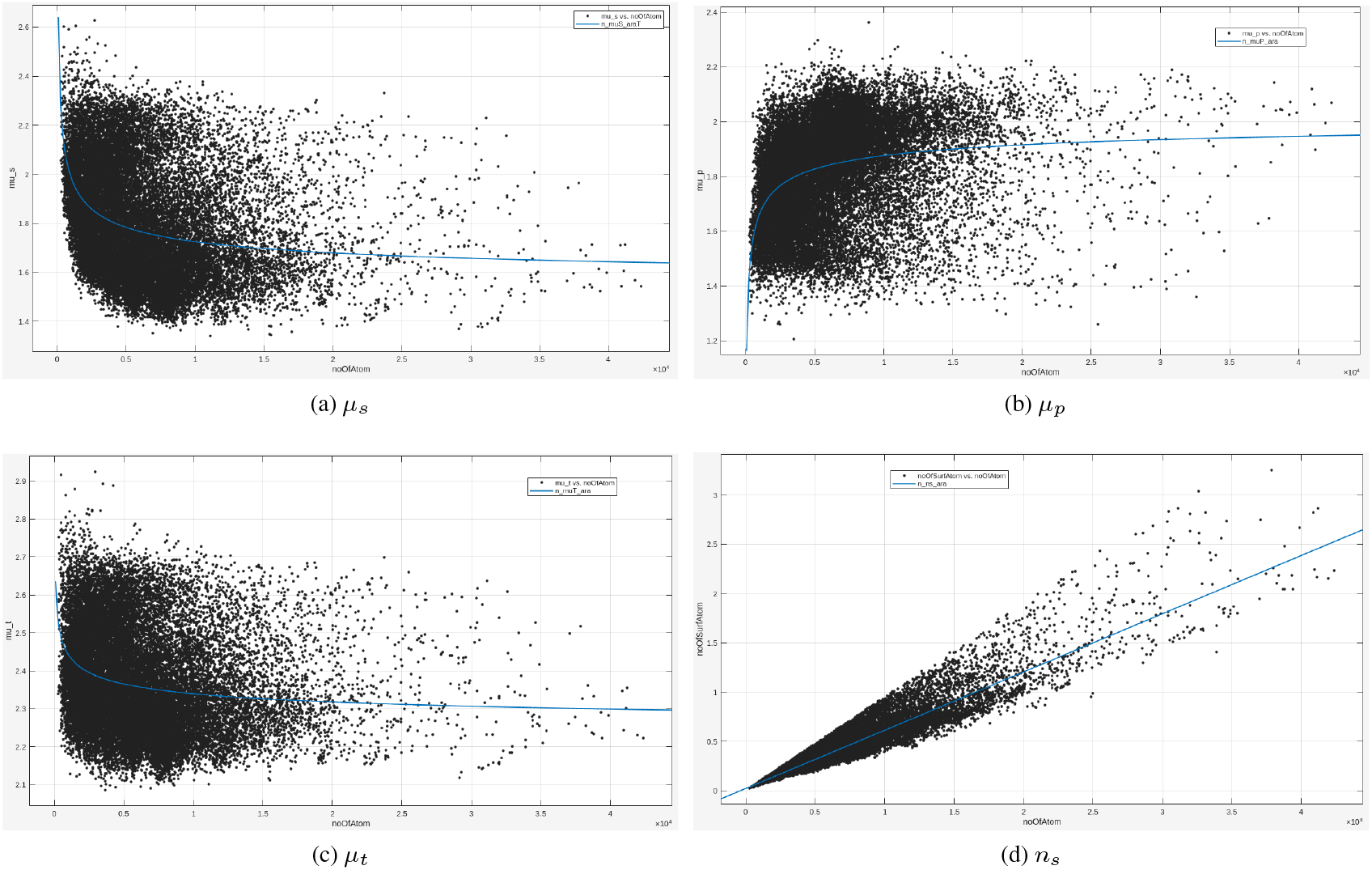
The changes with *n* of the *µ*_*s*_, *µ*_*p*_, *µ*_*t*_**s and** *n*_*s*_**s for the *Arabidopsis thaliana* proteome**.

**Figure S26:**
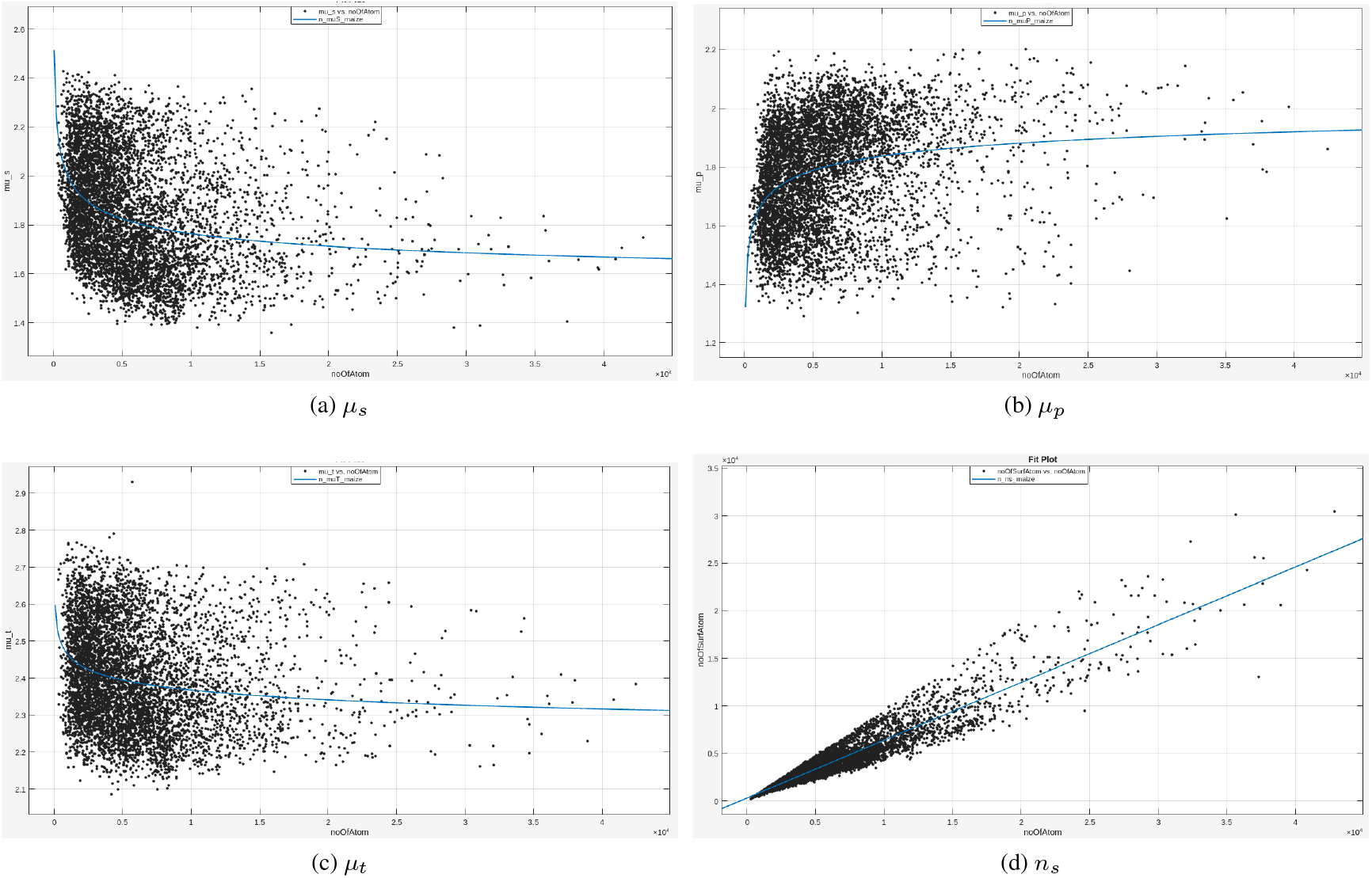
The changes with *n* of the *µ*_*s*_, *µ*_*p*_, *µ*_*t*_**s and** *n*_*s*_**s for the *Zea mays* proteome**.

**Figure S27:**
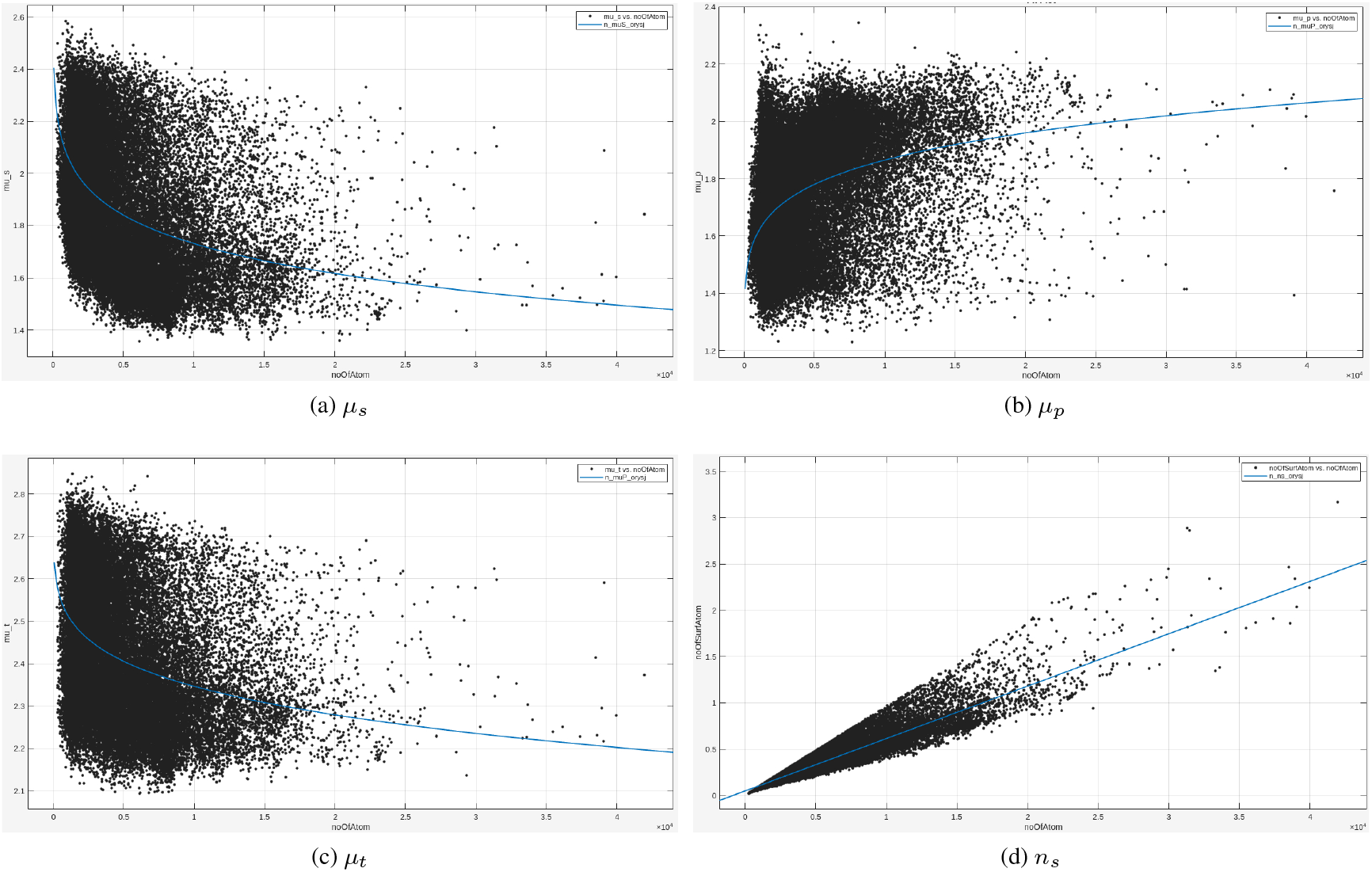
The changes with *n* of the *µ*_*s*_, *µ*_*p*_, *µ*_*t*_**s and** *n*_*s*_**s for the *Oryza sativa* proteome**.

**Figure S28:**
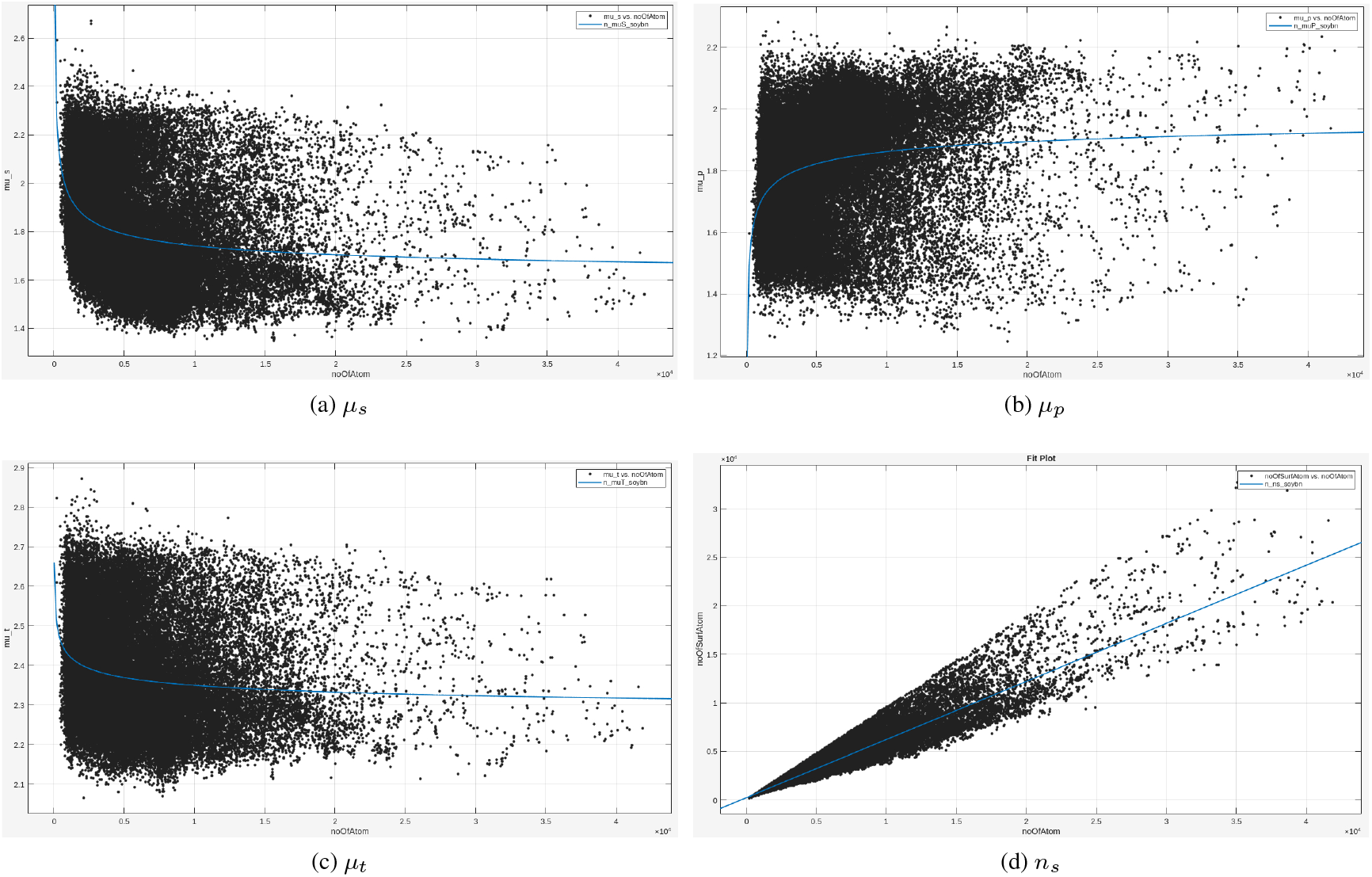
The changes with *n* of the *µ*_*s*_, *µ*_*p*_, *µ*_*t*_**s and** *n*_*s*_**s for the *Glycine max* proteome**.

##### S11.2.3: Seven animal proteomes

**Figure S29:**
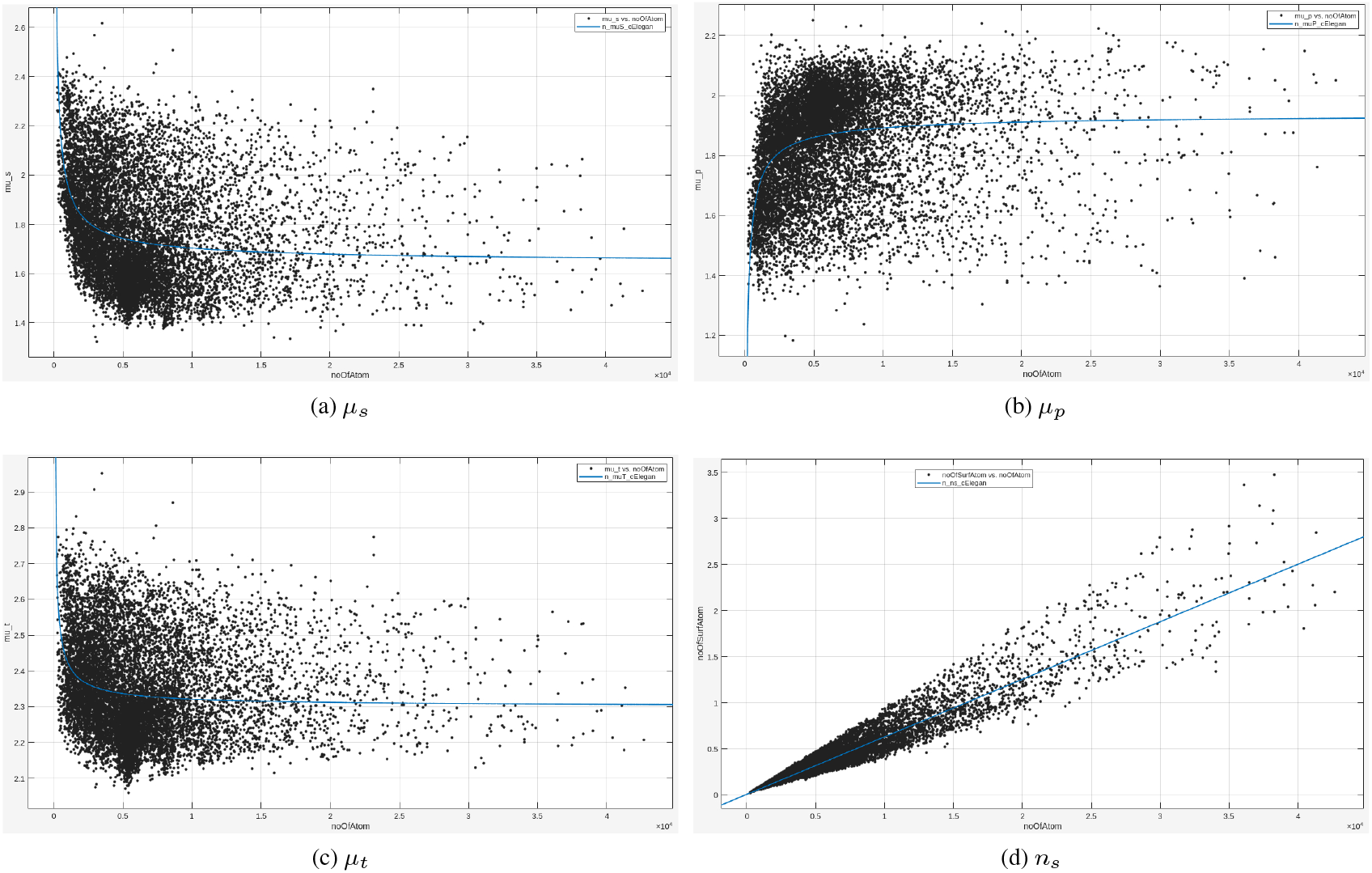
The changes with *n* of the *µ*_*s*_, *µ*_*p*_, *µ*_*t*_**s and** *n*_*s*_**s for the *Caenorhabditis elegans* proteome**.

**Figure S30:**
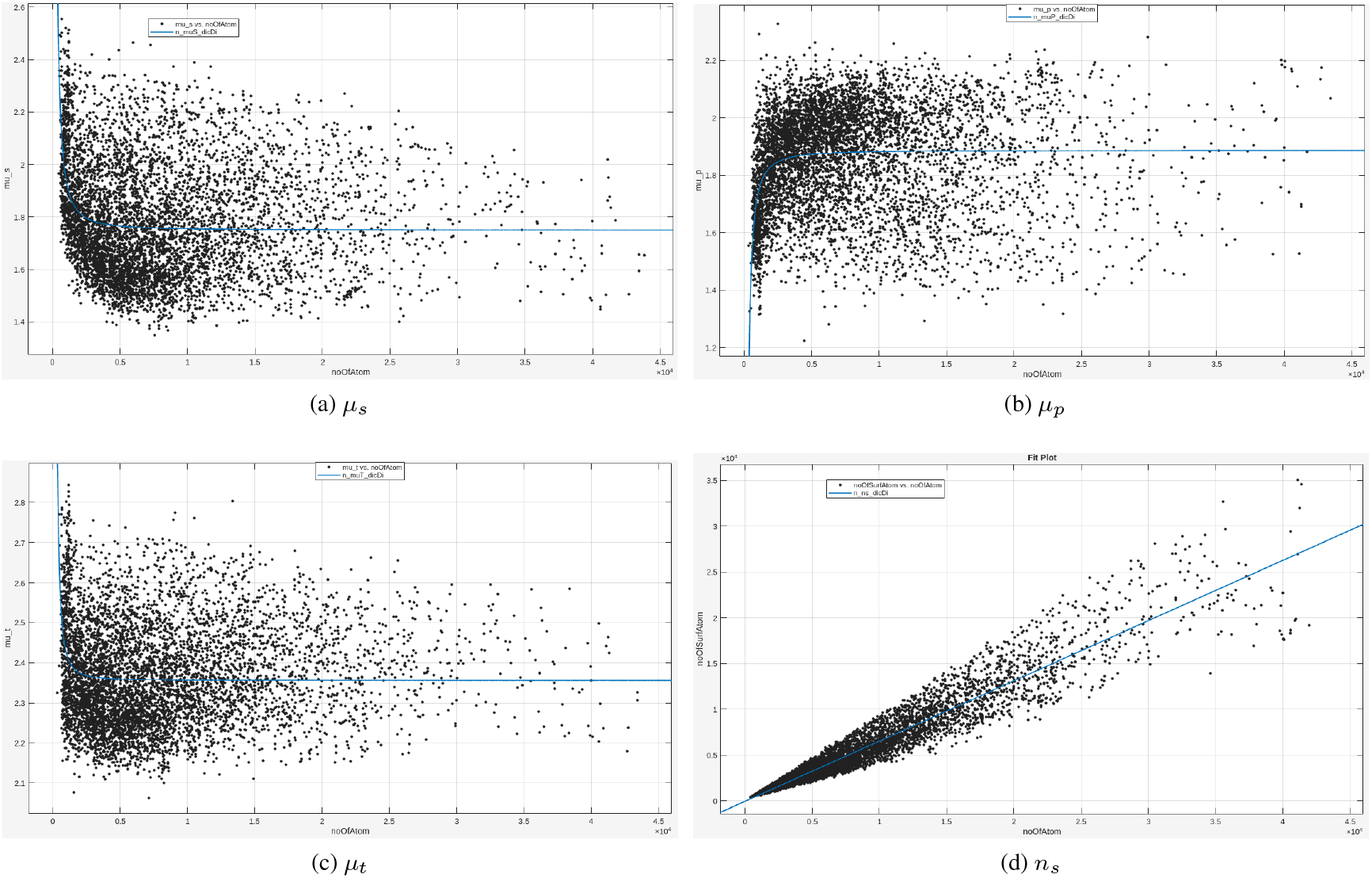
The changes with *n* of the *µ*_*s*_, *µ*_*p*_, *µ*_*t*_**s and** *n*_*s*_**s for the *Dictyostelium discoideum* proteome**.

**Figure S31:**
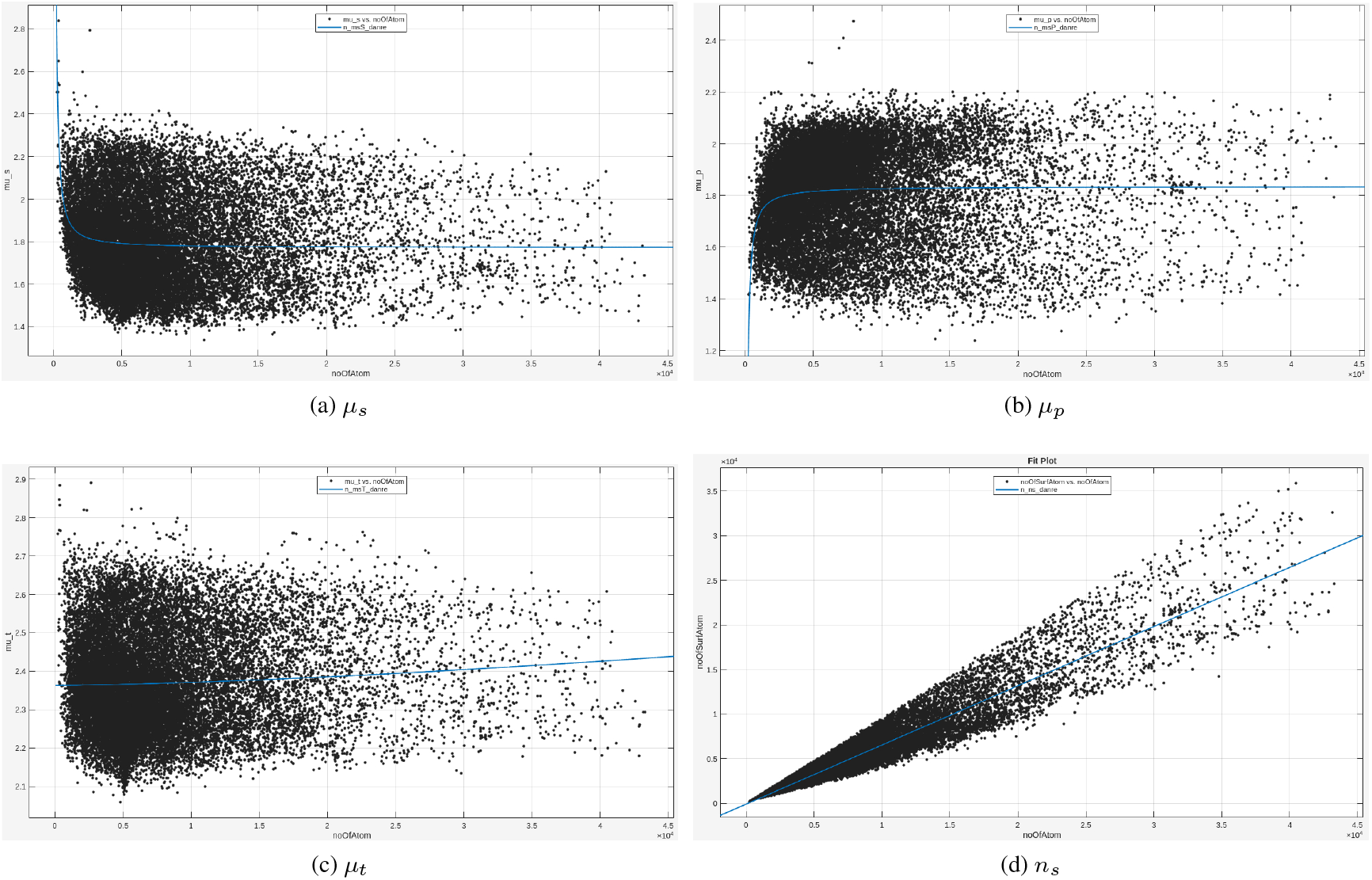
The changes with *n* of the *µ*_*s*_, *µ*_*p*_, *µ*_*t*_**s and** *n*_*s*_**s for the *Danio rerio* proteome**.

**Figure S32:**
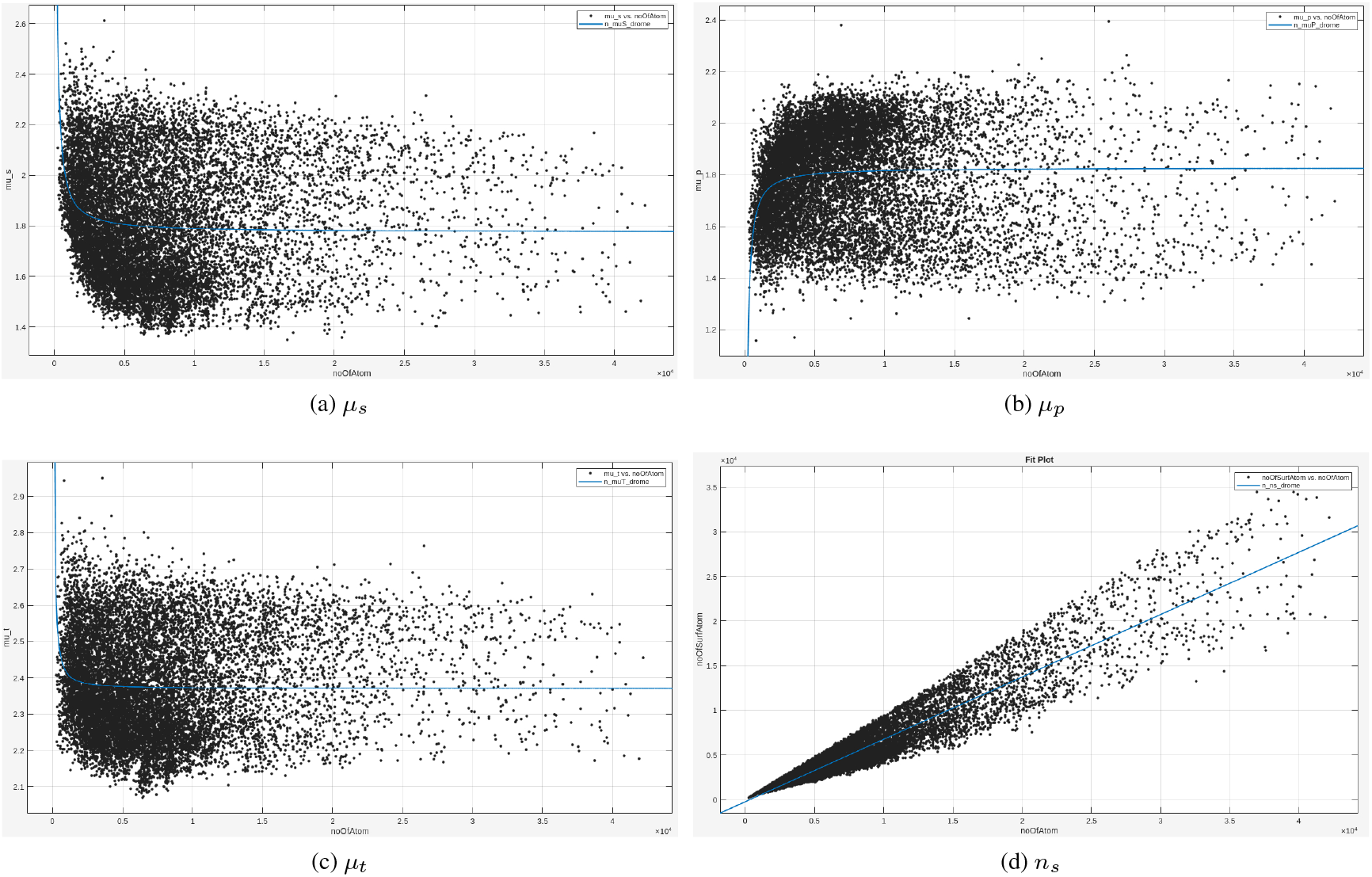
The changes with *n* of the *µ*_*s*_, *µ*_*p*_, *µ*_*t*_**s and** *n*_*s*_**s for the *Drosophila melanogaster* proteome**.

**Figure S33:**
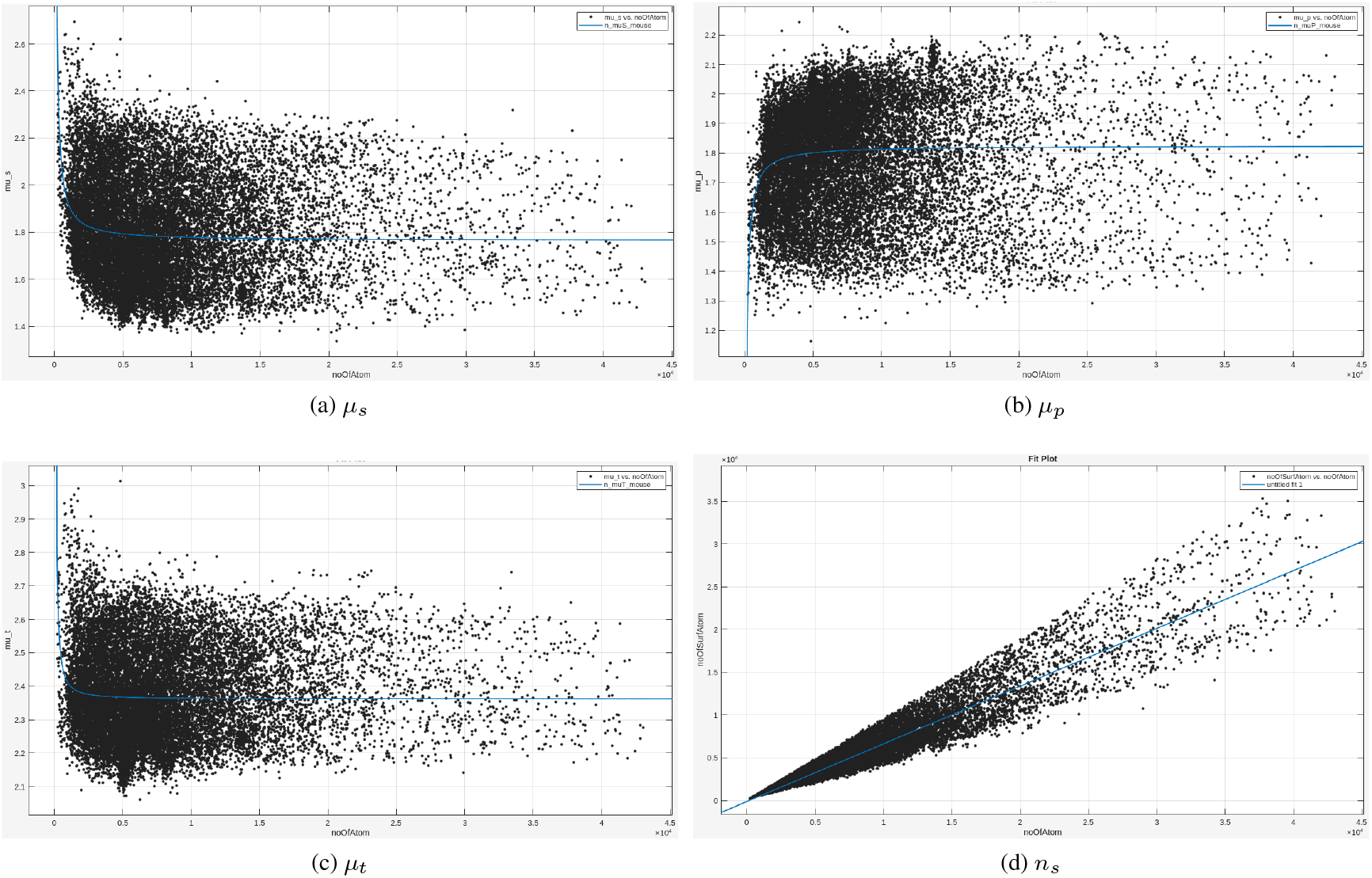
The changes with *n* of the *µ*_*s*_, *µ*_*p*_, *µ*_*t*_**s and** *n*_*s*_**s for the *Mus musculus* proteome**.

**Figure S34:**
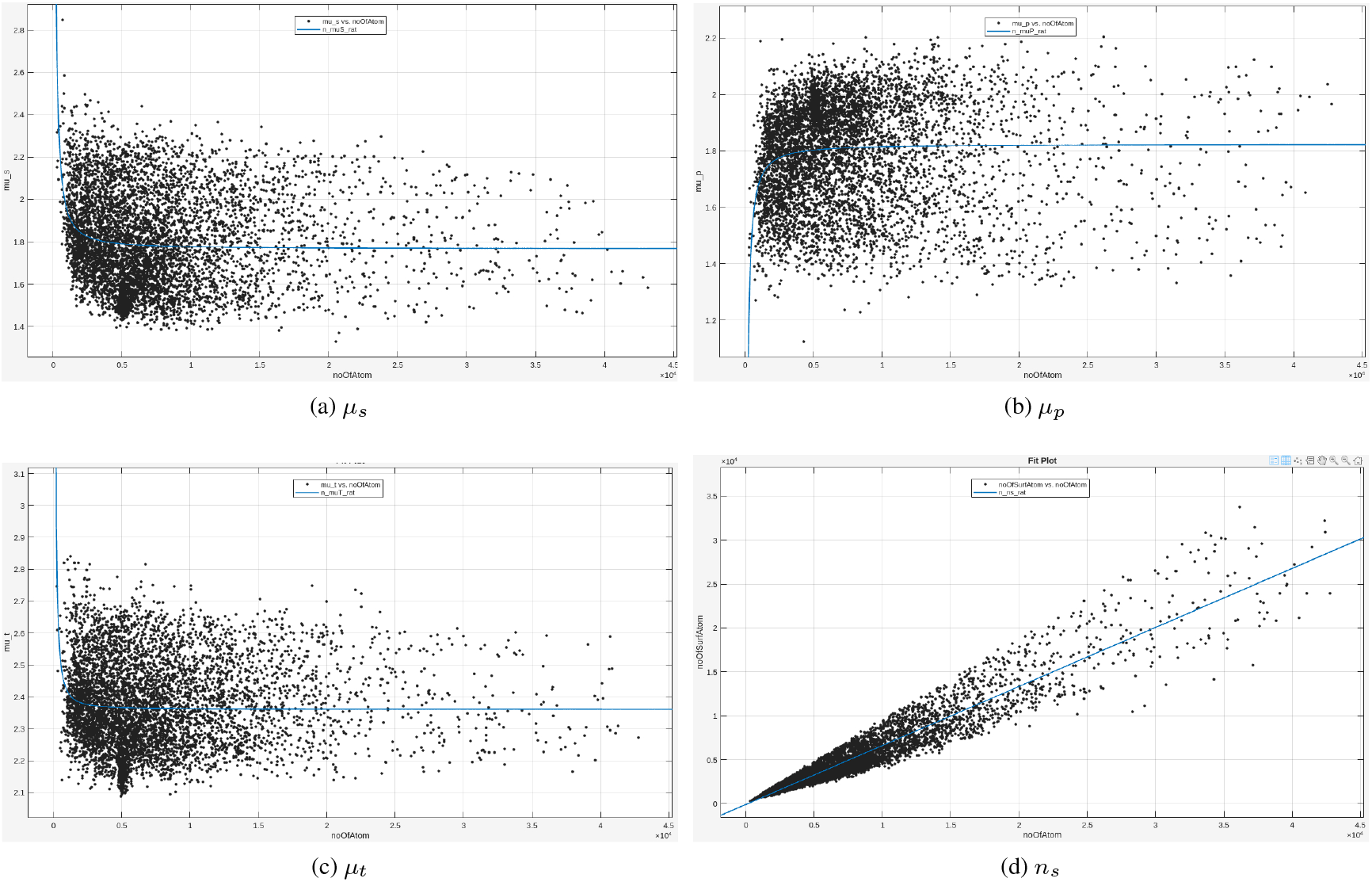
The changes with *n* of the *µ*_*s*_, *µ*_*p*_, *µ*_*t*_**s and** *n*_*s*_**s for the *Rattus norvegicus* proteome**.

**Figure S35:**
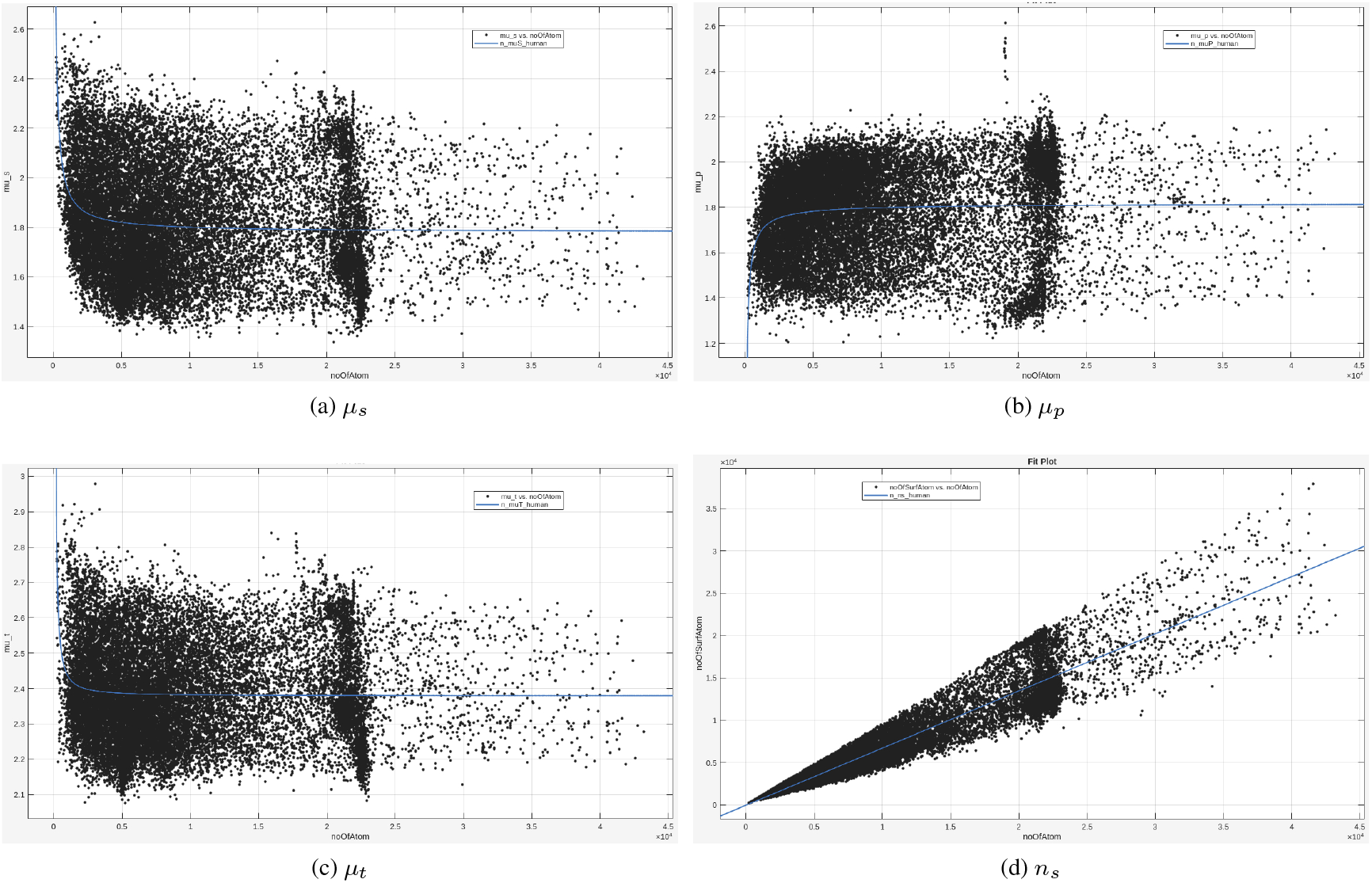
The changes with *n* of the *µ*_*s*_, *µ*_*p*_, *µ*_*t*_**s and** *n*_*s*_**s for the *Homo sapiens* proteome**.

### S12: The biological names of the twenty seven species and their abbreviations

Table S3 lists the biological names of the twenty seven species and their abbreviations used in Table 2 of the main text and Tables S1 and S2, and Fig. S5.

**Table S3:** The name of the twenty seven species and their abbreviations used in Table 2 of the main text and Tables S1 and S2 and Fig. S5.

| Species | Abbreviation |
| --- | --- |
| *Escherichia coli* | ecoli |
| *Haemophilus influenzae* | haeIn |
| *Helicobacter pylori* | helPy |
| *Klebsiella pneumoniae* | klePh |
| *Methanocaldococcus jannaschii* | metJa |
| *Mycobacterium leprae* | mycLe |
| *Mycobacterium tuberculosis* | mycTu |
| *Neisseria gonorrhoeae* | neiG1 |
| *Pseudomonas aeruginosa* | pseAr |
| *Salmonella typhimurium* | salTy |
| *Shigella dysenteriae* | shiDs |
| *Staphylococcus aureus* | staA8 |
| *Streptococcus pneumoniae* | strR6 |
| *Saccharomyces cerevisiae* | yeast |
| *Schizosaccharomyces pombe* | schPo |
| *Candida albicans* | canals |
| *Arabidopsis thaliana* | araTh |
| *Zea mays* | maize |
| *Oryza sativa* | orySj |
| *Glycine max* | soyBn |
| *Caenorhabditis elegans* | celEg |
| *Dictyostelium discoideum* | dicDi |
| *Danio rerio* | danRe |
| *Drosophila melanogaster* | droMe |
| *Mus musculus* | mouse |
| *Rattus norvegicus* | rat |
| *Homo sapiens* | human |

### S13: An example of the interface formed by crystal packing

**Figure S36:**
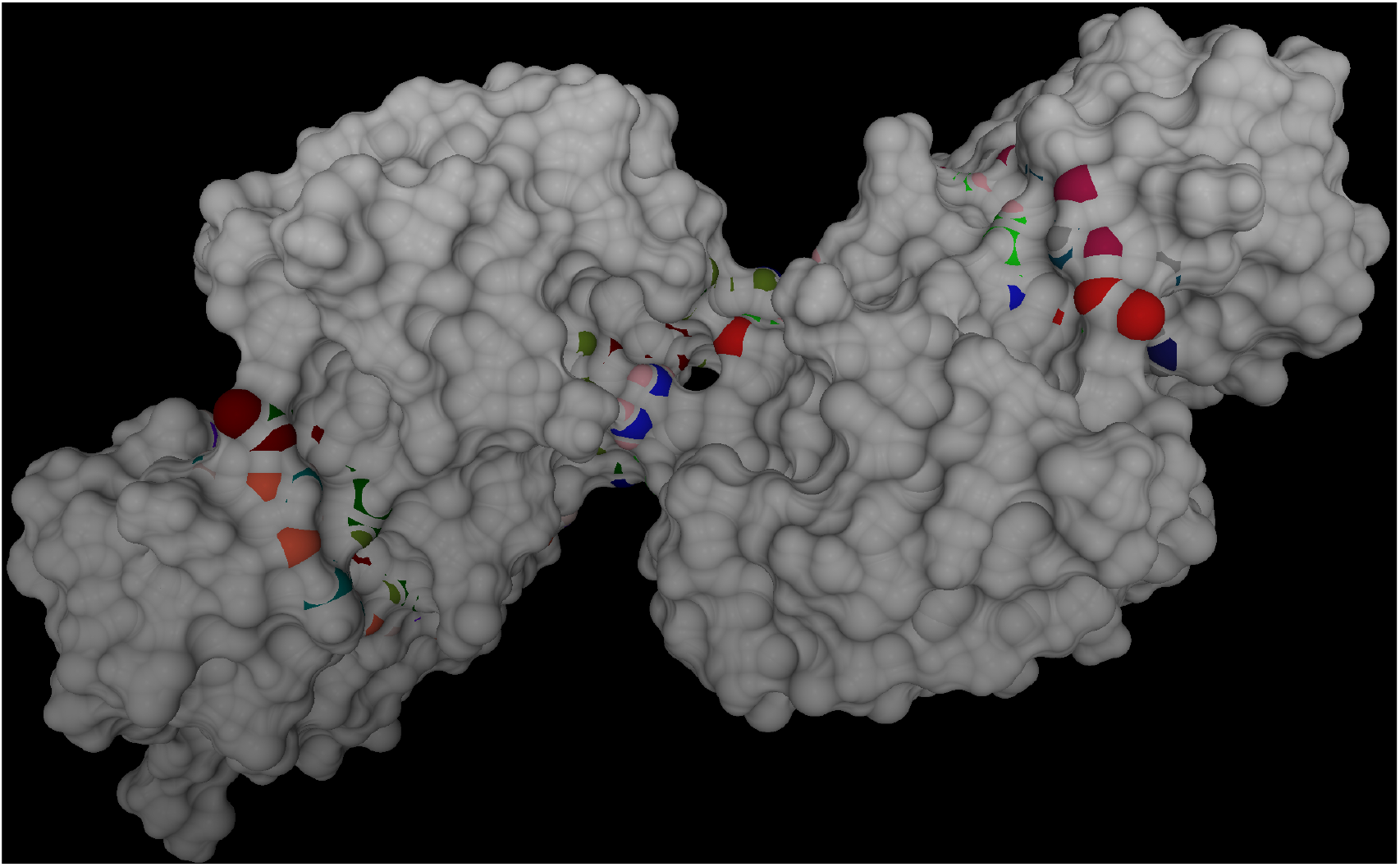
The crystal packing between barnase-barstar complex. The two PPI interfaces of barnase barstar complex are in the right and the left while the interface formed by crystal packing is in the middle. The SAS polygons of the interface atoms are depicted in bright colors while the rest of the SES in gray. As illustrated here, the interface of crystal packing is relatively small and thus its contribution to the geometric properties studied in this paper is minor. The figure is prepared by our program SESA which is freely available at https://github.com/wlincong/SESexact.

### S14: The pdbids of the three sets of crystal structures

The file “pdbidOfGSM.tar.gz” includes three files that list respectively the pdbids for **G, S** and **M**.

## Footnotes

1 Abbreviations: 2D, two-dimensional; 3D, three-dimensional; SES, solvent-excluded surface; SAS, solvent-accessible surface; SASA, solvent-accessible surface area; RMSD, root-mean square deviation; RMSE: root-mean squared error; PDB, Protein Data Bank; TM-score, template modelling score; GDT, global distance test; pLDDT, predicted local distance difference test; SI, Supporting Information.

2 In this paper, SES and molecular surface are used interchangeably.

3 An internal residue means a residue between the N-terminal and C-terminal residues.

4 For brevity, the symbol”*µ*s” (SES patch areas per atom) means *µ*_*s*_s, *µ*_*p*_s and *µ*_*t*_s.

5 The number of missing atoms for a crystal structure is computed as the difference between the number of atoms detected and that expected from the protein sequence.

6 SESA is freely available at www.github/wlincong/sesA.

7 The *µ*_*s*_ range here means the range of the *µ*_*s*_s with approximately the same *n*.

8 The 53.2% difference is calculated as follows: 53.2 = 57.7 −4.5 where 57.7 is the *P*_*s*_ value for the human proteome (Table S2). It means that 53.2% of the *µ*_*s*_s for the same *n* for the human proteome are out of the *µ*_*s*_ range for the *E*.*coli* proteome.

